# Accurate, comprehensive gene annotation and ortholog identification across thousands of vertebrate genomes with TOGA2

**DOI:** 10.64898/2026.06.30.735536

**Authors:** Yury V. Malovichko, Bernhard Bein, Alejandro Gonzales-Irribarren, Evgeny Leushkin, Leon Hilgers, Amy Stephen, Xueling Yi, Michele Albertini, Tim Stadager, Markus Zumpt, Luca Hoppach, Felix Götz, Niklas Himstedt, Lucas Koch, The Vertebrate Genomes Project Consortium Phase I, Michael Hiller

## Abstract

Inferring orthologs and annotating coding genes remain central challenges in genomics, evident by the growing gap between assembled and annotated genomes. TOGA (Tool to infer Orthologs from Genome Alignments) addresses this challenge by integrating gene annotation and orthology inference. Here, we present TOGA2, the next generation of TOGA, which substantially improves annotation completeness, accuracy, scalability, and orthology inference. TOGA2 leverages exon-level orthology and introduces an exon-wise annotation procedure that reduces memory usage 513-fold and runtime 6.1-fold. We show that human-trained deep learning models for splice site prediction generalize across vertebrates. Integrating these predictions enables robust handling of evolutionary changes in exon-intron structure, including splice site shifts, intron deletions, and exonization of introns. A new gene tree reconciliation step refines orthology inference, and UTR annotation improves gene model completeness. Across mammals, birds, turtles, and percomorph fishes, TOGA2 annotations generally achieve higher gene completeness than transcriptome-informed RefSeq annotations. TOGA2 identifies previously unannotated exons in mouse, assigns informative gene symbols, and annotates V(D)J segments of antigen receptors. TOGA2 scales to thousands of genomes, which we demonstrate by generating comprehensive comparative genomics resources for 2,162 vertebrate assemblies, including gene annotations, ortholog sets, gene losses and duplications, retrogene candidates, and outputs supporting downstream analyses. Together, TOGA2 provides a scalable and versatile framework for comparative genomics that bridges the genome annotation gap.

## Introduction

Comparative genomics enables a wide range of applications across biomedical, functional, and evolutionary research. This includes leveraging model organisms to elucidate human disease biology, identifying likely pathogenic variants, quantifying gene essentiality, and transferring functional annotations from model organisms to related species ^1–6^. It also enables the inference of species phylogenies, the reconstruction of ancestral genomes, and the inference of chromosome evolution and rearrangement history ^7–9^. In addition, comparative genomics provides the foundation for the characterization of changes in the gene repertoire, the identification of molecular adaptations, linking phenotypic changes to underlying genetic differences, and cross-species comparisons of gene expression ^10–16^.

Fundamental to these analyses is the identification of orthologous genes that originated from speciation events, and distinguishing them from paralogous genes that arose from duplication events. Orthology inference methods rely on pre-identified exon-intron structures and coding regions in the genomes (referred to as structural gene annotation) to infer orthologous gene relationships, either by clustering based on sequence similarity or by gene tree reconstruction ^17–25^. The reliance on pre-existing gene annotations represents a key limitation, as gene annotation itself is challenging ^26,27^. Despite rapid progress in deep learning–based *ab initio* gene prediction ^28–30^, accurately annotating rare genomic features and complex gene structures remains challenging ^31^, and transcriptome-based approaches are limited by the requirement for comprehensive transcriptome data ^32,33^. Furthermore, variation in annotation quality and completeness can substantially affect orthology inference ^34^. Together with the rapidly growing generation of high-quality genome assemblies ^35–37^, this has led to a situation in which the number of available genomes far exceeds those that are accurately annotated and amenable to orthology inference.

A method that addresses these gaps is TOGA1 (Tool to infer Orthologs from Genome Alignments), which explicitly integrates gene annotation and orthology inference in a single framework ^38^. TOGA1 leverages a whole-genome alignment between an annotated reference genome and an unannotated query genome, represented by chains of co-linear local alignments that capture orthologous, paralogous, and processed pseudogene loci ^39^. Unlike other methods that rely on protein-coding sequences ^17–25^, TOGA1 primarily relies on intronic and intergenic alignments and employs machine learning to accurately distinguish orthologous loci from paralogous loci and processed pseudogenes (intronless gene copies generated by retrotransposition). TOGA1 uses the reference annotation to annotate genes in orthologous loci in the query genome. It classifies each annotated transcript based on its open reading frame integrity and infers orthology relationships between reference and query (1:1, 1:many, many:1, many:many), thereby enabling the identification of gene loss and duplication events. TOGA1’s reference-based methodology scales linearly with the number of unannotated query genomes, making it well-suited for generating annotations and orthology data for hundreds of species. TOGA1 has been shown to be a top-performing annotation method in comparative benchmarks ^27^ and has been applied in multiple studies to generate genome-wide annotations and orthologs ^40–43^. Furthermore, it has been used as a powerful basis to identify evolutionary gene changes (loss, duplication, signatures of selection) and link them with phenotypic differences ^44–57^.

Despite TOGA1’s success, the method had several limitations, including substantial runtime and memory requirements for large genes, limited ability to capture evolutionary exon–intron structure changes, less comprehensive input annotations, occasional failure to identify correct orthology relationships, and the inability to annotate UTRs. Here, we present TOGA2, a new implementation that improves all aspects of the original TOGA1 method ^38^. We show that TOGA2 generally outperforms RefSeq annotations in terms of completeness, identifies previously unannotated exons in mouse, provides informative gene symbols, accurately identifies V(D)J segments in immune genes, and supports several applications downstream of annotation and orthology inference. By applying TOGA2 to thousands of vertebrate genomes, including high-quality assemblies generated as part of the Vertebrate Genomes Phase I ^37^, we generated the largest comparative genomics resource of vertebrates to date, spanning 2,162 genomes across mammals, birds, turtles, and percomorph fish.

## Results

### Leveraging orthology at the exon level

The first major advance in TOGA2 is the replacement of the transcript-wise annotation strategy of TOGA1 with a substantially more runtime- and memory-efficient exon-wise approach. TOGA1 considers the entire orthologous locus of a transcript and uses the splice site-aware coding exon aligner CESAR2 ^58^ to infer the positions and boundaries of all exons simultaneously. While this enables the detection of diverged exons that align only at the protein level and the identification of precise intron deletion events resulting in the fusion of adjacent query exons ^58,59^, it is computationally expensive because the complete transcript locus, including all introns, must be considered. In contrast, TOGA2 leverages that the positions of the vast majority of exons in the query genome can be accurately inferred from the genome alignment chain (Fig. 1A). Using this exon-level orthology, TOGA2 replaces the transcript-wise strategy with an exon-wise CESAR2-based annotation approach that restricts the analysis to individual exon loci with flanking regions to allow for potential splice site shifts. To retain the ability to detect precise intron deletions and diverged exons that align only at the protein level, TOGA2 pre-identifies such cases and performs joint analyses of the corresponding query loci spanning the affected exons. In addition, TOGA2 annotates terminal exons not covered by the alignment chain or located beyond chain boundaries, while such exons are missed by TOGA1 (Supplementary Figure 1).

**Figure 1:**
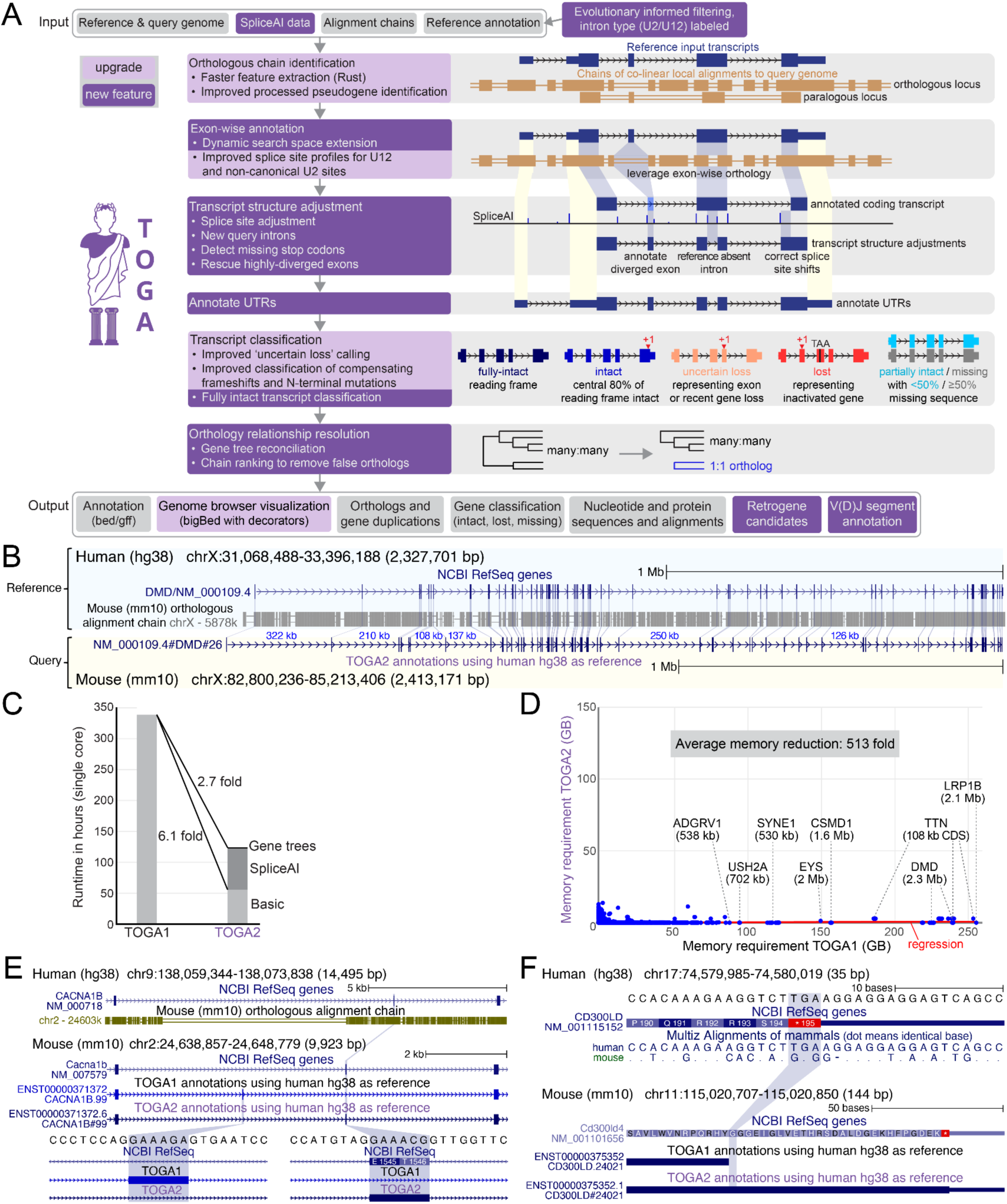
Overview of TOGA2. (A) Illustration of new and improved features in TOGA2, as well as its input and output. (B) Exon-level orthology illustrated for *DMD*, a gene with several large introns. Whereas TOGA1 considers the entire 2.4 Mb orthologous gene locus to identify and annotate all 79 coding exons, TOGA2 infers orthology at the exon level and aligns each reference exon independently to its orthologous locus (blue connecting lines), cumulatively restricting the search space to 26,708 bp. Introns >100 kb are labeled in blue font. (C, D) Runtime and memory efficiency, exemplified by human (reference) versus mouse (query). (C) Compared to TOGA1, exon-wise annotation and an efficient implementation substantially increase the speed of TOGA2, even when incorporating additional computational features. Runtime was determined in a sequential (single-core) setting. (D) Exon-wise annotation drastically reduces memory requirements. Individual genes with large query loci are labeled. (E) Exon-level orthology improves exon annotation accuracy, illustrated for *CACNA1B*, which contains a six bp microexon. TOGA1 searches for this microexon within a much larger query locus and misannotates it approximately 2.5 kb upstream. In contrast, TOGA2 leverages exon-level orthology, where intronic alignments flanking the exon clearly define its location, enabling its correct annotation. (F) Capturing mutated stop codons improves terminal exon annotation accuracy. In a mouse ortholog of *CD300LD*, the TGA stop codon is mutated to a GGG sense codon, resulting in a 81 bp C-terminal protein extension. In contrast to TOGA1, TOGA2 detects the mutated stop codon and scans the downstream sequence for an in-frame stop codon, resulting in the correct annotation.

We found that in a typical mammalian TOGA2 run with human as the reference species, approximately 89% of exons can be processed individually using exon-wise annotation, which drastically reduces runtime and memory requirements. This is illustrated by *DMD*, which spans 2.3 Mb in the mouse genome (Fig. 1B). While TOGA1’s transcript-wise annotation requires 240 GB of memory and 71 min, TOGA2’s exon-wise annotation procedure requires only 1.4 MB of memory and 6 sec. Even *TTN*, which produces transcripts comprising a total of 362 exons, including a 17 kb coding exon and a total of 108 kb of coding sequence, can be annotated by TOGA2 with 2.85 GB of memory in 1.7 min, whereas TOGA1 requires 253 GB and 2.8 hours. Exon-wise annotation and an optimized implementation of orthologous chain identification enable TOGA2 to reduce memory requirements by an average of 513-fold and runtime by 6.1-fold relative to TOGA1 in a genome-wide benchmark of 67,036 transcripts (Fig. 1C,D, Supplementary Table 1).

### Improved detection of short and terminal exons

Exon-wise alignment is also one of several strategies in TOGA2 that improve annotation accuracy, particularly for small internal and terminal coding exons. Correctly annotating such exons with homology-based methods is challenging, especially when they exhibit sequence divergence. However, flanking intronic alignments in genome alignment chains typically enable the precise localization of these exons in the query genome. By exploiting exon-level orthology, TOGA2 enables accurate annotation of very short exons, even when they are as short as two nucleotides (Fig. 1E, Supplementary Figure 2).

A second feature that improves annotation accuracy of terminal exons is the detection of mutations that remove the terminal stop codon in the query genome, which can lead to C-terminal protein extensions. TOGA1 ignores such mutations and terminates the coding region at the codon aligned to the reference stop codon (Fig. 1F, Supplementary Figure 3). In contrast, TOGA2 detects these events, searches for a downstream in-frame stop codon, and extends the terminal exon boundary accordingly. Together, these improvements increase exon annotation accuracy.

### Deep learning-based gene structure adjustments increase exon accuracy

The second major advance in TOGA2 is the use of deep learning-based splice site prediction to improve exon annotation accuracy. Although splice sites are often conserved, their positions can shift during evolution, and CESAR2 has only a limited ability to detect splice site shifts over large distances ^58,60^. Deep learning models such as SpliceAI ^61^ can accurately predict splice sites in genomic sequences. However, because SpliceAI was trained on the human genome, it is unclear whether it generalizes to other vertebrates. To assess this, we applied SpliceAI genome-wide to eight representative vertebrates (human, mouse, cow, pig chicken, zebra finch, slider turtle, zebrafish). We compared SpliceAI probabilities of NCBI RefSeq-annotated canonical U2 splice sites (GT/GC donor and AG acceptor sites) ^32^ with intronic non-splice site GT and AG positions as negative controls. Across all eight species, annotated donor and acceptor sites exhibited average SpliceAI probabilities between 86% and 94% (Fig. 2A-B, Supplementary Table 2). A probability threshold of 0.02 consistently separates real from false splice sites with 99% sensitivity and >99.3% specificity, demonstrating that SpliceAI accurately predicts splice sites across vertebrates.

**Figure 2:**
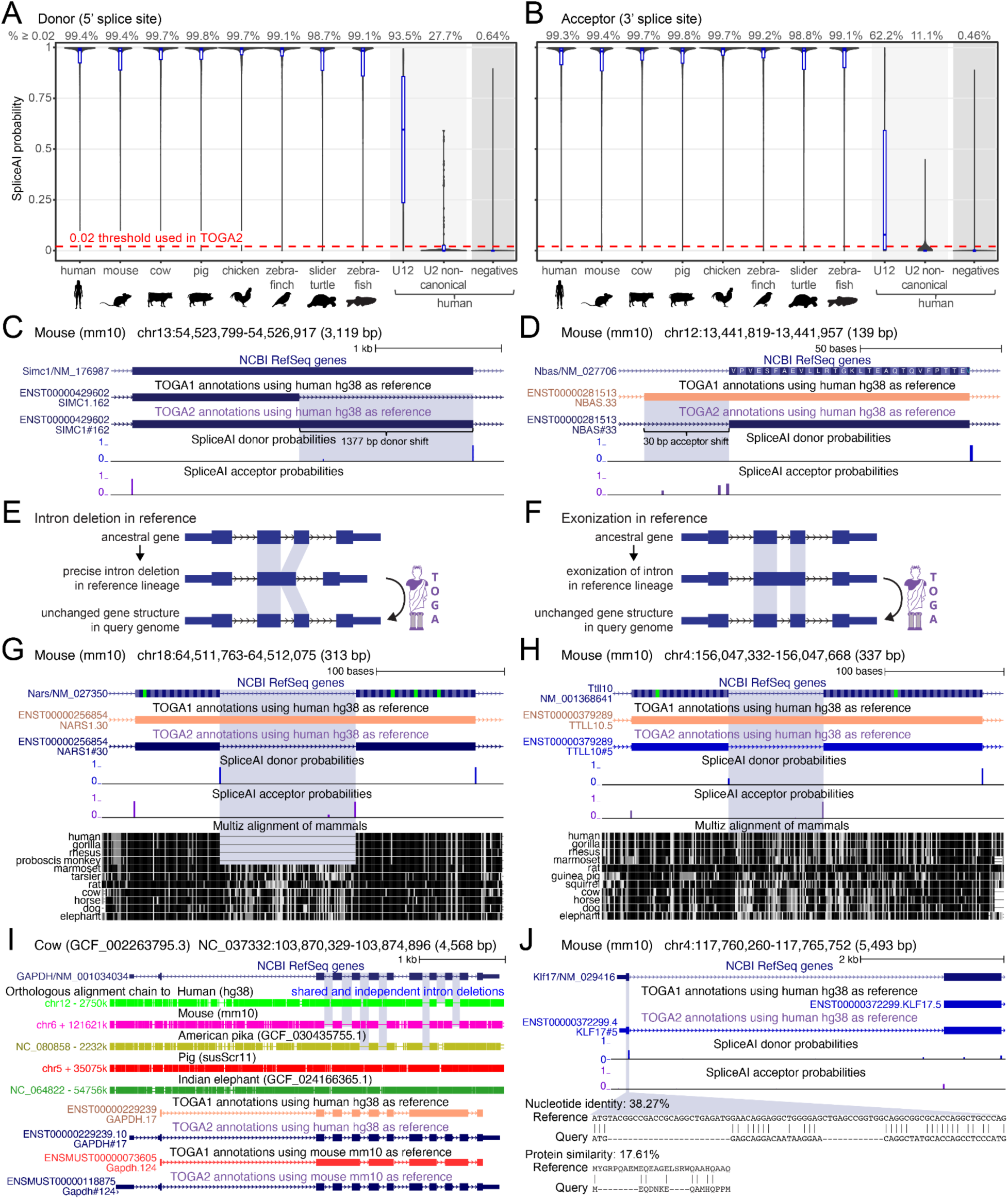
TOGA2 uses deep learning–based splice site predictions to improve exon annotation accuracy and capture gene structure changes. (A, B) Distributions of SpliceAI probabilities for donor (A) and acceptor (B) splice sites. Across the eight representative vertebrates, ≥98.7% of canonical donor (GT/GC) and ≥98.8% of canonical acceptor sites (AG) have SpliceAI probabilities ≥0.02, whereas only 0.64% of non-splice-site GTs and 0.46% of non-splice-site AGs (negatives) exceed this threshold. Non-canonical U2 and U12 splice sites exhibit markedly lower probabilities, indicating that SpliceAI has not learned their sequence patterns. (C, D) Examples of splice site shifts. (C) A large donor shift of 1,377 bp results in the orthologous mouse exon being twice as long as the human exon. Whereas TOGA1 mis-annotates the exon boundary, TOGA2 uses SpliceAI predictions to correctly annotate the exon. (D) TOGA1 fails to capture a 30 bp acceptor shift and predicts a CA acceptor site, which is interpreted as a splice site mutation, causing an uncertain loss transcript classification (orange transcript). In contrast, TOGA2 captures the shift, producing an intact transcript. (E) Illustration of precise intron deletion events. In the lineage leading to the reference, an ancestral intron is precisely deleted, resulting in the fusion of the flanking exons. Using this reference, TOGA2 must infer the correct position to insert the intron in the query genome. (F) Illustration of exonization events. In the reference lineage, an ancestral intron gained coding capacity and lost its splicing function, forming a larger composite exon. Using this reference, TOGA2 must infer which part of the exon corresponds to one or more introns in the query genome. (G) Precise intron deletion example. Intron 3 of *NARS* is deleted in Old World monkeys (blue highlight). Because TOGA1 is not aware of the query intron, it introduces a large insertion containing gene-inactivating mutations, leading to an uncertain loss transcript classification (orange transcript). In contrast, TOGA2 correctly identifies the query intron, resulting in a fully intact transcript. (H) Exonization of intronic sequence in *TTLL10*. The intronic sequence in mouse aligns to primates and is translatable in several species (e.g. orangutan, rhesus macaque, tarsier), indicating an exonization event in the lineage leading to human. Using human as the reference, TOGA1 treats its sequence as coding, which results in a frameshift and an uncertain loss transcript classification (orange). Using SpliceAI predictions, TOGA2 correctly annotates the intron, resulting in an intact transcript. (I) Genome browser view of cow, which exhibits the ancestral *GAPDH* exon-intron structure. Deletions (blue highlight) in orthologous alignment chains show that *GAPDH* exhibits multiple intron deletion events in human, mouse, and pika. Using human or mouse as references, TOGA1 fails to identify reference-absent introns and misclassifies *GAPDH* in the cow query as uncertain loss (orange) or lost (red). TOGA2 identifies all reference-absent introns and correctly annotates a fully intact *GAPDH* gene. (J) The first coding exon of *KLF17* is highly diverged in mouse due to large deletions. TOGA1 misclassifies the alignment as spurious and fails to annotate the exon, whereas TOGA2 correctly identifies it using a high-confidence donor splice site.

To leverage these predictions for gene annotation, TOGA2 provides a new mode that precomputes SpliceAI probabilities across the query genome. During transcript annotation, TOGA2 then identifies CESAR2-predicted splice sites with probabilities below the 0.02 threshold, subsequently searches for a reading frame-preserving, SpliceAI-supported splice site, and adjusts the exon boundaries accordingly.

Tested in a human–mouse TOGA2 run, splice-site correction correctly adjusted the boundaries of 452 coding exons, thereby increasing annotation accuracy. Notably, this includes exons with splice site shifts spanning more than a thousand bases (Fig. 2C, Supplementary Figure 4). In addition, this procedure prevents false splice site mutation calls, thereby improving gene integrity classification (Fig. 2D), and enables TOGA2 to correctly annotate exons with large internal insertions and deletions (Supplementary Figure 5).

Importantly, our analyses also show that splice sites of U12-type introns processed by the minor spliceosome and non-canonical U2 splice sites (e.g. TG acceptors ^62^), are not accurately predicted by SpliceAI (Fig. 2A, Supplementary Table 2), indicating that the model did not learn these rare exceptions. Instead, SpliceAI often predicts incorrect GT or AG sites in the vicinity of the true U12 or non-canonical U2 splice site, suggesting that it correctly infers the presence of a neighboring exon, but fails to identify the precise intron boundary (Supplementary Figure 6). To avoid misannotating such introns, TOGA2 pre-identifies intron types in the reference annotation using intronIC (Moyer et al. 2020) and restricts splice site correction to canonical U2 introns. To further improve U12 intron annotation accuracy, we instead refined the CESAR2 U12 splice site profiles (Supplementary Figure 7). Together with improved identification of U12 introns, these changes increase the annotation accuracy of exons flanked by U12 introns (Supplementary Figures 8-9).

### Annotating reference-absent ancestral introns

Evolutionary changes in exon-intron structure, including the presence or absence of entire introns, are a major challenge for homology-based gene annotation methods. While TOGA1 can handle precise intron deletion events in the query species, TOGA2 introduces a new functionality to also recognize ancestral introns that are absent from the reference lineage, which we found can arise through two evolutionary scenarios. The first comprises ancestral introns that were lost in the reference lineage through precise intron deletions, likely mediated by recombination with processed (intronless) pseudogenes ^59^, resulting in a fusion of the flanking exons (Fig. 2E). The second comprises events termed “exonizations”, where an ancestral intron gained coding capacity and lost its ability to be spliced, thereby becoming incorporated into a larger composite exon together with the two neighboring exons (Fig. 2F).

To correctly annotate transcripts affected by such exon-intron structure changes, TOGA2 recognizes situations where a single reference exon corresponds to multiple exons in the query species and uses SpliceAI predictions to infer the positions of one or more introns in the query genome. This results in correct annotation of the query gene structure and improves gene loss classification (Fig. 2G,H, Supplementary Figure 10).

The housekeeping gene *GAPDH* represents an extreme example. The ancestral *GAPDH* exon-intron structure, consisting of 10 introns between coding exons, is preserved in cow, pig, elephant, and other mammals. In contrast, multiple precise intron deletion events occurred in human, mouse, and pika, including deletions shared among all three species as well as up to three consecutive intron deletions (Fig. 2I). While TOGA1 cannot recognize reference-absent ancestral introns and therefore fails to infer the ancestral *GAPDH* gene structure in cow, TOGA2 identifies all introns absent from the human (three consecutive introns) or mouse (five introns in total) reference gene models and correctly annotates a fully intact *GAPDH* gene.

### Annotating highly diverged exons

TOGA2 also leverages SpliceAI information to recognize and annotate highly diverged exons. To distinguish true exon homology from random alignments arising when the homologous exon is deleted or overlaps an assembly gap, both TOGA1 and TOGA2 filter alignments based on minimum nucleotide identity (≥45%) and protein similarity (≥20%); these thresholds reliably separate true from random exon alignments with a sensitivity of 0.98 and a precision of 0.99 ^38^. However, highly diverged but genuine exons can still be misclassified as spurious alignments, leading to missing exon annotations. To address this, TOGA2 uses SpliceAI probabilities to detect and correctly annotate highly diverged exons that are supported by high-confidence splice sites.

In a human-mouse TOGA2 run, this procedure recovered and annotated 275 additional coding exons (Supplementary Figure 11). As an example, large deletions in the first coding exon of *KLF17* in mouse reduce nucleotide identity to 38.3% and protein similarity to 17.6%, causing TOGA1 to fail to annotate the exon (Fig. 2J). Because the exon is flanked by a high-confidence donor splice site predicted by SpliceAI, TOGA2 correctly annotates it.

### Evolutionarily informed transcript selection

The third major advance in TOGA2 is improved reference annotations. Because TOGA2 relies on comprehensive and accurate reference annotations, we developed an evolutionarily informed procedure to improve input transcript selection compared to TOGA1. Starting from a comprehensive initial transcript set for the reference species derived from NCBI RefSeq and, where available, GENCODE and Ensembl annotations ^32,33,63^, this approach evaluates transcript integrity across a representative set of query species. It then selects transcripts based on coding sequence length and evolutionary conservation, measured as how often their orthologs are intact. This procedure removes truncated transcripts, transcripts with non-ancestral exons, and fusion or other erroneous transcript models present in TOGA1 input annotations, while improving both quality and completeness by adding conserved exons and genes (Supplementary Figures 12-16, Supplementary Table 3).

To evaluate the impact of the improved input annotations alone, we applied TOGA1 to both the original and new human and mouse reference annotations across four query species. We quantified the number of TOGA1-annotated coding exons that precisely match (both boundaries) NCBI RefSeq-annotated exons in the query species. In all comparisons, the new input annotations recovered between 781 and 5,081 additional exons (insets in Fig. 3A-B, Supplementary Table 4), demonstrating improved annotation completeness.

**Figure 3:**
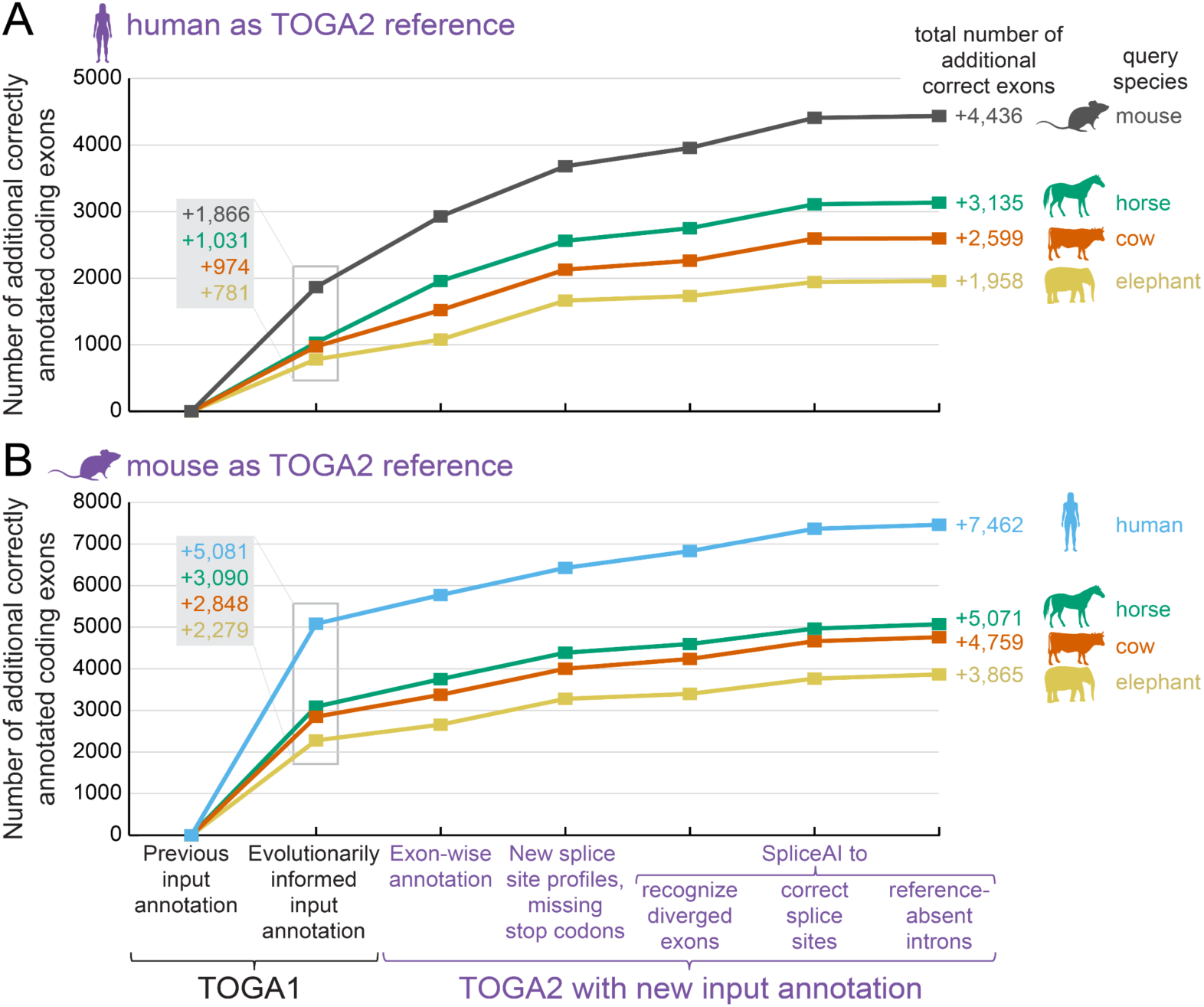
TOGA2 and new input annotations improve annotation completeness and accuracy. Comparison of the number of additional coding exons identified by TOGA1 and TOGA2 that precisely match RefSeq-annotated coding exons at both boundaries, using human (A) and mouse (B) as reference species and four mammals as representative query species. The impact of the evolutionarily informed transcript selection procedure was quantified using TOGA1 (insets). TOGA1 with the new input annotations serves as the baseline for all TOGA2 tests that use the same input annotations and enable new TOGA2 advances in a stepwise manner. The total gain in correctly annotated exons relative to the original TOGA1 setup is given on the right. Differences in the number of RefSeq-matching exons largely reflect the comprehensiveness of the benchmark annotations, with mouse and especially human having more annotated coding exons, whereas TOGA2 annotates a similar number of exons across species (Supplementary Table 4).

### TOGA2 systematically improves annotation accuracy

To systematically assess the impact of the new TOGA2 advances presented above, we used human and mouse as reference species and applied TOGA2 with the improved input annotations to four representative placental mammal query species, enabling the new advances in a stepwise manner. For each species, we determined the number of TOGA2-annotated coding exons that precisely match RefSeq-annotated coding exons, using TOGA1 with the new input annotations as a baseline.

Each TOGA2 feature increased the number of correctly annotated exons across all tests (Fig. 3, Supplementary Table 4). Overall, using human as the reference, TOGA2 correctly annotated between 1,177 (elephant) and 2,570 (mouse) additional coding exons in the query genomes. Similarly, using mouse as the reference, TOGA2 correctly annotated between 1,586 (elephant) and 2,381 (human) additional coding exons. Together, these results demonstrate that TOGA2 consistently improves annotation accuracy relative to TOGA1.

### Retrogene identification

Retrogenes are processed pseudogene copies that retain intact open reading frames and may acquire functional roles ^64,65^. To identify retrogene candidates, TOGA2 provides two advances. First, TOGA2 improves the classification of chains representing processed pseudogenes, which also enhances their discrimination from orthologous loci (Supplementary Figure 17). Second, unlike TOGA1, which did not distinguish processed pseudogene copies with intact reading frames, TOGA2 explicitly identifies these retrogene candidates and annotates them as potential query genes (Supplementary Figure 18).

### Improved orthology inference

To further refine orthology relationships, TOGA2 implements a new gene tree reconciliation approach that is applied to many:many ortholog groups and identifies additional well-supported 1:1 orthologs (Supplementary Figures 19-20). A comparison with orthology relationships provided by Ensembl Genes 115 ^63^ shows that this approach further improves the resolution of orthology relationships (Supplementary Figure 21, Supplementary Table 5).

Gene tree reconciliation and other new features, such as SpliceAI-based exon-intron structure correction, are computationally-intensive additions to TOGA2 that are absent from TOGA1. Despite this added functionality, TOGA2 remains substantially faster than TOGA1, achieving a 2.7-fold speedup (Fig. 1C).

### Improved gene loss identification

TOGA1 has been widely used to identify evolutionary gene loss events ^44–46,49,50,52,53,55–57^. By more rigorously classifying compensating frameshifts and terminal mutations (Supplementary Figures 22–23), TOGA2 enables a more accurate identification and classification of lost genes. Furthermore, TOGA2 introduces a new “Fully Intact” classification to explicitly identify transcripts with a completely intact reading frame and no missing sequence or inactivating mutations (Fig. 1A).

### TOGA2 predicts untranslated regions (UTRs)

The fourth major advance in TOGA2 is the prediction of UTRs. Whereas TOGA1 annotates only coding exons, TOGA2 additionally leverages the observation that not only coding exons but also UTR regions and exons are often conserved and align between species, enabling incorporation of aligning UTRs into the query transcript annotation (Fig. 1A, Supplementary Figure 24). To benchmark prediction accuracy, we compared TOGA2-predicted UTRs with RefSeq-annotated UTRs inferred from transcriptome data (Fig. 4A,B). Across 45 reference-query comparisons, we computed median UTR similarities for each run and found average similarities of 0.75 for 5′ UTRs (range 0.66-0.88) and 0.91 for 3′ UTRs (range 0.76-0.995), indicating that UTRs - and particularly 3′ UTRs - can often be predicted accurately. Prediction accuracy was often higher between more closely related species pairs, such as mouse–rat and cow–sheep.

**Figure 4:**
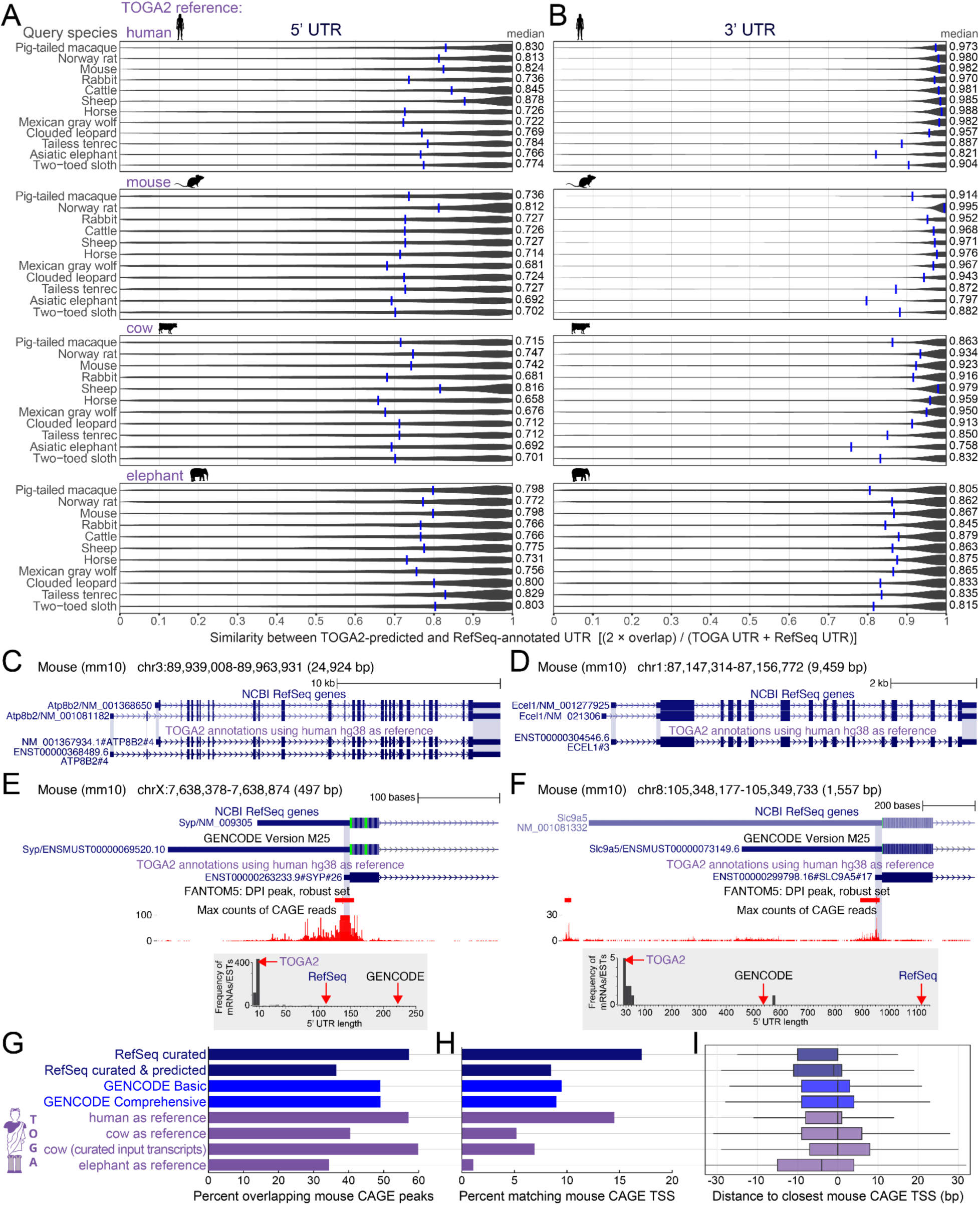
TOGA2 annotates untranslated regions (UTRs). (A, B) Violin plots showing the similarity between TOGA2-predicted and RefSeq-annotated 5′ UTRs (A) and 3′ UTRs (B), using human, mouse, cow, and elephant as references and representative placental mammals as queries. UTR similarity was calculated as twice the overlap divided by the summed lengths of the predicted and annotated UTRs, considering all UTR regions and exons. Blue tick marks indicate the median. (C, D) Examples of predicted 5′ and 3′ UTRs that perfectly match annotated UTRs (blue highlight), except for the 5′ UTR of NM_001367934.1#ATP8B2 (panel C), which has a similarity of 0.88 and is 47 bp shorter than the annotated 5′ UTR. (E, F) Two examples where shorter TOGA2-predicted 5′ UTRs are supported by CAGE (Cap Analysis Gene Expression) read counts from the FANTOM5 project ^66^ and by the majority of mRNA and EST evidence (histogram shown in the grey inset), whereas the RefSeq- and GENCODE-annotated UTRs represent rarer but longer transcript isoforms. (G, H) Percentage of RefSeq-, GENCODE-, and TOGA2-annotated TSSs overlapping mouse CAGE peaks (G) and CAGE-defined TSSs (H). (I) For annotated TSSs overlapping CAGE peaks, the distributions of distances to the closest CAGE-defined TSS is shown as boxplots (boxes indicate the first and third quartiles, the median is the vertical line, whiskers extend to 1.5 times the interquartile range).

As illustrated in Fig. 4C,D, TOGA2-predicted UTRs can perfectly or near-perfectly overlap RefSeq-annotated UTRs. We further identified cases where RefSeq- or GENCODE-annotated UTRs represent longer but rare isoforms, whereas the TOGA2-predicted UTRs are supported by the majority of transcriptomic evidence (Fig. 4E,F).

To investigate this further, we compared annotated transcription start sites (TSSs) with mouse TSSs identified by CAGE (Cap Analysis of Gene Expression) data from the FANTOM5 project ^66^. We analyzed overlap with both CAGE peaks (clusters of nearby TSSs; median peak length is 15 bp) and individual CAGE-defined TSSs (Supplementary Table 6). Of the mouse TSSs predicted by TOGA2 using human as the reference, 57.1% overlapped mouse CAGE peaks, which is comparable to curated RefSeq TSSs (57.2%) and higher than curated+predicted RefSeq (36.5%) and GENCODE TSSs (49.1%) (Fig. 4G). Furthermore, 14.5% of TOGA2-predicted mouse TSSs precisely matched CAGE-defined TSSs, which is slightly lower than the 17.1% for curated RefSeq annotations, but substantially higher than the 8.5% for curated+predicted RefSeq and 9.5% for GENCODE annotations (Fig. 4H). Among annotated mouse TSSs overlapping CAGE peaks, TOGA2-predicted TSSs showed a distance distribution to the closest CAGE-defined TSS similar to, but slightly narrower than, curated RefSeq annotations (Fig. 4I, Supplementary Figure 25). Mouse TSSs predicted by TOGA2 using cow and elephant as references also show substantial overlap with CAGE peaks (40.4% and 34.4%), although precise matches with CAGE-defined TSSs are reduced (5.2% and 1.1%; Fig. 4G-I), likely reflecting the greater phylogenetic distance and differences in input annotation quality. Indeed, when restricting the analysis to curated RefSeq cow input transcripts, the fraction of TOGA2-annotated TSSs overlapping CAGE peaks increased from 40.4% to 59.7% (Fig. 4G), indicating that input annotation quality substantially influences TSS prediction accuracy.

Together, these analyses show that aligning 5′ UTRs contain substantial information about the TSS location. Nevertheless, whenever available, transcriptomic data should be preferred for UTR annotation; in its absence, TOGA2 can generally provide accurate UTR predictions.

### TOGA2 generally achieves higher annotation completeness

We next benchmarked gene annotation completeness using compleasm ^67^, which determines the percentage of completely detected, near-universally conserved genes from clade-specific BUSCO ODB12 gene sets ^68^. We compared annotations generated by TOGA2 using one or more reference species with annotations generated by the NCBI RefSeq pipeline, which integrates transcriptomic data, homology-based data, and *ab initio* gene predictions ^32^. To avoid discrepancies between NCBI GenBank and RefSeq assembly versions, we used RefSeq assemblies (GCF_*) matching those used for TOGA2, with no genome exclusions.

We first analyzed annotations for 70 diverse placental mammals (Fig. 5A, Supplementary Figure 26, Supplementary Table 7) using four reference species ^69,70^. Using human, house mouse, cow, or Indian elephant as single references, TOGA2 achieved higher annotation completeness than RefSeq for 63 (90%), 5 (7%), 51 (73%), and 9 (13%) species, respectively. Combining annotations from all four placental mammal references reduced reference bias and further increased annotation completeness, with TOGA2 outperforming RefSeq for all 70 species (Fig. 5A). Across these 70 species, TOGA2 annotated between 17 and 183 additional completely detected genes that were either incompletely annotated due to missing conserved exons or entirely missed by RefSeq (Supplementary Figures 27-28). This high completeness is facilitated by TOGA2’s ability to correctly annotate not only typical protein-coding genes, but also genes with unusual features and complex gene structures, such as stop-codon readthrough and selenocysteine-containing genes, non-ATG start codons, distinct genes with overlapping ORFs, genes with dual ORFs, genes nested within other genes, and genes with long introns and/or hundreds of exons (Supplementary Figures 29-35).

**Figure 5:**
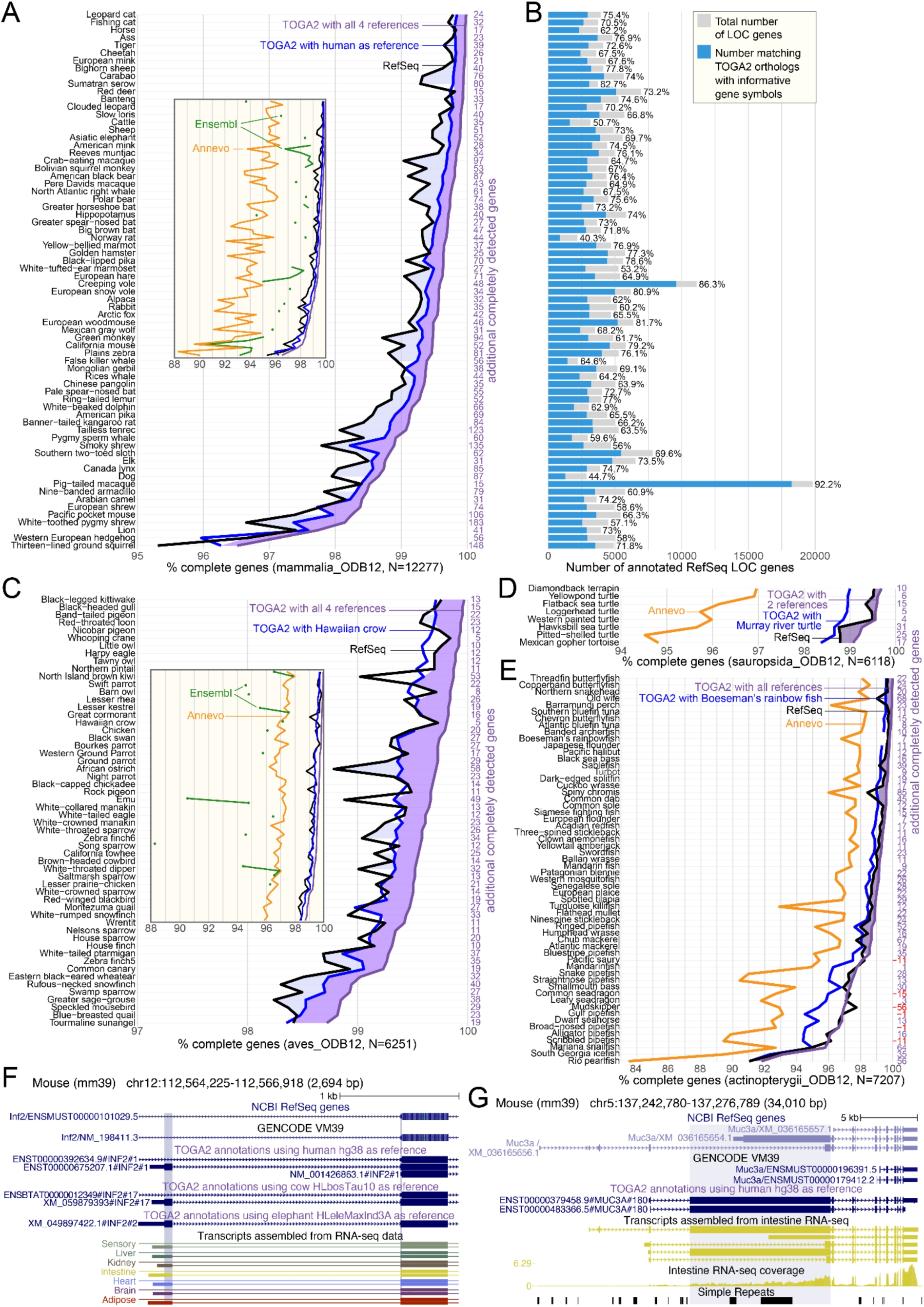
TOGA2 generally achieves higher annotation completeness. (A,C–E) Line plots show the percentage of completely detected, near-universally conserved genes from clade-specific BUSCO ODB12 gene sets in 70 placental mammals (A), 56 birds (C), eight turtles (D), and 58 percomorph fishes (E). Numbers of additional completely detected genes in combined TOGA2 annotations are shown in purple; red indicates higher RefSeq completeness. Colored areas represent the difference in completeness between NCBI RefSeq and TOGA2 using a single best reference (light blue) or combined annotations from multiple references (purple). Insets show the completeness of annotations generated by Ensembl and Annevo. Ensembl annotations are included only when available for the same genome assembly. (B) Bar charts show the total number of RefSeq genes with locus (LOC) identifiers and the subset for which matching TOGA2 orthologs provide informative gene symbols. (F) Example of an alternative first coding exon (blue highlight) annotated by TOGA2 that is supported by RNA-seq data, but absent from the mouse GENCODE and RefSeq annotations. (G) For *MUC3A*, TOGA2 annotates a large second coding exon (blue highlight) that overlaps two tandem-repeat regions and is supported by assembled RNA-seq transcripts, but is not present in GENCODE and RefSeq annotations.

We next analyzed annotations for 56 bird species (Fig. 5B, Supplementary Figure 26, Supplementary Table 8) using five reference species ^71–73^. With chicken, Hawaiian crow, or black-legged kittiwake as single references, TOGA2 achieved higher annotation completeness than RefSeq for 26 (46%), 31 (55%), and 22 (39%) species, respectively. In contrast, zebra finch and emu as single references did not outperform RefSeq, likely reflecting the lower completeness of their input annotations (Supplementary Table 3). Combining annotations from all five bird references similarly reduced reference bias and increased annotation completeness, with TOGA2 outperforming RefSeq for all 56 species, identifying up to 58 additional completely detected genes per species (Fig. 5C).

To further test annotation completeness for another vertebrate group, we ran TOGA2 for eight turtles with available RefSeq annotations, using the European pond turtle and Murray River turtle as representatives of the two major turtle suborders (Cryptodira and Pleurodira) ^74^. Combining the two reference-based TOGA2 annotations again consistently achieved higher completeness in annotating conserved genes than RefSeq annotations, adding up to 31 completely detected genes (Fig. 5D, Supplementary Figure 26, Supplementary Table 9).

Finally, we tested TOGA2 on 58 percomorph fishes representing the most diverse teleost group. These species diverged ∼115 Mya ^75^, include several rapidly evolving lineages, and exhibit more than fourfold genome size differences (0.427–1.89 Gb). We used Pacific halibut, Boeseman’s rainbowfish, turquoise killifish, and three-spined stickleback as reference species ^71,76,77^. For the highly divergent Syngnathiformes (pipefishes and seahorses), we additionally included mandarinfish as a reference. Multi-reference TOGA2 achieved higher annotation completeness than RefSeq for 90% (52 of 58) of the tested genomes, adding up to 85 additional completely detected genes (Fig. 5E, Supplementary Figure 26, Supplementary Table 10). RefSeq annotations were more complete for the remaining six species: mudskipper (-56 complete genes), Pacific saury (-11 genes), and four Syngnathiformes species (between -15 and -1 complete genes). These exceptions are partially explained by the choice of reference species and reference annotation quality. For mudskipper, a gobiiform species, we did not include a reference from this speciose and early-diverging percomorph lineage ^78^. Likewise, although mandarinfish was included as a Syngnathiform reference, its input annotation is less complete (97.74%) than those of the other percomorph references (98.63–99.72%) (Supplementary Table 3). These results suggest that adding reference species from currently uncovered, deeply diverging percomorph lineages will further improve TOGA2 annotation completeness across this highly diverse fish radiation.

We further compared Ensembl annotations generated for the same mammal and bird genome assemblies, which consistently showed substantially lower completeness than both TOGA2 and RefSeq annotations (insets in Fig. 5A,C, Supplementary Figure 26). We also benchmarked Annevo, a state-of-the-art deep learning-based *ab initio* gene prediction method ^28^, which also achieved lower completeness for mammals, birds, turtles, and fish (Fig. 5A,C-E); however, we note that *ab initio* gene prediction is inherently more challenging than evidence-based gene annotation.

In conclusion, combining TOGA2 annotations from multiple reference species generally achieves higher annotation completeness across diverse vertebrate groups than transcriptome-informed RefSeq or Ensembl annotations and deep learning-based *ab initio* predictions. This improvement arises because TOGA2 uses genes fully annotated – by RefSeq or other sources – in at least one reference genome to annotate many other species, whereas transcriptome-informed annotations are generated independently for each genome, and insufficient transcriptome coverage can therefore lead to incomplete annotations. As an illustration, even though cow is used as a TOGA2 reference species, combining TOGA2 annotations with human, mouse, and elephant as references detects 35 additional complete genes in cow compared to RefSeq.

### TOGA2 multiple reference integration

Motivated by the increased annotation completeness achieved by combining multiple reference species, we developed a TOGA2 integration mode to reduce reference bias. Reference bias arises when the reference lineage has lost genes or exons present in other species, or when reference annotations fail to include conserved genes, preventing their annotation in the query genome. TOGA2 integration merges multiple reference-based annotations for the same query into a single annotation by removing non-conserved gene predictions while retaining transcript diversity, thereby mitigating reference bias and increasing both annotation accuracy and completeness.

### TOGA2 identifies unannotated exons in mouse

Given TOGA2’s high annotation completeness, we next investigated whether TOGA2 can identify coding exons and exon variants not represented in the current mouse GENCODE VM39 annotation, despite mouse being a well-annotated model organism. Using human, cow, and elephant as references, we identified 4,564 TOGA2-annotated exons for which one or both splice sites were absent from the mouse GENCODE annotation (Supplementary Table 11). Of these 4,564 candidate exons, 39.2% (1,790) are supported by two or all three reference species, and 21.7% (989) are annotated by RefSeq. To further assess whether these candidates represent genuine mouse exons, we intersected them with transcripts assembled from available RNA-seq data. Of the 4,564 candidate exons, 78.2% (3,566) were confirmed by RNA-seq, indicating that they represent bona fide coding exons. These represent a diverse set of previously unannotated splicing events, including alternative exons, alternative introns, and alternative donor or acceptor splice sites (Fig. 5F, Supplementary Figures 36-37).

An interesting example is the intestinal mucin gene *MUC3A*. In human, *MUC3A* contains a large second coding exon of 8,805 bp. Using human as the reference, TOGA2 predicts a fully intact mouse transcript containing a substantially enlarged second coding exon of 12,390 bp (Fig. 5G), consistent with previous reports of a >11 kb exon ^79^. Using available intestinal RNA-seq data, we assembled several transcripts, the longest of which precisely matches the TOGA2 annotation, supporting the existence of this full-length transcript. In contrast, both GENCODE and RefSeq do not annotate this *MUC3A* isoform: GENCODE gene models represent incomplete annotations that begin only at the sixth coding exon, whereas RefSeq predicts two transcripts, neither of which captures the large 12,390 bp exon. TOGA2 additionally identifies this long-exon-containing transcript across taxonomically diverse placental mammals, where it is frequently incompletely annotated by RefSeq, suggesting that it represents an ancestral but difficult-to-annotate transcript (Supplementary Figure 38). Together, these results indicate that TOGA2 can uncover additional conserved coding exons absent from current reference annotations.

### TOGA2 provides informative gene symbols for thousands of orthologs

TOGA2 not only produces accurate and comprehensive gene annotations, but also implicitly identifies orthologous relationships between genes. Unlike TOGA1, TOGA2 leverages these relationships to assign informative query gene names based on orthologous reference genes, also for non-1:1 orthologs (Supplementary Figure 39). Inspired by ^80^, we investigated whether TOGA2 can identify orthologs and assign informative gene names to RefSeq-annotated genes with automatically assigned, uninformative locus identifiers (LOC gene symbols).

Across 70 placental mammals, an average of 3,496 (median 3,063) RefSeq LOC genes could be assigned informative gene symbols with TOGA2 orthologs (Fig. 5A, Supplementary Table 12). Even for the model organism rat, the RefSeq annotation contains 2,176 LOC genes, of which 876 (40.3%) match TOGA2-annotated orthologs with informative gene symbols. These include fragmented RefSeq gene models, retrogene-derived annotations, and genes such as *CUPIN1*, which is lost in human and mouse but correctly annotated and named in cow (Supplementary Figure 40). An extreme case is the pig-tailed macaque, for which the RefSeq annotation contains 19,802 LOC genes, of which 18,252 (92%) can be assigned informative gene symbols by TOGA2. This shows that TOGA2 systematically resolves uninformative LOC symbols into biologically interpretable gene names.

### Annotation of V(D)J segments of immunoglobulin and T cell receptor genes

Immunoglobulins (antibodies) and T-cell receptors (TCRs) are critical components of the adaptive immune system that rely on hyperdiverse gene sequences to recognize antigens. This diversity is generated by somatic V(D)J recombination, in which variable (V), diversity (D), and joining (J) gene segments, flanked by conserved recombination signal sequences (RSSs), are randomly rearranged and joined to form functional antigen receptors ^81^. V(D)J segments are short, highly similar to one another, exhibit allelic variation, and undergo non-templated modifications, making them notoriously difficult to annotate.

Because our sensitive alignment chains capture V(D)J segments (Supplementary Figure 41), we developed a dedicated TOGA2 module to annotate antigen receptor loci. Candidate segments are identified by sequence similarity, RSSs are detected, and functional segments are distinguished from pseudogenes based on the presence of both an RSS and an intact open reading frame (Figure 6, Supplementary Figure 42). We compared mouse annotations generated by TOGA2 and IGDetective, a specialized method for V(D)J annotation ^82^, against the GENCODE, RefSeq, and IMGT annotations ^32,63,83^.

**Figure 6:**
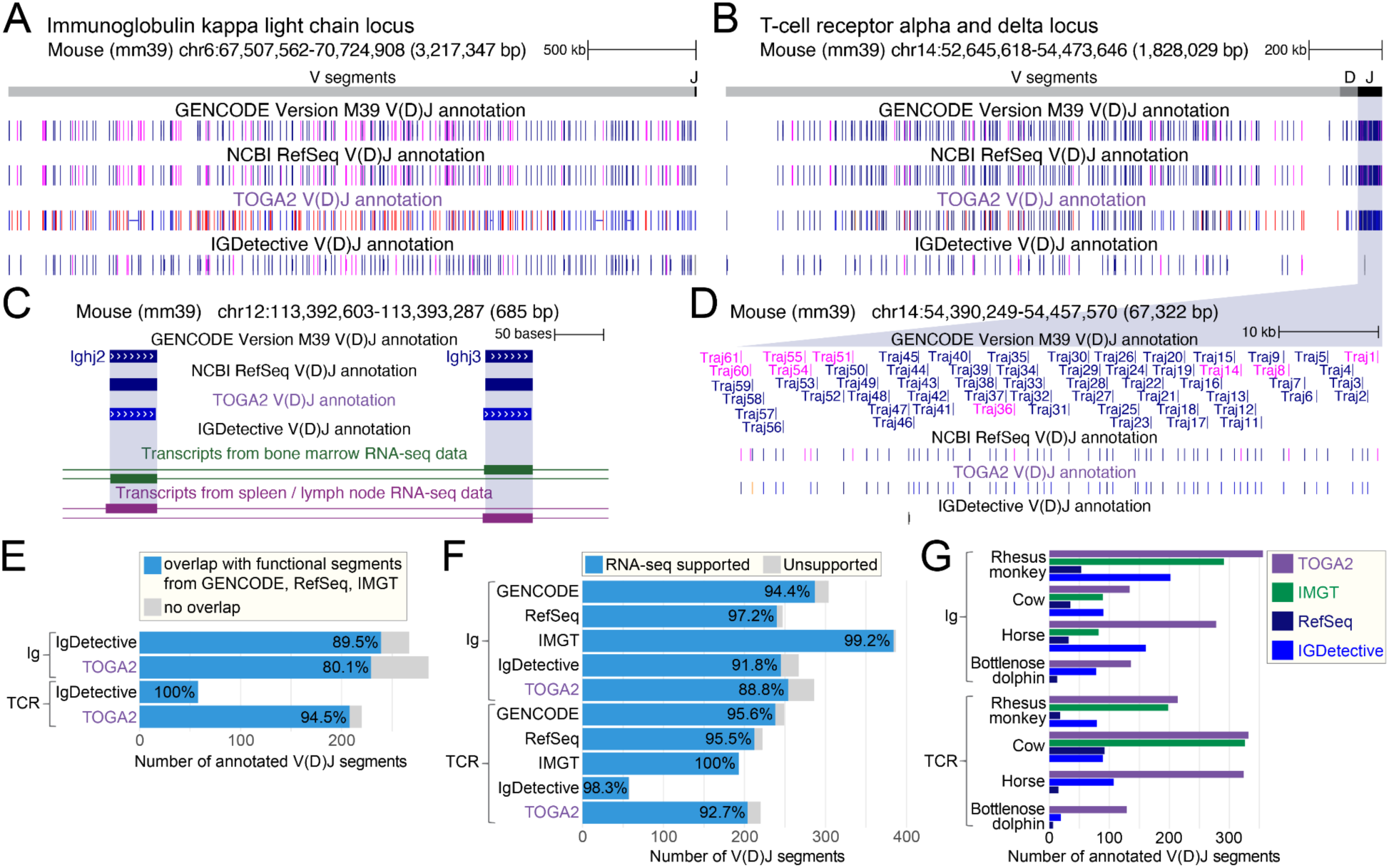
TOGA2 annotates V(D)J segments of antigen receptors. (A-B) Genome browser screenshot showing V(D)J segments in the mouse immunoglobulin kappa light chain (A) and the T-cell receptor alpha/delta locus (B). The TOGA2 annotation agrees well with GENCODE, RefSeq and IGDetective, but IGDetective reports only a subset of the segments in the T-cell receptor locus (B). (C-D) Examples of J segments in the immunoglobulin heavy (C) and T-cell receptor alpha locus (D) identified by TOGA2 but not by IGDetective. (F) Overlap between V(D)J segments annotated by TOGA2 and IGDetective in mouse and the corresponding annotations in GENCODE, RefSeq, and IMGT. (G) The large majority of V(D)J segments annotated by TOGA2, IGDetective, and other annotation sources in mouse are supported by RNA-seq data. (H) Number of V(D)J segments annotated in other species. IMGT is not available for all species and does not annotate T-cell receptor segments for horse.

We found that both TOGA2 and IGDetective annotations showed high agreement with existing annotations, and the large majority of identified V(D)J segments are supported by mouse RNA-seq data (Figure 6A-B,E-F). However, TOGA2 detected numerous annotated V(D)J segments, particularly T-cell receptor segments, that were missed by IGDetective (Figure 6B-D, Supplementary Figure 43). TOGA2 also identified segments with putative alternative RSSs that are supported by transcriptome data (Supplementary Figure 44). We note, however, that TOGA2 has limitations in identifying short and highly diverged D segments and can misannotate segments with extreme sequence divergence (Supplementary Figure 45).

Applied to other species, where V(D)J loci are generally less completely annotated, TOGA2 consistently identified more segments than IGDetective and, in particular, RefSeq, while showing better agreement with available IMGT annotations (Figure 6G). These results indicate that TOGA2 can facilitate V(D)J annotation in non-model species, providing a foundation for comparative immunogenomics studies.

### Annotations and orthology resources for thousands of vertebrate genomes

Leveraging that TOGA2’s reference-based annotation strategy scales linearly with respect to the number of query species, we applied the method to thousands of vertebrate genomes, including all genomes used in the annotation completeness benchmarks as well as 230 high-quality assemblies generated as part of VGP Phase I ^37^. Using the above-described taxonomically representative reference species, we applied TOGA2 a total of 9,376 times across 1,069 mammalian assemblies (covering 788 placental and 20 non-placental species), 867 bird assemblies (800 species), 68 turtle assemblies (60 species), and 158 percomorph fish assemblies (156 species) (Fig. 7, Supplementary Tables 14-17). For each query species, annotations generated from multiple references were also merged using TOGA2 integration mode.

**Figure 7.**
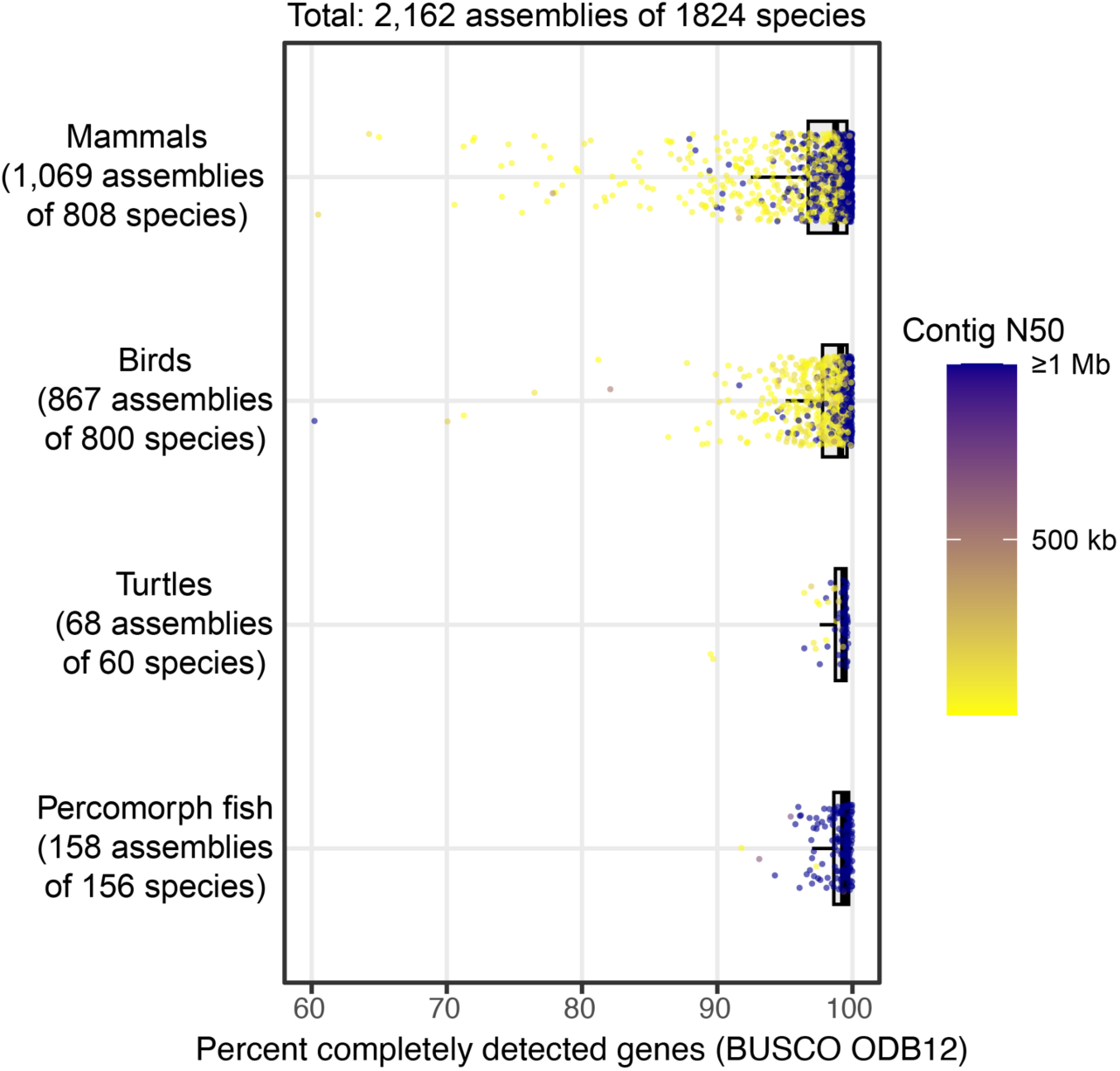
TOGA2 gene annotations across 2,162 vertebrate genomes. Percentage of completely detected, near-universally conserved genes from clade-specific BUSCO ODB12 gene sets for integrated TOGA2 annotations. Each dot represents one assembly and is colored by contig N50. Integrated TOGA2 annotations generally achieve high completeness across these 2,162 assemblies, with lower completeness primarily associated with lower assembly contiguity.

For each of the 2,162 assemblies, TOGA2 generated comprehensive resources, including gene annotations with UTR predictions in BED and GTF format; orthologs and orthology relationships; gene duplication data (Supplementary Figure 46); gene loss classifications together with the underlying inactivating mutations (Supplementary Figure 22); processed pseudogenes and retrogene candidates; nucleotide and protein sequences; and nucleotide and protein alignments to the reference. We additionally generated clade-wide sets of 1:1 orthologs for mammals, birds, turtles, and fish. Summary statistics, including the number of inferred query genes, retrogenes, gene loss events, and orthologs, are provided in Supplementary Tables 14-17.

For all assemblies, we provide genome browser tracks at https://genome.senckenberg.de, visualizing annotations using the TOGA2-generated bigBed files together with UCSC decorator tracks ^84,85^, which highlight inactivating mutations (Supplementary Figure 47). Clicking on an annotation entry provides additional information, including a visualization of the exon-intron structure with inactivating mutations, features used for orthology and transcript classification, nucleotide and protein sequences, and protein and exon-level alignments. We also provide genome-wide SpliceAI splice site predictions in bigWig format for the 2,162 assemblies, together with a total of 9,373 pairwise genome alignments to the reference species.

Together, these datasets constitute, to our knowledge, the largest comparative annotation resource currently available for these vertebrate clades and provide a foundation for numerous comparative and evolutionary analyses, including those described below.

### Application range of TOGA2

TOGA2 also provides a set of tools to directly facilitate downstream analyses of the generated gene annotations and orthology relationships. First, we provide *toga2orthogroups* to infer multi-species orthogroups (gene families) from TOGA2 orthology assignments by modeling reference-query orthology relationships as a graph and extracting connected components using the Union-Find algorithm. Optionally, a gene family database such as PANTHER ^86^ can be incorporated to further consolidate orthogroups by adding family-level edges. The resulting orthogroups can be directly used with CAFE5 ^87^ to analyze gene family expansions and contractions.

Second, we provide *toga2agora*, which automatically extracts and reformats gene orthology and synteny information from TOGA2 results given a species tree and uses it for ancestral gene order reconstruction with AGORA ^9^. We previously used this strategy to infer ancestral karyotypes and reconstruct the evolution of genetic sex determination in turtles ^57^.

Third, TOGA2 generates nucleotide FASTA files of all annotated transcripts, along with gene membership and orthology relationships. We provide *toga2kbpython* to leverage these data for comparative gene expression analyses using kb-python and DESeq2 ^88,89^, including genes with non-1:1 orthology relationships, thus enabling the inference of gene expression shifts across species.

Fourth, using an updated set of ancestral placental mammal genes (defined as human genes with conserved orthologs in Afrotheria and Xenarthra), *toga2stats* leverages TOGA2’s gene classification to benchmark query genome assembly quality by quantifying the proportion of ancestral genes with missing sequence or inactivating mutations.

## Discussion

By delivering accurate and comprehensive coding gene annotations together with orthology relationships, gene loss and duplication events, processed pseudogenes and retrogene candidates, codon- and protein-level alignments, and standardized outputs for downstream analyses, TOGA2 provides a versatile, multi-purpose framework for comparative genomics. Compared to its predecessor, TOGA2 leverages exon-level orthology, gene tree reconciliation, and deep learning based gene structure adjustment, among other advances, thereby markedly reducing runtime and memory requirements while improving the accuracy of exon annotation and orthology inference. By annotating UTRs, TOGA2 further improves gene model completeness.

Achieving high annotation completeness requires comprehensive input annotations, which TOGA2 provides through a new, evolutionarily informed transcript selection procedure. With a single comprehensive input annotation, TOGA2 achieves higher annotation completeness than transcriptome-informed and homology-based annotation pipelines or *ab initio* deep learning methods for many query species. To reduce reference bias, TOGA2 can be run with multiple reference species, which generally achieves higher completeness, except for very divergent query species. Even in the model organism mouse, TOGA2 identifies exons missing from current reference annotations. Finally, TOGA2 reliably annotates antigen receptor V(D)J gene segments, identifying previously unannotated segments across all tested species. By addressing the growing gap between genome assembly and annotation, TOGA2 provides a scalable solution for large-scale comparative genomics. We demonstrate this by applying TOGA2 to thousands of vertebrate genomes, generating the largest comparative gene resources to date.

The quality of the reference annotation is a key determinant of TOGA2’s performance. This is exemplified by the large exon isoform of *MUC3A*, which seems to be correctly annotated only in human, but rarely identified in other mammals (Fig. 5G, Supplementary Figure 38). Using human as reference, TOGA2 can systematically propagate this high-quality annotation to other species, suggesting that accurate and complete reference annotations strongly influence gene inference across the focal clade. Accordingly, maximizing annotation quality within a lineage would benefit strongly from investing in generating highly accurate annotations for the reference species.

Transcriptomic data has the advantage of providing direct evidence for genes, enabling the identification of lineage-specific exons and genes, and providing accurate UTR annotations. However, because many genes show condition-specific expression patterns, transcriptomic-based annotations can be incomplete when transcriptomic data do not comprehensively cover tissues, cell types, and conditions. In contrast, homology-based annotations, as provided by multi-reference TOGA2, are not limited by available transcriptomic data, but cannot identify lineage-specific exons or genes. Therefore, homology-based annotation and available transcriptomic data are ideally combined to generate comprehensive and accurate annotations of both conserved and lineage-specific genes.

## Materials and Methods

### TOGA2 input and output files

As input, TOGA2 requires (i) the reference and query genomes in 2bit format (an indexed and compressed format that can be generated from a fasta file using the UCSC Genome Browser utility faToTwoBit), (ii) the genome alignment between reference and query in chain format, (iii) SpliceAI donor and acceptor probabilities for both strands of the query genome in bigWig format, (iv) CESAR2 splice site profiles for canonical and non-canonical U2 and U12 splice sites, and (v) the reference input annotation. The reference annotation consists of (i) a BED12 file describing exon-intron structures of coding transcripts, (ii) an isoform file specifying which transcripts belong to the same reference gene in tsv format, and (iii) a BED6 file listing reference introns with non-canonical U2 splice sites or U12 splice sites, as inferred by IntronIC.

Genome alignment chains can be generated using the make_lastz_chains pipeline (https://github.com/hillerlab/make_lastz_chains), SpliceAI probabilities can be computed with the TOGA2 module “spliceai” or our containerized solution in https://github.com/hillerlab/containers, and all files required for the reference annotation can be generated with the TOGA2 module “prepare-input”.

Although possible, running TOGA2 without SpliceAI data is not recommended, as splice site and exon-intron structure correction cannot then be performed. If no isoform file is provided, TOGA2 assumes that each transcript represents a separate gene. If no intron classification file is provided, TOGA2 assumes that all reference introns contain canonical U2 splice sites.

TOGA2 provides rich output, including: (i) the query gene annotation in BED12 and GTF format, with and without predicted UTRs; these files include retrogene candidate and paralogous gene annotations; (ii) annotation tracks in bigBed format together with UCSC decorator tracks highlighting inactivating mutations, ready for visualization in the UCSC Genome Browser ^84,85^; (iii) a BED9 file listing processed pseudogenes detected in the query genome; (iv) lists of inferred query genes based on orthologous, paralogous, and retrogene candidate projections in TSV and BED format; (v) TSV files describing gene orthology relationships (1:1, 1:many, many:many, etc.) between reference and query, together with orthology probabilities for all projections; (vi) loss status classifications for genes, transcripts, and projections, as well as lists of inferred gene-inactivating mutations (both in TSV format); (vii) protein and codon alignments for all annotated transcripts in FASTA format; (viii) protein and nucleotide sequences for all annotated transcripts in FASTA format; and (ix) per-exon nucleotide alignments in 2bit files, used to generate multiple codon alignments. A detailed description of these and additional output files is available on the TOGA2 wiki (https://github.com/hillerlab/TOGA2/wiki/Output-structure).

### TOGA2 pipeline overview

Apart from TOGA1’s machine-learning classifier for orthologous chain identification ^38^, all other parts of the TOGA2 pipeline have been newly implemented in Python and Rust to improve speed and incorporate numerous new advancements. The TOGA2 pipeline contains the following major steps:

1. Initialization: All input files and third-party utilities undergo sanity checks, and the output directory structure is created.
2. Feature extraction: For each reference transcript, TOGA2 identifies alignment chains that overlap or span at least one coding exon. The features used for orthology classification are extracted for every transcript-chain pair (termed projection). Unlike TOGA1, TOGA2 runs this step in a single thread because of substantially improved feature extraction speed.
3. Orthology classification: The gradient-boosting machine learning classifier is used to compute orthology probabilities for all projections. The improved processed pseudogene identification procedure is also performed at this step.
4. Projection preprocessing: For all projections eligible for annotation, the preprocessing steps extracts the reference exon sequences, query genomic sequences, and the corresponding query SpliceAI probabilities and stores them in a temporary HDF5 file. Memory requirements for the subsequent CESAR2 alignment are also estimated. This step consists of multiple independent jobs that are executed in parallel.
5. Exon-wise annotation and transcript structure adjustment: CESAR2 is used to align reference coding exons and identify their loci and boundaries in the query genome. The resulting alignments are subsequently processed and filtered, highly diverged exons are identified, SpliceAI-mediated splice site and exon–intron structure correction is performed, inactivating mutations are detected, and the projection loss status is assigned. This step is executed as multiple parallel jobs, with memory estimates from the preprocessing step used to bin jobs into memory buckets.
6. Query gene inference: Query genes are inferred from projections, taking orthology status, loss status, paralogous projections, and retrogene candidates into account. This step includes a new chain ranking and filtering procedure to reduce false orthology assignments.
7. Loss status inference: Projection loss statuses are used to infer the loss status of reference transcripts and genes, generating the corresponding loss classification table.
8. Orthology relationship inference: Orthology relationships between reference and query genes (1:1, 1:many, many:many, 1:0, etc.) are identified and refined using the weak-edge graph reduction algorithm.
9. Gene tree reconciliation: The gene tree reconciliation procedure is optionally applied to many:many ortholog groups to identify additional well-supported 1:1 orthologs and update orthology relationships. This step uses parallel jobs for gene tree inference and reconciliation.
10. Post-processing: At this step, TOGA2 (i) annotates UTRs, (ii) uses a new procedure to assign query gene names, (iii) filters annotation and sequence files, and (iv) uses postoga (https://github.com/alejandrogzi/postoga) to generate summary tables.
11. Genome browser tracks: BigBed files and UCSC decorator tracks (Supplementary Figure 47) are generated to enable visualization of annotations and associated metadata in the UCSC Genome Browser.

TOGA2 is modular, allowing users to stop pipeline after any step and restart from any completed step, provided that all files required for downstream analyses are available.

### Faster feature extraction

To identify orthologous query loci from co-linear local alignment chains, TOGA2 extracts characteristic features capturing intronic and intergenic alignments. To improve runtime relative to TOGA1, the feature extraction code was reimplemented in Rust using the chaintools library (https://github.com/alejandrogzi/chaintools). In a genome-wide human vs. mouse run, the TOGA2 implementation requires only ∼70 seconds for feature extraction, representing a ∼30-fold speedup compared with the 36.4 minutes required by TOGA1. TOGA2 uses the same classifier model and classification features as TOGA1.

### Improved discrimination of processed pseudogenes from orthologous loci

TOGA2 introduces an improved procedure to identify processed pseudogenes from alignment chains and distinguish them from orthologous loci. TOGA1 identified processed pseudogenes by applying a post hoc classification step only to chains classified as non-orthologous and filtering for chains with an “alignment-to-query span” value ≥0.95. This feature was computed as *e/Q*, where *e* is the number of reference exonic bases (CDS and UTR) that align via the chain to the query and *Q* is the span of the entire chain in query coordinates. UTR alignments were included because UTR sequence is also reverse transcribed and therefore typically present in processed pseudogenes. However, we found that this procedure fails to identify processed pseudogenes when reference transcripts differ in their UTR structure and can even cause TOGA1 to incorrectly classify processed pseudogenes as orthologs (Supplementary Figure 17).

To overcome these problems, TOGA2 implements a different procedure for processed pseudogene identification based on two new chain features and applies it to projections of multi-exon transcripts classified as either orthologous or non-orthologous. The first new feature is the *clipped intron coverage*, computed as *i/S*, where *i* is the number of aligned intronic bases for introns located between coding exons of the focal transcript, and *S* is the length of the intersection between the transcript CDS span and the chain span in the query genome. By considering only introns between coding exons, this feature avoids misclassifying aligned UTR sequence of alternative isoforms as intronic alignments. The second feature is *clipped exon coverage*, computed as *e/L*, where *e* is the total number of reference coding exon bases from all transcripts that align to the query through this chain, and *L* is the distance in query coordinates between the start of the most upstream aligning CDS exon and the end of the most downstream aligning CDS exon, considering all transcripts. Together, these features capture characteristic properties of processed pseudogene alignments, which contain little or no intronic alignments and predominantly consist of coding sequence. By default, processed pseudogenes are identified as projections with *clipped intron coverage* <0.1 and *clipped exon coverage* >0.3. This revised procedure improves both processed pseudogene detection and orthology classification accuracy (Supplementary Figure 17).

### Exon-wise annotation and dynamic search space extension

TOGA1 defines the search space of a transcript in the query genome as the entire orthologous locus inferred from the alignment chain and annotates all exons simultaneously using the multi-exon mode of CESAR2 ^58^. In contrast, TOGA2 leverages exon-level orthology, which drastically reduces the search spaces, and invokes a series of single-exon CESAR2 runs. To this end, TOGA2 infers the positions of individual exons in the query genome from the alignment chain, which can be applied to the majority of exons that align at the nucleotide level and are bounded by flanking intronic alignments. For each exon, TOGA2 adds flanking regions of user-defined length (100 bp by default) on both sides to enable splice site detection and to allow CESAR2 to capture small-distance splice site shifts. If an exon does not fully align, TOGA2 estimates the extent of the unaligning sequence and extends the search space accordingly. The resulting query genome interval is referred to as the exon’s search space. The projection search space is then defined as the genomic span between the start of the most upstream exon search space and the end of the most downstream exon search space.

Single-exon mode has two limitations. First, it cannot detect precise intron deletion events in the query, where an intron is removed, likely through germline recombination with intronless processed pseudogenes ^58,59^, resulting in the fusion of two or more exons. Second, some highly diverged exons do not align at the nucleotide level and are therefore absent from the alignment chain, although they may be detectable at the protein level by CESAR2. To address these limitations, TOGA2 identifies candidate cases of intron deletion by detecting neighboring query exons with minimal inter-exon distances. For these cases, TOGA2 performs a multi-exon CESAR2 run spanning the query locus that covers all affected exons. Similarly, TOGA2 groups unaligning exons with flanking exons that do align and applies a multi-exon CESAR2 run to the query locus that covers this exon group and their intervening introns. By combining the strengths of both single- and multi-exon modes, TOGA2 substantially improves memory usage and runtime efficiency, while retaining the ability to detect precise intron deletions and highly diverged exons that are only identifiable at the protein level.

TOGA2 also implements a new feature to dynamically extend the search space when the orthologous alignment chain does not cover terminal exons. Such exons are more difficult to recover because terminal unaligning exons lack a flanking aligned exon on one side (Supplementary Figure 1A). Similarly, exons located outside the boundaries of the alignment chain are challenging to detect (Supplementary Figure 1B). To annotate these exons, TOGA2 estimates the length of the exonic and intronic sequence based on the reference transcripts, multiplies this estimate by a given factor (1.2 by default), and extends the search space in the direction of the unaligning exon(s) accordingly. Running CESAR2 on the extended search space enables the recovery of additional exons, as illustrated in Supplementary Figure 1. Search space extension is not applied to projections classified as processed pseudogenes, because processed pseudogenes are frequently truncated and the absence of terminal exons in the alignment chains therefore often reflects genuine sequence loss.

Misalignments can result in alignment chains where one or more terminal exons are mapped far upstream or downstream of the main gene locus. To prevent exon misannotations, TOGA2 identifies projections containing putative introns that are at least fivefold larger than the corresponding reference intron and longer than 500 kb. For these projections, TOGA2 retains the chain segment covering the majority of the transcript’s coding sequence and extrapolates the search space toward the missing side.

### Splice site profiles and reference intron type classification

TOGA2 features an improved approach to distinguish introns recognized by the major (U2) and minor (U12) spliceosomes, which is necessary because CESAR2 requires splice site profiles that capture the respective splice site sequence patterns (Supplementary Figure 7). Whereas TOGA1 simply assumed that all reference introns with splice site dinucleotides other than the GT-AG consensus are U12 introns, TOGA2 relies on more accurate reference intron classifications using IntronIC ^90^, which improves annotation accuracy (Supplementary Figure 8).

Furthermore, TOGA2 distinguishes between canonical and non-canonical U2 and U12 splice sites, because they differ in their sequence patterns. Canonical U2 splice sites contain GT or GC dinucleotides at the donor site and AG at the acceptor site, whereas canonical U12 splice sites contain GT dinucleotides at the donor site and AG at the acceptor site. Because individual introns can contain both canonical and non-canonical splice sites, TOGA2 assigns canonicality separately to donor and acceptor sites. For example, a U2 intron with a GT-GG dinucleotide profile is considered to have a canonical donor site but a non-canonical acceptor site. Intron type information is provided as part of the input annotation in BED format. If no intron type annotation is supplied, TOGA2 assumes that all reference introns are canonical U2 introns.

To obtain CESAR2 splice site profiles for the eight different donor and acceptor splice site classes, users can use the TOGA2 input preparation module, which infers these profiles based on the IntronIC classification of reference introns. In addition, we provide CESAR2 profiles generated from introns contained in the human annotation sources used to construct the reference annotation prior to the evolutionarily informed transcript filtering step (Supplementary Figure 7). To reduce the impact of misannotated introns, introns shorter than 70 bp were excluded from the training dataset. Using human as reference and mouse (mm10) as query, we observed improved U12 intron annotation accuracy when relaxing sequence constraints in the non-canonical U12 acceptor splice site profile (Supplementary Figure 9). These modified profiles were used for all TOGA2 runs in this study.

### Capturing mutated stop codons

Mutations of the terminal stop codon in a query species can lead to C-terminal extensions of the encoded protein. In contrast to TOGA1, TOGA2 detects these events and searches for a downstream in-frame stop codon within the original alignment search space. If such a stop codon is identified, TOGA2 extends the boundary of the terminal exon to this position and adds the respective insertion to the exon alignment.

### Using SpliceAI for exon identification and splice site correction

By default, TOGA2 uses SpliceAI probabilities to annotate highly diverged exons, correct boundaries of canonical U2 introns, and identify reference-absent introns (below). SpliceAI donor and acceptor probabilities are precomputed for each query genome on both strands and provided to TOGA2 as four bigWig files.

TOGA2 explicitly distinguishes between introns recognized by the major (U2) and minor (U12) spliceosomes, because U12 and non-canonical U2 splice sites are not accurately predicted by SpliceAI. For all reference input transcripts, intron type (U2 or U12) is inferred using intronIC. Canonical U2 splice sites contain GT or GC dinucleotides at the donor site and AG at the acceptor site, whereas canonical U12 splice sites contain GT at the donor site and AG at the acceptor site. Canonical splice sites are considered mutated when the dinucleotide differs from the consensus. A splice site is considered supported by SpliceAI when its SpliceAI probability is ≥0.02, which is a user-adjustable threshold.

TOGA2 implements several splice site correction modes, which are additive such that each successive mode enables correction of additional splice site classes. The currently implemented modes support: (i) no splice site correction (SpliceAI is used only to identify highly diverged exons); (ii) correction of mutated canonical U2 splice sites; (iii) correction of SpliceAI-unsupported canonical U2 splice sites; (iv) correction of mutated canonical U12 splice sites; (v) correction of SpliceAI-unsupported canonical U12 splice sites; (vi) correction of SpliceAI-unsupported non-canonical U12 splice sites; and (vii) correction of SpliceAI-unsupported non-canonical U2 splice sites. Although users can select alternative splice site correction modes, we currently recommend the default settings, which restrict correction to canonical U2 splice sites without attempting to correct U12 or non-canonical U2 splice sites, because these splice site classes are not accurately predicted by SpliceAI. Nevertheless, implementation of the different correction modes supports all intron classes, enabling future use of deep-learning-based splice site predictors that capture U12 and non-canonical U2 splice sites more accurately.

To improve the annotation of highly diverged exons compared to TOGA1, TOGA2 determines whether internal exons have SpliceAI support for both splice sites and whether terminal exons have support for their single splice site. Exons fulfilling this criterion are classified as present and annotated regardless of their sequence similarity to the reference exon.

Splice site correction is performed independently for each donor and acceptor site in the query sequence, except in cases of precise intron deletion. For each splice site subjected to correction, TOGA2 searches for an alternative candidate site satisfying the following criteria: (i) the candidate splice site must lie within the search space of the adjacent exon; (ii) the candidate site must preserve the reading frame of the adjacent exon, unless the exon already contains a frameshifting mutation; (iii) the candidate site, or combination of candidate donor and acceptor sites, must not introduce an in-frame stop codon; (iv) the candidate site must not completely eliminate the adjacent exon; (v) the candidate site or their combination cannot reduce the intron length to less than ten nucleotides; and (vi) if the CESAR2-annotated splice site has a non-zero SpliceAI probability but below the threshold, the candidate site must exceed this probability by at least a user-defined margin (default 0.02). Criteria (iii) and (iv) assure that splice site correction neither introduces in-frame stop codons in the new exonic sequence when the splice site is shifted into the intron, nor creates stop codons across the newly formed exon-exon boundaries.

If multiple splice site combinations satisfy these criteria, TOGA2 selects the site with the highest SpliceAI support. If both the donor and acceptor sites are corrected, TOGA2 first selects the best-supported donor, followed by the best-supported valid acceptor. For exons whose boundaries are modified during splice site correction, the exon alignment is updated accordingly by introducing insertions when the splice site shifts into the intron and deletions when the splice site shifts into the exon. These induced insertion or deletion events are ignored during exon classification but may affect coding sequence integrity statistics and, consequently, the inferred projection loss status.

### Using SpliceAI for identification of reference-absent introns

To capture exon–intron structure changes, TOGA2 additionally uses SpliceAI predictions to identify reference-absent introns that are present in the query genome but absent from the reference transcript due to intron deletion or exonization events (Fig. 2E,G). In the TOGA2 output files, such events are referred to as gained introns in the query, reflecting the reference perspective, although the intron is typically absent from the reference species. Identification of reference-absent introns is performed independently of and before splice site correction and can be disabled by the user.

TOGA2 tests each CESAR2-aligned exon for the presence of one or more candidate reference-absent introns, defined as pairs of SpliceAI-supported donor and acceptor sites. To reduce spurious intron predictions, TOGA2 applies a stringent donor and acceptor probability threshold (0.8 by default). TOGA2 further uses two types of mutational patterns as evidence supporting the presence of a genuine reference-absent intron. First, if the corresponding reference intron was precisely deleted, the query intron is represented as a large insertion in the exon alignment. TOGA2 therefore uses the presence of one or more insertions with a combined length of at least 30 bp within the candidate intron region as the first supporting feature. Second, if exonization occurred in the reference lineage, the corresponding intronic sequence in the query is often not translatable. TOGA2 therefore uses the presence of frameshifting insertions/ deletions or in-frame stop codons within the candidate intron region as a second supporting feature. Because these features increase the likelihood of a genuine reference-absent intron, TOGA2 lowers the SpliceAI probability threshold to 0.2 if one supporting feature is present and to 0.1 if both features are present. All threshold values are user-adjustable.

In addition to SpliceAI support, candidate introns must satisfy the following criteria. First, removal of the candidate intron must preserve the reading frame of the downstream exon part, unless intron removal resolves pre-existing frameshifting mutations. Second, intron removal must not introduce an in-frame stop codon at the newly formed exon-exon boundary. Third, the candidate intron must not occupy more than a user-adjustable proportion of the reference exon length (50% by default).

To obtain updated exon alignments after adjusting the exon-intron structure, TOGA2 removes the identified introns from the reference and query sequences and realigns the resulting exon structure with CESAR2 using canonical U2 splice site profiles. Candidate introns are rejected if the resulting alignment introduces frameshifts or inframe stop codons, or reduces exon nucleotide identity by more than 10% in comparison to the original alignment.

TOGA2 allows the identification of multiple reference-absent introns within a single exon, exemplified by *GAPDH*, which exhibits multiple consecutive intron deletions in both human and mouse (Figure 2). In such cases, the criteria described above must be satisfied both for each individual intron and for the combined exon-intron structure. By default, TOGA2 allows up to four reference-absent introns per exon. To identify the optimal exon-intron structure among multiple candidate configurations, TOGA2 evaluates all compatible intron combinations and selects the structure with the highest overall nucleotide identity.

### Improved loss status classification

TOGA2 uses a loss status classification system similar to TOGA1, but introduces new categories and updated classification criteria:

● Fully Intact (FI): This status is new in TOGA2 and indicates that the projection has a completely intact reading frame without any missing sequence or inactivating mutations.
● Intact (I): This status refers to projections for which the middle 80% of the CDS is present (not missing from the assembly) and lacks inactivating mutations. However, mutations may be present in the N- or C-terminal 10% of the CDS and may indicate alterations of the protein’s termini.
● Partially Intact (PI): For these projections, at least 50% of the CDS is present and the middle 80% of the CDS lacks inactivating mutations. In contrast to intact projections, parts of the CDS are missing from the assembly, potentially due to assembly gaps. Such projections often still encode functional proteins, but the evidence is weaker.
● Uncertain Loss (UL): This status indicates the presence of at least one mutation treated as inactivating; however, the evidence is insufficient to conclude that the projection can no longer encode a functional protein. The term “uncertain” reflects that such projections may represent (i) early stages of gene loss, where not enough inactivating mutations have accumulated; (ii) rare assembly base errors (false mutations); or (iii) conserved genes for which individual exons are no longer conserved or were mispredicted by TOGA2.
● Lost (L): This status indicates strong evidence that the projection can no longer encode a functional protein. Loss is typically inferred from the presence of multiple inactivating mutations affecting at least two exons or from extensive sequence deletions. Exon lengths are taken into account when a single exon constitutes a substantial fraction (≥40%) of the CDS. TOGA2 implements a revised and improved loss inference procedure compared to the one used in TOGA1.
● Missing (M): This status refers to projections for which less than 50% of the CDS is present and the middle 80% of the CDS lacks inactivating mutations. In such cases, there is currently no evidence for gene loss; however, confidence is lower because more than half of the CDS is absent. Missing projections can also arise when no genome alignment chain spans the transcript.

The Partially Missing (PM) category used in TOGA1 is no longer used in TOGA2 because of its ambiguous interpretation. In addition, TOGA2 defines two special categories for non-orthologous projections annotated in the query genome:

● ParaloGous (PG): In the absence of orthologous projections, TOGA2 annotates transcripts using chains classified as paralogous. This scenario may reflect the loss of the true ortholog in the query lineage or a rare orthologous chain misclassification. Paralogous projections in the query are only retained if the corresponding query locus lacks orthologous projections, classifying their coding sequence integrity as for orthologous projections, except that projections otherwise classified as L or M are assigned the PG status instead. Transcripts and genes in the reference genome that are represented exclusively by paralogous projections are assigned PG status regardless of coding sequence integrity in the query.
● Processed Pseudogene (PP): This status is new in TOGA2 and refers to projections classified as processed pseudogene copies. As for paralogous projections, retrotransposed transcript copies are assigned integrity statuses ranging from FI to UL based on their coding sequence integrity, but projections that would otherwise be classified as L or M are assigned PP status to indicate genuine processed pseudogenes. This category is exclusive to query transcript annotations and does not propagate to reference transcript or gene levels.

Finally, the special category Undefined (N) is assigned to items discarded for technical reasons prior to the alignment step, preventing evaluation of coding sequence integrity.

### Improved identification of gene loss events

Several improvements for identifying gene losses were introduced after the publication of TOGA1.0. First, TOGA2 implements an improved handling of compensating frameshifts. In TOGA1, all compensating frameshifts were not considered inactivating, because they restore the reading frame without introducing a stop codon in the frameshifted region. However, in rare cases, compensating frameshifts span large portions of the transcript (Supplementary Figure 22). Because the region between these frameshifts is translated into a protein sequence that is likely very different from the reference protein, treating the transcript as intact is often inaccurate. TOGA2 addresses this by treating the region between compensating frameshifts as a breakpoint in the reference reading frame when computing the longest fraction of the coding sequence that lacks inactivating mutations.

Second, TOGA2 improves the distinction between UL and L by “unmasking” otherwise masked nonsense, frame-shifting, and splice site mutations, including compensated frame shifts, when a transcript already contains at least one unmasked inactivating mutation. In such cases, masked mutations, including N- or C-terminal mutations, often provide additional evidence for gene loss. Therefore, TOGA2 considers these mutations as inactivating when evaluating the overall loss status of the transcript (Supplementary Figure 22).

Third, an improved classification of N-terminal mutations is used. Inactivating mutations located within the first 10% of the reading frame are typically masked, as a downstream inframe ATG codon can often serve as an alternative translation initiation site; however, this is not always the case. TOGA2 now identifies the next inframe ATG located downstream of a potential inactivating mutation. If this ATG is in the first 10% of the reading frame, the mutation remains masked; otherwise, it is treated as inactivating, because the resulting protein would lack at least 10% of its N-terminal sequence (Supplementary Figure 22).

Fourth, TOGA2 further improves the handling of spanning chains, defined as alignment chains that span a reference gene without any aligning blocks overlapping coding exons of that gene. TOGA1 ran CESAR2 on the query locus defined by the closest upstream and downstream aligning blocks of each spanning chain, resulting in substantial runtime overhead. TOGA2 no longer runs CESAR2 on spanning chains, because our alignment chains are generated using highly sensitive alignment parameters and RepeatFiller ^91,92^, making it unlikely that a genuine gene exists at such loci. If no trustworthy exon alignments are identified, TOGA1 classifies the gene as missing if the corresponding query locus contains an assembly gap, and otherwise as lost. However, when multiple spanning chains cover the same gene, TOGA1 considers all of them and classifies the gene as missing if any spanning query locus contains an assembly gap, which can result in incorrect loss status assignments. To address this issue, TOGA2 selects the most likely orthologous chain by identifying the “most nested spanning chain”, defined as the chain whose closest upstream and downstream aligning blocks span the smallest reference interval around the gene (Supplementary Figure 23). The query locus defined by this chain is then used to determine whether the gene is classified as missing or lost.

### Query gene inference

While it is known for the reference which transcripts belong to which gene, TOGA2 must infer which projections belong to the same gene in the query genome. Query gene inference in TOGA2 follows the same general strategy as TOGA1 but includes several improvements. In general, projections that overlap in the query genome by at least one coding exon base on the same strand are assigned to the same query gene. Unlike TOGA1, which did not distinguish processed pseudogene copies with intact reading frames (retrogene candidates), TOGA2 explicitly identifies retrogene candidate projections and uses them to infer potential query genes (Supplementary Figure 18). However, retrogene candidate projections and paralogous projections are discarded if they overlap orthologous projections. Consequently, query genes inferred from retrogene or paralogous projections do not contain orthologous projections. Processed pseudogene projections lacking an intact reading frame are not used for query gene inference, but such projections are removed from the processed_pseudogenes.bed output file if they overlap orthologous, paralogous, or retrogene projections.

TOGA2 uses search space extension to annotate additional terminal exons that are not covered by the genome alignment chain. Because this procedure can occasionally introduce incorrect overlaps between terminal exons, thereby confounding query gene inference, TOGA2 records which exons were aligned in the alignment chain and tests for same-strand coding exon overlap only for those exons (but see Supplementary Figures 48 and 1B for exceptions to this rule).

Another improvement upon TOGA1 is a chain ranking and filtering procedure that reduces false orthology assignments. For each reference transcript with multiple orthologous projections (orthology probability ≥ 0.5 by default), TOGA2 first selects the projection with the highest orthology probability that is classified as non-missing, and considers it as a reliable orthologous projection. To retain co-orthologous query loci, additional projections are also considered as reliable and retained in the final annotation if they have an equal or higher orthology probability or at least 60% of their CDS is covered by the genome alignment chain. The remaining projections frequently represent orthology mispredictions that confound orthology relationship classification. TOGA2 therefore filters these questionable projections as follows. Questionable projections are removed if they overlap at their query locus either (i) a reliable projection of the same or a different reference gene or (ii) a questionable projection of a different reference gene. Questionable projections that overlap another questionable projection of the same reference gene or that do not overlap any other projection are retained only if they are classified as FI or I.

In certain cases, transcripts assigned to distinct reference genes already have same strand coding exon overlap in the reference annotation (Supplementary Figures 15 and 31). Because these transcripts also overlap in the query genome, they mislead query gene inference by merging both genes into a single inferred query gene, thereby producing incorrect many:1 orthology relationships. Unlike TOGA1, TOGA2 incorporates prior knowledge of overlapping reference genes into the gene inference procedure and infers separate query genes for the overlapping projections, resulting in the correct 1:1 orthology relationships. Additional rare situations that complicate query gene inference, together with the strategies implemented by TOGA2 to resolve them, are illustrated in Supplementary Figures 48 and 1B.

### Gene tree-based orthology resolution

TOGA2 utilizes the graph-based orthology resolution approach implemented in the original TOGA, including the weak-edge reduction algorithm to identify 1:1 orthologs from confounded many:many orthology groups. To further improve orthology resolution, TOGA2 additionally implements a gene tree reconciliation approach that is applied to many:many orthology groups and aims at identifying additional well-supported 1:1 orthologs. Our analyses showed that additional 1:1 orthologs are rarely identified in large orthology groups containing more than 50 genes and that such groups account for most of the runtime. Therefore, as a default, gene tree reconciliation is restricted to many:many orthology groups containing at most 50 genes (user-adjustable threshold) after the initial orthology graph resolution step.

To infer gene trees, TOGA2 selects a single protein sequence per gene based on projection loss status, sequence length, and alignment chain score. Frameshifting mutations are masked with ‘X’ symbols in the protein sequences. Selected sequences must be at least 50 amino acids long, must not consist of a homopolymer of a single amino acid, and must contain at least 10% defined amino acids (non-‘X’ characters). To minimize false-positive 1:1 ortholog assignments, TOGA2 subjects a many:many group to gene tree reconciliation only if all genes provide sequences satisfying these requirements.

Protein sequences are aligned with PRANK ^93^ using default settings. The resulting alignments are used to reconstruct gene trees with IQ-TREE2 ^94^ using the parameters ‘--seqtype AA -T AUTO --seed 12345 --subsample-seed 12345 --keep-ident’. By default, model selection in IQ-TREE2 is restricted to the following amino acid substitution models: JTT, WAG, JTTDCMut, Q.LG, Q.pfam, Q.pfam_gb, and Q.mammal. Ultrafast bootstrap approximation is performed using the IQ-TREE2 ‘-B’ parameter, with the number of bootstrap replicates adjustable by the user (5,000 by default). The resulting gene trees are rooted using midpoint rooting.

To infer additional well-supported 1:1 orthologs, we devised a bottom-up node-labeling algorithm that traverses the gene tree from the leaves toward the root and classifies each internal node into one of five categories (Supplementary Figure 19). Leaf nodes are labeled as R for reference genes or Q for query genes. Internal node labels are then inferred recursively based on their two child nodes, using the following symmetric rules.

First, labels Dr and Dq represent reference and query gene duplication events, respectively, and are assigned according to these two rules:

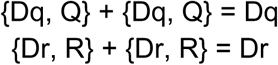

Second, the label S represents speciation events:

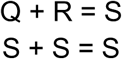

Third, the label P represents a problematic node, which is assigned to nodes that connect a speciation node and a subtree containing only reference or query genes:

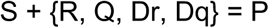

This label is used to prevent removal of orthology relationships that would otherwise orphan a remaining gene in the many:many group.

Finally, the label M represents mixed events that cannot be unambiguously classified as speciation or lineage-specific duplication events:

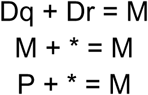

where * refers to any node label.

After all nodes have been labeled, TOGA2 identifies additional 1:1 orthologs as reference-query gene pairs that (i) are connected by an edge in the original orthology graph, (ii) are represented by a speciation (S) node that connects two leaf nodes, (iii) satisfy a bootstrap support threshold (90% by default), and (iv) have no problematic node on the path between the speciation node and the tree root (Supplementary Figure 19).

Newly inferred 1:1 orthologs are removed from the original many:many group, and orthology relationships connecting these genes to other genes are discarded from the annotation. TOGA2 then uses the pruned gene tree to reevaluate orthology relationship types for the remaining genes, which were originally classified as many:many. These relationships may subsequently be reclassified to 1:many or many:1 if a single reference or query gene retains several co-orthologs, or to 1:0 if a reference gene no longer retains chain-supported orthologs in the pruned gene tree.

To evaluate accuracy, we used human as reference and several representative placental mammal species (chimpanzee, rhesus macaque, mouse, rat, cow, and horse) as query species (Supplementary Table 5). We extracted all additional 1:1 orthologs identified by the gene tree reconciliation step and compared them to orthology relationships specified in Ensembl Genes 115 ^63^ (accessed May 19, 2026) (Supplementary Figure 21).

### Annotating untranslated regions (UTRs)

TOGA2 leverages information contained in pairwise genome alignment chains to annotate UTRs of query transcripts. UTR annotation is performed after coding sequence annotation, using the same alignment chain used for coding exon annotation. From this chain, TOGA2 identifies aligning UTR regions, infers the corresponding query coordinates, and adds valid UTRs to the coding gene model, as illustrated in Supplementary Figure 24. Valid UTRs cannot overlap annotated coding sequences, 5′ UTRs must be located upstream of the coding region and 3′ UTRs downstream. TOGA2 predicts UTRs only when the corresponding terminal coding exon is annotated.

We distinguish between two classes of UTR-containing reference exons. First, complete UTR exons consist entirely of UTR sequence. For these exons, TOGA2 identifies the query coordinates corresponding to the first and last aligning bases in the alignment chain and adds the resulting UTR exon to the query transcript model, if it satisfies the validity criteria (Supplementary Figure 24A). Second, exons containing both UTR and coding sequence are handled separately, because the CDS portion is already annotated and TOGA2 only needs to infer the UTR boundary on one side. For exons starting with UTR (UTR:CDS), TOGA2 infers the query coordinate corresponding to the first aligning UTR base and extends the query exon start to this position, while leaving the CDS unchanged (Supplementary Figure 24A). Likewise, for exons ending with UTR (CDS:UTR), TOGA2 infers the query coordinate corresponding to the last aligning UTR base and extends the query exon end accordingly.

If the start or end of a UTR region or exon does not align, TOGA2 performs an optional length extrapolation step that users can disable via a parameter setting. In this procedure, TOGA2 calculates the length of the unaligning reference UTR segment and extends the corresponding UTR region or exon in the query proportionally (Supplementary Figure 24B). If the extrapolated region overlaps an upstream or downstream UTR exon, the UTRs are merged.

Large insertions or alignment errors can produce excessively large predicted UTRs or UTR introns. To avoid such artifacts, TOGA2 discards UTR annotations that (i) exceed an absolute length of 3,000 bp or (ii) are more than 2.5-fold longer than the corresponding reference UTR. In addition, excessively large inferred introns can lead to UTR exon misannotations. To avoid these artifacts, TOGA2 discards UTR exon candidates for which the distance to the corresponding CDS boundary (i) exceeds 5,000 bp and (ii) is more than fivefold larger than in the reference. These thresholds are user-adjustable. Both filters are applied sequentially to all UTR regions and exons, starting with the CDS-proximal ones, and are applied both before and after length extrapolation.

### Query gene naming

TOGA2 assigns names to query genes based on orthology relationships to reference genes. For 1:1 orthologs, the query gene inherits the name of its orthologous reference gene. For genes duplicated in the query (1:many), the different query genes also inherit the name of the orthologous reference gene, but different copies are distinguished by appending lowercase letter suffixes (_a, _b, _c, etc.).

Unlike TOGA1, which assigns uninformative region (reg_) gene identifiers for all query genes, TOGA2 also assigns informative names for these genes. For many:1 relationships, TOGA2 distinguishes how many reference orthologs are associated with a query locus. If only two or three reference genes correspond to the same query locus, the names of the orthologous reference genes are concatenated as a comma-separated list. Otherwise, TOGA2 selects a representative reference gene and appends a “+” symbol to its name. For many:many orthology groups, TOGA2 follows the same strategy as for many:1 relationships, but appends copy number suffixes (_1, _2, etc.) to distinguish multiple query genes (Supplementary Figure 39). Overall, this naming strategy preserves information about orthology relationships while providing informative and compact gene identifiers.

### TOGA2 run summary

TOGA2 provides a summary file reporting key annotation statistics, including the number of query genes identified, the numbers of lost and missing genes, and the number of reference genes having orthologs in the query (Supplementary Figure 49). To determine the number of query genes, TOGA2 counts all genes inferred from orthologous projections with loss status FI, I, PI, or UL. In addition, genes inferred only from paralogous projections with loss status FI, I, PI, or UL are also counted, because these loci provide evidence for the presence of a gene despite the absence of orthologous projections. Retrogene candidates are reported separately because it is generally unknown whether these loci are transcribed and therefore represent bona fide genes. TOGA2 also reports the numbers of query genes classified as Lost and Missing. Finally, the summary reports how many reference genes have at least one orthologous query gene and how many reference genes are represented only by paralogous projections in the query genome.

### Integration of multiple reference-based annotations

TOGA2’s integration mode reduces reference bias, arising from genes or isoforms absent from a reference lineage, by merging multiple reference-based annotations for the same query species into a single integrated annotation. Unlike a simple union, the integration procedure removes compromised and false-positive gene predictions while preserving transcript diversity, thereby increasing both annotation accuracy and completeness. Users can specify a reference priority order to preferentially retain projections from certain references; for example, transcripts annotated using human as the reference can be prioritized over otherwise identical projections generated from other references because their gene names are often more informative.

The first step of the integration procedure is conceptually similar to query gene inference in TOGA2. All projections generated from the different reference species are used to construct an overlap graph, connecting two projections if they overlap by at least one coding exon base on the same strand and if the overlapping exons have alignment support by the genome alignment chain. As in the gene inference procedure, retrogene candidates annotated using one reference are discarded if they overlap orthologous or paralogous projections generated from other references. Likewise, paralogous projections overlapping orthologous projections are removed unless they exhibit loss status specified by the user (FI or I by default).

The resulting overlap graph is split into connected components representing candidate query genes containing projections from all references. Projections derived from reference genes that already have same strand coding exon overlap in the reference annotation, and which therefore also overlap in the query genome, are separated into distinct query genes. For each candidate query gene, projections are subsequently filtered to remove duplicate, false-positive, and spurious paralogous annotations in three steps. First, orthologous projections are filtered according to their loss status, retaining projections with the highest-confidence status and marking the remaining projections for potential removal. If no projection has a functional loss status (FI, I, PI, and UL by default), projections with the highest-ranking loss status are retained to also retain genes inferred to be lost in the query species. Second, for projections with identical CDS coordinates, the integration mode retains only the projection with the higher-confidence loss status or, if the loss statuses are identical, the projection derived from the higher-priority reference specified by the user. Third, to preserve transcript diversity, the integration mode evaluates whether paralogous projections with FI or I status, or orthologous projections marked for removal, contribute coding sequence not represented by the currently retained projections. Such projections are then retained if they introduce a novel coding exon with a length of at least 15 bp, or extend an existing exon by at least 15 bp and by at least 30% relative to its original length (both user-adjustable thresholds). The retained projections are then split again into connected components representing query genes to resolve situations in which discarded projections, for example transcripts classified as lost, artificially merged distinct genes during the initial graph construction step.

Integrated query genes are named by concatenating the reference gene names that contribute projections. For example, because human gene nomenclature is generally more informative than elephant gene nomenclature, users may assign higher priority to human-derived annotations. Because orthology is fundamentally a pairwise relationship and integration considers one query species together with multiple references, the integration procedure does not infer orthology relationships or gene losses.

TOGA2’s integration mode outputs the query projections in BED12 format. If UTR annotation was performed, this BED12 file will contain UTRs. The corresponding reference species is added as a prefix to each projection name to indicate their origin. Nucleotide and amino acid sequences are provided in FASTA format. Files describing inferred query genes and query gene-to-projections mappings are output in BED format. If genome browser tracks are requested, TOGA2 additionally produces integrated annotation bigBed tracks together with corresponding decorator tracks.

### Annotation of V(D)J segments with TOGA2

To obtain the input annotation of V(D)J segments for the human hg38 reference genome, we integrated segment annotations from Ensembl (Homo_sapiens.GRCh38.116.gtf), RefSeq (GCF_000001405.40_GRCh38.p14_genomic.gff) and IMGT (https://www.imgt.org/, last accessed on 5.12.2025) ^32,63,83^. Because IMGT does not provide genomic coordinates, we mapped the *Homo sapiens* sequence entries to the hg38 genome using BLAT version 36x2 with default parameters ^95^ to infer their genomic locations. To avoid incorrect annotations, we excluded segments that could not be mapped unambiguously to the genome. Ensembl served as the primary annotation source because it generally provided the most accurate segment boundaries and included leader exons for most V segments. Because leader sequences are not annotated in IMGT, IMGT annotations were used only when corresponding Ensembl and RefSeq annotations were not available. To maximize the completeness of the V(D)J input annotation, we additionally included pseudogene segments with intact ORFs in the input annotation.

To annotate these segments, TOGA2 uses special parameter settings “-no_utr -k --paralogs_over_spanning --max_chains_per_transcript 1000 --min_chain_score 3000 --min_orth_chain_score 3000 --skip_gene_trees --no_u12_file --no_isoform_file --no_spliceai”, which disable UTR annotation, skip gene trees, prioritize paralogous projections over spanning chains, increase the maximum number of candidate chains to 1,000 per segment, and lower the minimum chain score threshold to 3,000. These settings increase sensitivity for segments with numerous candidate alignments and allow smaller alignment chains to be considered.

V(D)J segments represent a special class of coding exons. Unlike conventional exons that are bounded by splice sites, functional V segments are flanked by a recombination signal sequence (RSS) at their 3′ end, functional J segments at their 5′ end, and functional D segments at both ends. Therefore, TOGA2 evaluates the presence of RSS motifs as key evidence for functional segments. We used previously described heptamer and nonamer motifs from ^82^. In addition, we considered downstream heptamers of V and D segments that start with *CAC* and upstream heptamers of D and J segments that end with *GTG*, as these conserved nucleotides are critical for recombination activity ^96,97^. Because RSS motifs are directly adjacent to V(D)J segments, TOGA2 first selects the predicted segment whose boundary is closest to the best-matching RSS motif within the projected region. If multiple segment candidates have equally well-supported boundaries, the longest segment is selected. Finally, segment boundaries are adjusted to exactly match the identified RSS boundary.

Because functional antigen receptor genes are assembled through genomic V(D)J recombination, several criteria used by TOGA2 to assess integrity of normal protein-coding genes are not applicable. To assess reading-frame integrity, TOGA2 considers only internal stop codons, irrespective of where they occur within a segment. Because V(D)J segments can contain frameshifting mutations (Supplementary Figure 42), frameshifts are not considered inactivating mutations, unlike in conventional protein-coding genes.

Each segment is then classified based on the presence of an RSS and its reading-frame integrity. TOGA2 defines four segment classes, two of which represent likely functional segments. Fully intact segments have an intact reading frame and both the RSS heptamer and nonamer motifs are present (shown in dark blue in the browser track). Intact segments have an intact reading frame and a detectable heptamer, but no identifiable nonamer motif (shown in blue in the browser). These segments are still considered likely functional because the heptamer is the more conserved and functionally critical component of the RSS, whereas the nonamer is more variable^98^. The remaining two classes capture likely non-functional segments. Non-intact segments lack an intact reading frame (shown in red), whereas RSS-lacking segments have an intact reading frame but lack a detectable heptamer motif (shown in orange).

Because most VDJ segments are covered by several alignment chains, resulting in many:many relationships, TOGA2 reduces the number of segment annotations per query locus as follows. First, only the annotated segment with the highest classification is retained, using the ranking fully intact > intact > RSS-lacking > non-intact. Second, if multiple annotations with the same classification remain, TOGA2 keeps the longest segment.

### Benchmarking annotations of V(D)J segments

We ran IGDetective v1.1.0 ^82^ with default settings for all benchmark assemblies. To distinguish functional and pseudogenized segments, we used the “Productive” classification given by IGDetective, which indicates whether a segment can be translated without internal stop codons. Because this classification is provided only for V segments, all D and J segments were treated as functional.

To compare TOGA2 and IGDetective predicted segments and existing annotations, we extracted all annotated V(D)J segments from NCBI RefSeq Other annotations of GCF_000001635.27 (mouse), GCF_002263795.3 (cow), GCF_049350105.2 (rhesus monkey), GCF_041296265.1 (horse), and GCF_011762595.1 (bottlenose dolphin), using the “pseudogene” classification to distinguish functional and pseudogenized segments. For mouse and cow, we additionally used IMGT annotations, which were mapped to the mm39 and HLbosTau10 (GCF_002263795.3) genomes using BLAT as described above. For mouse, we also used GENCODE VM39 annotations obtained from the UCSC Genome Browser tables wgEncodeGencodeCompVM39 and wgEncodeGencodePseudoGeneVM39. Overlap between predicted and annotated mouse segments was assessed using the UCSC tool overlapSelect with parameters -strand and - overlapSimilarity=0.5, requiring strand-specific overlap with a similarity of at least 50%. Similarity is defined in as (2*overlap length in bp) / (predicted segment + annotated segment).

To assess RNA-seq support of segments predicted in mouse mm39, we considered each V, D, and J segment individually. Because TOGA2, RefSeq, and GENCODE annotate the leader exon of V segments, but IMGT and IGDetective do not, leader exons were excluded from this analysis to ensure a fair comparison. We then split assembled RNA-seq transcripts into exons using the UCSC tool bedToExons and removed redundancy by merging exons with identical coordinates. RNA-seq support was assessed using the UCSC tool overlapSelect with parameters -strand and -overlapSimilarity=0.8, requiring strand-specific overlap between a segment and an RNA-seq exon with a similarity of at least 80%, defined as (2*overlap length in bp) / (segment + exon length).

### Repeat masking and generating alignment chains

For newly processed reference or query genomes, we generated a *de novo* repeat library using RepeatModeler (http://www.repeatmasker.org/, parameter -engine NCBI) and used the resulting library to soft-mask the genome with RepeatMasker v4.0.9. Because the genomes of Boeseman’s rainbowfish and turquoise killifish contain extensive tandem repeat regions ^99^ that are not adequately masked by RepeatModeler/RepeatMasker, we added repetitive regions identified by WindowMasker and Satellite Repeat Finder ^100,101^ to the repeat softmask.

We generated pairwise genome alignments between reference and query genomes following previously established procedures ^102^. Briefly, local alignments were obtained using LASTZ v1.04.15 ^103^ with parameters (K = 2400, L = 3000, Y = 9400, H = 2000, and the LASTZ default scoring matrix) that provide sufficient sensitivity to align orthologous exons between placental mammals ^91^. Local alignments were then chained using axtChain ^39^ with default parameters, except for linearGap=loose. To recover missed alignments overlapping repetitive regions, we applied RepeatFiller ^92^ with default parameters. To improve alignment specificity, we used chainCleaner ^104^ with default parameters, except for minBrokenChainScore = 75,000 and - doPairs.

### SpliceAI benchmark across vertebrates

We obtained the NCBI RefSeq annotation for representative vertebrate genomes including human (hg38) from the UCSC genome browser, using either the curated subset (table ncbiRefSeqCurated) or all (curated and predicted; table ncbiRefSeq) annotations. For pig and dog, we used both the curated and all annotations to compare how the results depend on the annotation type. We then extracted all splice sites (donor and acceptor), considering only introns located between coding exons. We used intronIC ^90^ to determine U12 introns and U2 introns with non-canonical splice sites, and kept only U2 introns with canonical (GT/GC donors, AG acceptors) splice sites. We then extracted the SpliceAI donor and acceptor probabilities for real splice sites on their respective strand.

We constructed a negative dataset, consisting of intronic non-splice site positions that start with an GT (negative donors) or end with an AG (negative acceptors). To this end, we used the human curated NCBI RefSeq annotation, extracted all introns located between coding exons, and excluded 20 bp from each splice site. Since introns of one transcript can overlap exons present in other transcripts, we constructed a comprehensive set of potential exons by pooling exons contained in the RefSeq (curated and predicted), GenCode V45 comprehensive and GenCode V45 pseudogene annotation. We extended each exon by 20 bp on both sides to mask the splice sites. We then removed these extended exon regions from the intronic regions and filtered for + strand introns for simplicity, resulting in 465.7 million sites. From these intronic regions, we extracted the coordinates of all positions starting with a GT (25.5 million sites) or ending with an AG (32.4 million sites). For each site, we extracted the spliceAi donor and acceptor probabilities and plotted their distribution in the main text figure.

TOGA2 uses a SpliceAI probability threshold of 0.02, because this cutoff achieves a very high sensitivity in capturing real splice sites (>99% for donors and acceptors from curated RefSeq transcripts), while limiting false positives. Although a threshold of 0.02 may appear low, we note that the vast majority of real splice sites have substantially higher probabilities (averages ranging from 86% to 94%). While splice sites with probabilities close to this threshold include some false positives, real exons, especially short or alternatively spliced exons, do contain such low-scoring sites. Furthermore, although the chance of incorrectly annotating an intronic site as a splice site is not zero, TOGA2 primarily relies on homology-based evidence to infer exon presence.

### Evolutionarily informed transcript selection for input annotations

As a reference-based method, TOGA2 critically relies on a comprehensive and accurate reference annotation. If conserved exons or genes are absent from the input annotation, they cannot be annotated in the query species. Likewise, incorrect or incomplete reference annotations lead to annotation errors in the query. To address this, we developed an evolutionarily informed procedure to generate high-quality input annotations by selecting conserved transcripts, while removing transcripts with non-ancestral exons and likely erroneous transcripts present in the TOGA1 input annotations.

Our input annotations are based on NCBI RefSeq annotations ^32^ for human (hg38), house mouse (mm10), cow (HLbosTau10, GCF_002263795.3), Indian elephant (HLeleMaxInd3A, GCF_024166365.1), chicken (HLgalGal7, GCF_016699485.2), Hawaiian crow (HLcorHaw3, GCF_020740725.1), black-legged kittiwake (HLrisTri2, GCF_028500815.1), zebra finch (HLtaeGut5, GCF_003957565.2), emu (HLdroNov3, GCF_036370855.1), European pond turtle (HLemyOrbi1A, GCF_028017835.1), Murray River turtle (HLemyMacqMac1, GCF_026122565.2), Pacific halibut (HLFhipSten1, GCF_022539355.2), Boeseman’s rainbowfish (HLFmelBoes1, GCF_017639745.1), turquoise killifish (HLFnotFurz3, GCF_043380555.1), three-spined stickleback (HLFgasAcul3A, GCF_964276395.1), and mandarinfish (HLFsynSple1, GCF_027744825.2) as reference species. We additionally incorporated available GENCODE annotations for human (V46) and mouse (M36) ^33^, and available Ensembl version 113 annotations for cow and chicken ^63^. Because newer annotations are available for mm39, we used liftOver to map these to the mm10 reference. For human, we incorporated MANE annotations ^105^, and for mouse, cow, and chicken we included APPRIS principal isoforms ^106^. All annotation sources and versions are listed in Supplementary Table 3.

Our evolutionarily informed transcript selection procedure is based on evaluating the integrity of a comprehensive initial transcript set across a diverse panel of query species using TOGA2. Transcripts are selected based on both coding sequence length and evolutionary conservation, measured as how often their orthologs are intact across species. Importantly, transcript length and conservation represent competing criteria. Short transcripts tend to exhibit fewer inactivating mutations and are therefore more often classified as intact, but may represent truncated or artifactual isoforms. In contrast, longer transcripts may include exons specific to the reference lineage that are not conserved and therefore often contain inactivating mutations in other species (Supplementary Figure 13).

To balance these factors, we implemented a two-step procedure. First, we constructed a comprehensive initial transcript set by combining all transcripts from the sources listed above. From this set, we removed problematic transcripts lacking a start or stop codon, containing premature stop codons, predicted to be nonsense-mediated decay targets, not divisible by three, or containing micro-introns (<20 bp) or micro-exons (<3 bp). We also rigorously removed fusion transcripts spanning the CDS of two distinct neighboring genes, as such transcripts lead to incorrect orthology assignments in TOGA2. Fusion transcripts may arise from genomic rearrangements or transcriptional readthrough events, often observed in cancer, or misannotations. Improving upon the TOGA1 filtering procedure, we identified and removed candidate fusion transcripts as those having same-strand exon overlap to different genes (Supplementary Figure 14). However, this approach also captures cases where distinct gene annotations actually represent alternative transcripts with different first coding exons but shared downstream exons of the same gene (e.g., *UGT1A*, *PCDHA*). To avoid removing these valid transcripts, we manually curated the candidates and excluded only real fusion transcripts (Supplementary Figure 15). Because MANE and APPRIS transcripts represent high-quality annotations, they were retained regardless. To remove unnecessary redundancy across the annotation sources for human, mouse, cow and chicken, we retained GENCODE/Ensembl transcripts when exon coordinates were identical to RefSeq.

In the second step, we evaluated this initial transcript set using TOGA2 (v2.0.1) across a representative panel of 101 mammals, 85 birds and 32 turtles (Supplementary Table 3). For each isoform, we determined the number of species for which the ortholog was classified as fully intact, intact, partially intact, uncertain loss, or lost. We also determined the average number of inactivating mutations and the CDS length. We then applied an iterative selection procedure to identify representative transcripts. For each gene, a “winner” transcript was selected based on a penalty function combining CDS length and average number of inactivating mutations: penalty = 0.5 × CDS length - 0.9 × average number of mutations. The parameters were empirically determined to preferentially select human MANE transcripts as the winner. Iterating over all remaining transcripts of the focal gene, we included additional transcripts if they were classified as fully intact in at least as many species as the winner and had a CDS length of at least 60% of the winner, which avoids overly short isoforms. MANE and APPRIS transcripts were always retained.

Manual inspection indicated that this procedure effectively removes truncated and lineage-specific transcripts while retaining or even adding conserved genes or exons (Supplementary Figures 12-13). However, we observed that some transcripts supported by BUSCO gene sets or independent RefSeq annotations in query species were excluded. To address this, we performed an additional refinement step by adding transcripts whose orthologs were classified as missing in a compleasm analysis (mammalia_odb10,aves_odb10 and sauropsida_odb12 datasets) of the filtered transcript set but present in the initial set. Similarly, for the four mammalian references, we added transcripts whose orthologs are independently supported by curated RefSeq (NM_) annotations in human, mouse, cow, and rat (GCF_015227675.2-RS_2023_06), using rat instead of elephant due to the limited number of curated RefSeq transcripts in elephant. Likewise, for the five bird references, we added transcripts whose orthologs match curated RefSeq transcripts in the other four reference species. For the two turtles, we added transcripts that match RefSeq transcripts from eight RefSeq annotated turtles and the respective other reference.

Transcripts lacking annotated UTRs in the reference can interfere with orthology inference, as alignments between reference and query that overlap UTRs will be misclassified as intergenic sequence. Furthermore, TOGA2 can annotate UTRs in the query species. For these reasons, we preferentially selected transcripts with annotated UTRs (Supplementary Figure 16).

Because evolutionarily informed transcript selection removed relatively few transcripts in halibut and because TOGA2 annotated more genes as paralogs in the distantly related percomorph fish, we did not apply this filtering procedure to the other four fish reference species and retained only the filtering step for valid transcripts.

All input annotations are available at https://github.com/hillerlab/TOGA2/tree/main/reference_annotation. The code for the evolutionarily informed transcript selection procedure is available as the TOGA2 module “prepare-input”, along with a simplified framework for filtering input annotations based on the initial removal of problematic transcripts.

### UTR annotation benchmarks

We used the four mammalian references (human, mouse, cow and elephant) with the following taxonomically diverse placental mammal assemblies *Macaca nemestrina* (HLmacNeme2A=GCF_043159975.1), *Rattus norvegicus* (HLratNor8=GCF_036323735.1), *Mus musculus*(mm10=GCF_000001635.20), *Oryctolagus cuniculus* (HLoryCuni5=GCF_964237555.1), *Bos taurus* (HLbosTau10=GCF_002263795.3), *Ovis aries* (HLoviAri6=GCF_016772045.2), *Equus caballus* (HLequCaba4=GCF_041296265.1), *Canis lupus baileyi* (HLcanLupBai8A=GCF_048164855.1), *Neofelis nebulosa* (HLneoNeb2=GCF_028018385.1), *Tenrec ecaudatus* (HLtenEcau2A=GCF_050624435.1), *Elephas maximus indicus* (HLeleMaxInd3A=GCF_024166365.1), and *Choloepus didactylus* (HLchoDid2=GCF_015220235.1). For each query assembly, we used the coding transcripts from the NCBI RefSeq assembly as the benchmark annotation (ground truth). For mouse, we added coding transcripts from the GenCode V25 annotation, which we obtained from the UCSC genome browser table wgEncodeGencodeCompVM25. For each fully-intact TOGA2 transcript, we then determined which transcripts in the ground truth annotation have an identical N- or C-terminal coding exon, as UTR comparisons make no sense if the terminal CDS does not match. We computed the strand-specific overlap between the TOGA2 predicted UTR and the UTR of the matching benchmark transcripts, using the UCSC tool overlapSelect with parameters - statsOutput -strand to obtain the similarity value, defined as (2*overlap in bp) / (TOGA2 UTR length + annotated UTR length). In case of several matching transcripts, the maximum similarity was reported. TOGA2 or benchmark transcripts that lack a 5’ or 3’ UTR were ignored.

To compare annotated transcription start sites (TSSs), defined as the first 5’ UTR base, to CAGE data, we obtained for mouse (mm10 assembly) the FANTOM5 CAGE peaks (clusters of CAGE tags identified by the decomposition-based peak identification algorithm) and annotated TSSs from the UCSC genome browser download server (bigBed file https://hgdownload.soe.ucsc.edu/gbdb/mm10/fantom5/mm10.cage_peak.bb). Fantom5 provides 164,672 peaks with a median length of 15 bp (average 20.7 bp). We compared the strand-specific overlap between FANTOM5 CAGE peaks and CAGE TSSs with TOGA2-inferred TSSs (considering fully intact transcripts as above), RefSeq-annotated TSSs (using curated and curated+predicted coding transcripts obtained from the UCSC genome browser tables ncbiRefSeqCurated and ncbiRefSeq), and GENCODE-annotated TSSs (using the basic and comprehensive coding transcript annotations obtained from the tables wgEncodeGencodeBasicVM25 and wgEncodeGencodeCompVM25). We analyzed all transcripts located on mouse chromosomes 1-19 and X. For the annotated TSSs that overlapped CAGE peaks, we also calculated the distance of the annotated TSS to the closest CAGE TSS with bedtools closest (parameters -s -t first) ^107^. Negative distances refer to annotated TSS that are upstream of the CAGE TSS. Boxplots of the distance distributions are shown in the main text figure. Finally, to evaluate the effect of reference input annotation quality on the TOGA2 TSS prediction accuracy, we used cow as a reference and benchmarked TOGA2-annotated TSSs derived from curated RefSeq cow input transcripts (NM_ transcripts).

### Annotation completeness benchmarks

For the annotation completeness comparison, we collected all RefSeq assemblies from the focal clades (mammals, birds, turtles, and percomorph fishes) that were annotated by RefSeq after 2020 and for which TOGA2 was also run (note that GenBank assemblies are not annotated by NCBI). Because GenBank assemblies are typically released earlier than their corresponding RefSeq versions, and many TOGA2 runs were performed prior to the availability of RefSeq assemblies, we only used genomes where TOGA2 had been run on the corresponding RefSeq assembly, in order to avoid potential discrepancies between GenBank and RefSeq genome versions. No genomes were excluded from the analysis for any other reason.

We downloaded RefSeq annotations (GCF*protein.faa.gz files) from NCBI. For Ensembl, we downloaded protein annotations (pep.fa.gz files) for the subset of NCBI test species with a matching genome accession. For all these assemblies, we ran Annevo ^28^, using the ANNEVO_Mammalia.pt model for mammals and the ANNEVO_Vertebrate_other.pt model for birds, turtles and fishes. Protein sequences were generated from annotation files using genePredToProt from the UCSC kent source code (https://github.com/ucscGenomeBrowser/kent). For TOGA2, we used the protein.fa.gz files generated by the method.

For each protein FASTA file, we ran compleasm ^67^ using the mammalia_odb12 dataset of near universally conserved genes for mammals, aves_odb12 for birds, sauropsida_odb12 for turtles, and actinopterygii_odb12 for fishes ^68^. For combined TOGA2 analyses, we computed the union of completely detected genes across single-reference TOGA2 annotations. All results, including RefSeq and Ensembl protein fasta files, their annotation timestamps, and the number of completely detected BUSCO genes for all benchmarked annotations, are provided in Supplementary Tables 7-10.

### Mouse GENCODE benchmark

To identify TOGA2-annotated coding exons absent from GENCODE annotations, we first generated a filtered set of TOGA2 exons using human, cow, and elephant as reference species. We considered only exons from fully intact (FI), intact (I), partially intact (PI), or uncertain loss (UL) projections. We further required that internal exons have both splice sites supported by SpliceAI (probability ≥0.02). For terminal exons, we required SpliceAI support for their sole splice site. Finally, we retained only exons without any (masked or unmasked) inactivating mutations.

Using this filtered exon set, we identified exons for which at least one splice site was not annotated in GENCODE, which avoids differences caused only by terminal exon length variation. To exclude frequent short distance 3 bp alternative donor or acceptor site variation ^108,109^, we further required that the novel splice site was located at least 5 bp away from the nearest GENCODE splice site. For these exons, we determined whether they were supported by one, two, or all three TOGA2 reference species, whether RefSeq annotates them with identical splice sites, and whether they were supported by RNA-seq data.

### RefSeq locus gene identifiers

Inspired by ^80^, we investigated how often TOGA2 assigns informative gene symbols to RefSeq LOC genes. For the same 70 mammalian species used in the compleasm comparison, we first extracted all RefSeq transcripts from the GCF*genomic.gff.gz files that are annotated with gene symbols corresponding to locus identifiers matching the pattern “LOC[0-9]+”. In addition, we downloaded the human and mouse RefSeq annotation from the UCSC genome browser table ncbiRefSeq. Only coding transcripts were retained. For each of the 70 query species, we used the four TOGA2 annotations and selected transcripts classified as fully intact, intact, partially intact, or uncertain loss. We removed paralogous transcripts as well as transcripts with uninformative reference gene symbols (LOC[0-9]+, ENSG/ENSMUS/ENSBTAG, KIAA*, Gm[0-9]+, MGC[0-9]+, BC/AI/AW[0-9]+, *Rik, C*orf*). TOGA2 retrogene annotations were retained, as retrogenes are frequently annotated as RefSeq LOC genes.

To enable comparison at the coding level, we removed annotated UTRs from both the RefSeq LOC transcripts and the filtered TOGA2 transcripts, retaining only coding exons. We then used overlapSelect from the UCSC kent source code (https://github.com/ucscGenomeBrowser/kent) with parameters -strand -statsOutput to match RefSeq and TOGA2 transcripts, requiring that either inOverlap or selectOverlap exceed 0.3. This threshold captures fragmented RefSeq annotations overlapping longer TOGA2 transcripts and vice versa. The main text figure shows the total number of RefSeq LOC genes and the subset for which TOGA2 assigns informative gene symbols.

### Multi-species orthogroups (toga2orthogroups)

To infer orthogroups across multiple species from pairwise TOGA2 orthology assignments, we modeled orthology relationships between reference and query as a graph. The Union-Find algorithm is then used to efficiently extract connected components representing orthogroups. In the graph, each reference gene is represented as a node. Whenever two reference genes are co-orthologs to the same query gene, they are merged into the same component. Union-Find handles this incrementally through path compression and rank-balanced union operations, resulting in effectively linear time complexity, which enables genome-wide analyses across hundreds of species. Optionally, toga2orthogroups allows to incorporate a gene family database such as PANTHER ^86^ to consolidate orthogroups by adding family-level edges between reference genes prior to component extraction. The output is readily formatted for direct input into gene family evolution tools such as CAFE5 ^87^.

After constructing orthogroups, toga2orthogroups computes a diagnostic report to identify species contributing to unusually large orthogroups using two metrics. First, a species may exhibit globally abnormal orthogroup sizes, potentially caused by assembly issues, such as a high proportion of missing or inactivated genes, or by inflated one-to-many orthology relationships. To identify such species, toga2orthogroups computes, for each orthogroup, a per-species Z-score defined as (N−μ)/σ, where N is the orthogroup size of the species, and μ and σ are the mean and standard deviation of orthogroup sizes across species, respectively. For every species, the number of orthogroups with an outlier Z-score, defined as Z≥T or Z≤−T, where T is a user-defined threshold (default 3), is counted. From these per-species outlier counts, toga2orthogroups computes a second per-species Z-score, termed FamZ. High FamZ values indicate consistently abnormally small or large orthogroup sizes across many gene families.

Second, orthogroup inflation can arise when incorrect orthology relationships between query and reference genes merge otherwise distinct orthogroups into the same connected component. To identify species contributing to such inflation, toga2orthogroups computes, for each orthogroup and species, the orthogroup size (N) and the number of non-1:1 orthology relationships (O) to reference genes within that orthogroup. Specifically, for each query gene with k orthologs in the reference, we add k-1 to O (a 1:1 ortholog would contribute 0; a 3:1 orthology relationship would contribute 2). For each orthogroup, the ratio O/N is converted into a per-species Z-score by considering the distribution of O/N values across all species. Across all orthogroups, positive Z-scores are summed for each species. From these summed values, toga2orthogroups computes a second per-species Z-score, termed OrthoZ. High OrthoZ values (default value 3) indicate that a species systematically contributes to orthogroup inflation through excessive orthology relationships. Both metrics enable users to identify problematic assemblies in an initial pass and exclude them prior to final orthogroup inference and downstream analyses.

### Synteny of orthologs (toga2agora)

The module toga2agora extracts synteny information of TOGA2-inferred orthologous genes generated using a single reference species and produces output files compatible with AGORA for ancestral gene order reconstruction ^9^. The module requires as input a phylogenetic tree in Newick format describing the relationships between the reference and query species and a path to reference annotation and TOGA2 results for all queries listed in the tree.

First, for each query species present in the tree, toga2agora selects 1:1 orthologs classified with selected loss statuses (FI, I, PI, and UL by default) and generates ordered gene lists as tab-separated five-column files containing query chromosome, start, stop, orientation, and gene identifier information. Second, toga2agora generates orthology group files for each ancestral node, listing genes present as 1:1 orthologs in all descendant child nodes.

These files can subsequently be used by AGORA in basic or generic mode. In basic mode, AGORA performs pairwise genome comparisons to identify conserved gene adjacencies, constructs ancestral adjacency graphs, and linearizes these graphs into contiguous ancestral regions (CARs). In generic mode, AGORA evaluates multiple parameter combinations and selects the parameter set producing the highest G50 value independently for each ancestor, with G50 being defined as the CAR length at which 50% of the total reconstructed ancestral genome length (measured in gene units) is contained within CARs of equal or greater size.

### TOGA2 for transcript quantification (toga2kbpython)

The module toga2kbpython uses kb-python ^110^ to build an index of TOGA2-annotated transcripts (with or without UTR predictions). The genome sequence is included as a decoy to increase the accuracy of transcript quantifications by avoiding false RNA-seq read assignments. For quantification of coding gene expression and cross-species expression comparisons, we recommend using the annotations without UTRs, as CDS can be annotated more reliably and CDS vary less between species than UTRs. In this case, we mask the coding sequences in the genomic decoy to enable assigning reads that partially overlap CDS and UTR to transcripts. Retrogene candidates are excluded from the annotation because they are often not expressed and may lead to underestimation of parental gene expression due to ambiguous read mapping. The generated index is then used by kb-python to quantify RNA-seq data by pseudomapping. Transcript-level expression values (read counts or transcripts per million) are summed across all (including fragmented) transcripts of the same gene to obtain gene-level expression values.

Duplicated genes with 1:many or many:many orthology relationships are merged by toga2kbpython into a single gene unit, summing the expression values across all gene copies. Large gene duplication units (e.g., comprising more than 10 genes) are excluded, as they typically represent large gene families for which annotation and expression estimates are less accurate. For genes duplicated in the reference (many:1 orthology relationships), toga2kbpython assigns merged gene names.

The final result of toga2kbpython is a gene-level expression matrix. This matrix can be further filtered, for example to keep only 1:1 orthologs, and can be used as input for differential expression analyses with DESeq2 ^89^.

### Assembly quality metrics using an updated ancestral gene set (toga2stats)

We previously defined a set of 18,430 genes that likely already existed in the placental mammal ancestor ^45^. This set comprised genes present in the human GENCODE v38 annotation for which TOGA1 identified an ortholog with an intact reading frame in at least one afrotherian and at least one xenarthran genome. Because the genome assembly quality of previously available afrotherian and xenarthran genomes was limited, we used rather permissive criteria (presence in at least one species per clade).

Since genome assembly quality has substantially improved, we updated the placental mammal ancestral gene set with the goal of obtaining a more robust, high-confidence gene set. To this end, we used TOGA2 annotations for the following six high-quality afrotherian assemblies: *Elephas maximus indicus* (GCF_024166365.1), *Loxodonta africana* (GCA_030014295.1), *Heterohyrax brucei* (GCA_028571685.1), *Dugong dugon* (GCA_030035585.1), *Trichechus inunguis* (GCA_046562895.1), and *Rhynchocyon petersi* (GCA_043290085.1), as well as the following high-quality xenarthran assemblies: *Bradypus torquatus* (GCA_963992745.1), *Choloepus didactylus* (GCF_015220235.1), *Dasypus novemcinctus* (GCF_030445035.1), *Tamandua tetradactyla* (GCA_023851605.1), *Myrmecophaga tridactyla* (DNAZoo assembly), and *Tolypeutes matacus* (GCA_026826555.1).

For afrotherians, we retained genes that were fully intact in at least three species or intact in at least five of the six species. The same criteria were applied to xenarthrans. We then retained only genes that passed these filters in both afrotherians and xenarthrans, yielding 15,713 genes. Finally, we excluded genes belonging to large, rapidly-evolving gene families (olfactory receptors, zinc finger proteins, and keratin-associated proteins), as well as genes that only have a LOC or ENSG identifier without an assigned gene symbol. This resulted in a final set of 15,196 ancestral placental mammal genes.

The module toga2stats then uses this gene set and TOGA2’s gene classification to benchmark the quality of one or more query genomes by quantifying the proportion of ancestral genes with missing sequence or inactivating mutations.

## Acknowledgment

We thank Graham Hughes for advice on IQTree2, Giulio Formenti for manuscript feedback, Hiram Clawson for providing TOGA2 tracks in the UCSC genome browser, and Christoph Sinai for excellent HPC support. This project has received funding from the European Research Council (ERC) under the European Union’s Horizon 2020 research and innovation programme (grant agreement No. 101118919), the National Institute On Aging of the National Institutes of Health under Award Number U19AG023122, the German Research Foundation (HI1423/5-1 and HI1423/6-1), the Leibniz Association’s Competition Procedure (K419/2021), the LOEWE-Centre for Translational Biodiversity Genomics (TBG) funded by the Hessen State Ministry of Higher Education, Research and the Arts (LOEWE/1/10/519/03/03.001(0014)/52).

## Data and Code Availability

All data generated in this study, including softmasked genomes, repeat libraries, whole genome alignments, gene annotations with UTR predictions, orthologs, gene loss and duplication events, inactivating mutations, processed pseudogenes and retrogene candidates, nucleotide and protein sequences and alignments, and clade-wide sets of 1:1 orthologs, are available for download at https://genome.senckenberg.de/download/TOGA2/. The TOGA2 source code and all associated tools are on github https://github.com/hillerlab/TOGA2. We also provide genome browsers showing all annotations at https://genome.senckenberg.de/.

## Supplemental Data

**Supplementary Figure 1:**
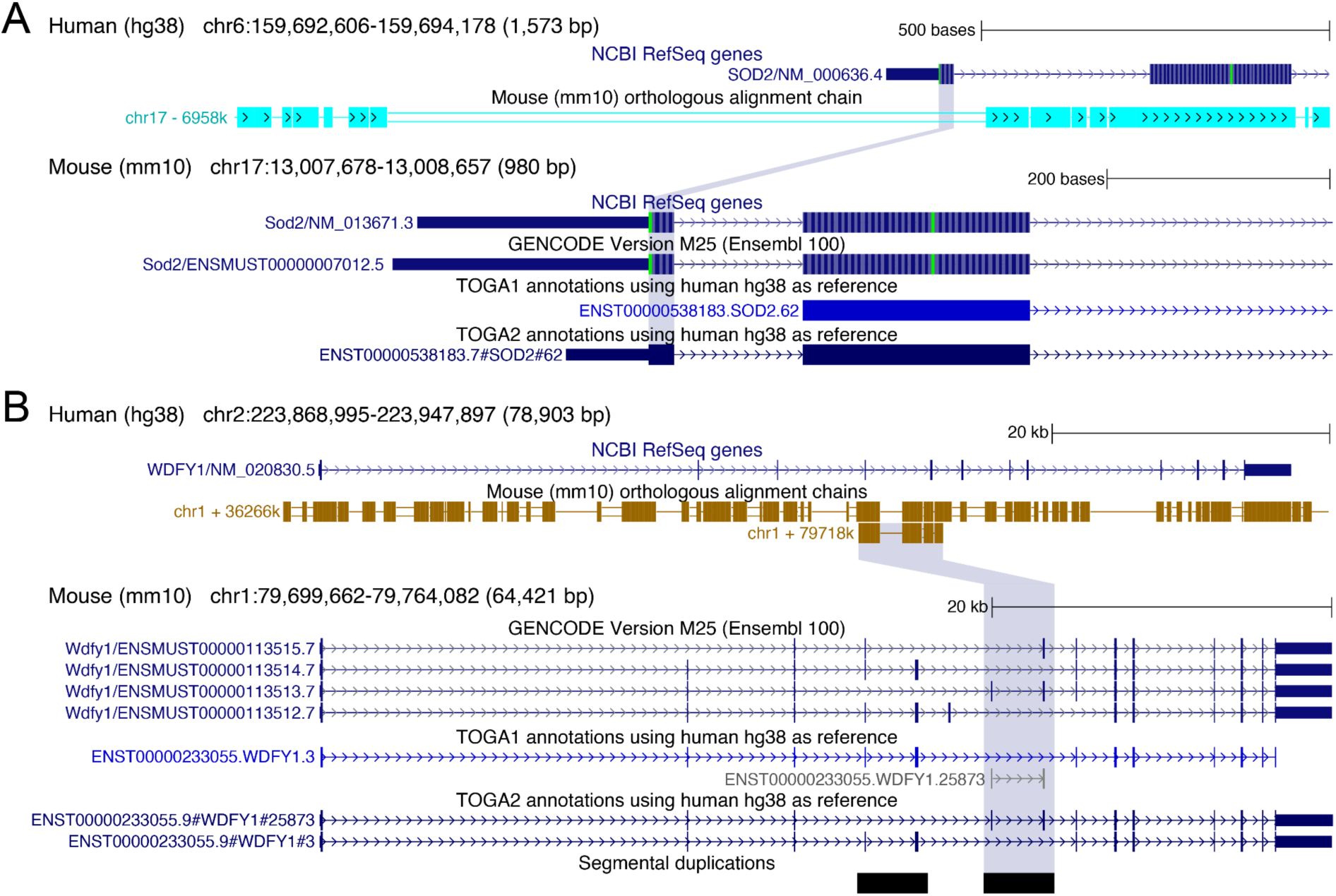
Search space extension in TOGA2 enables annotation of additional exons. Examples illustrating how TOGA2’s ability to extend the search space when exons of a reference transcript are not covered by the orthologous alignment chain allows recovery of additional exons. (A) Coding exon 1 of *SOD2* has no detectable nucleotide alignment in the orthologous chain (top). As a result, TOGA1 fails to annotate exon 1. In contrast, TOGA2 recognizes that exon 1 is missing from the alignment and extends the search space, allowing CESAR2 to recover the exon at the protein alignment level, leading to the inclusion of exon 1 in the annotated transcript. (B) While chain ID 3 (top chain) covers the entire *WDFY1* gene locus, chain ID 25873 (blue highlight) spans only two exons but corresponds to an orthologous segment of the *WDFY1* gene in mouse. Both TOGA1 and TOGA2 recognize this chain as an orthologous fragment of the gene. However, whereas TOGA1 can annotate only the aligned exons, resulting in a highly-truncated transcript that is classified as missing, TOGA2 estimates the upstream and downstream portions of the transcript not covered by the chain alignment and adds these regions to the query search space. Running CESAR2 in multi exon mode then recovers the complete transcript. The blue-highlighted chain (25873) represents a segmental duplication in mouse (mm10, also present in mm39) that includes two exons that appear to be spliced in a mutually exclusive manner. This putative tandem exon duplication complicates query gene inference because it results in two projections of the same transcript in the same query locus, but chain-supported coding exons do not overlap. Because TOGA2 normally assesses same-strand coding exon overlap only for exons directly supported by the alignment chain, this situation would incorrectly lead to inference of two separate query genes. To avoid this problem, TOGA2 makes an exception for projections of the same transcript ID and assesses overlap of all coding exons, including exons annotated by search space extension, thereby correctly inferring a single query gene.

**Supplementary Figure 2:**
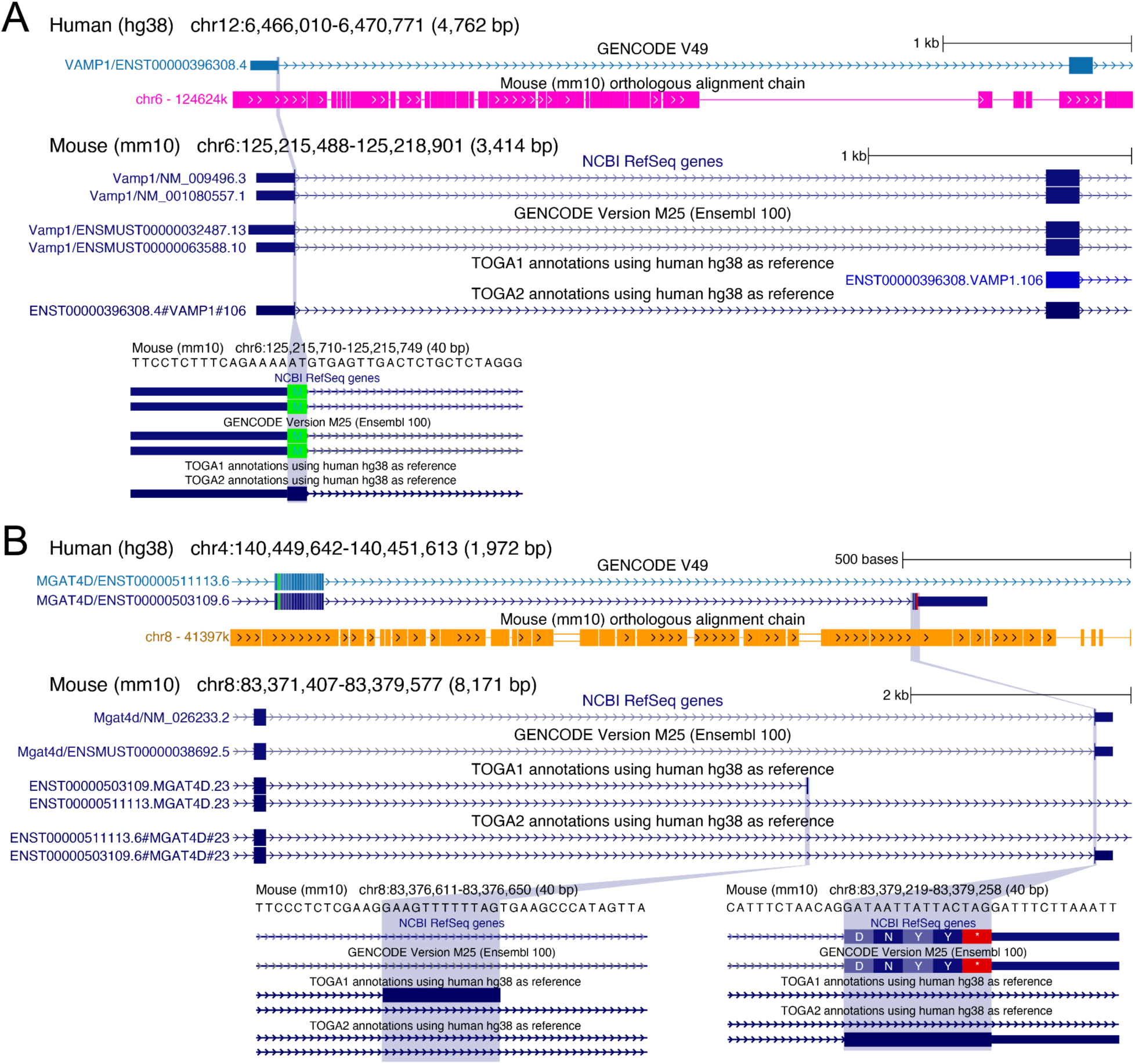
Single-exon mode enables TOGA2 to correctly annotate microexons. TOGA1 simultaneously infers the positions and boundaries of all exons across the entire orthologous query locus using CESAR2 in multi-exon mode. While locating normal-sized exons is generally unambiguous, identifying very small coding exons is challenging because they provide limited sequence for alignment. TOGA2 leverages the fact that most exons, including microexons, are flanked by intronic alignments, which define their location in the query genome. Exon-level orthology allows TOGA2 to run CESAR in single-exon mode, enabling precise annotation of such exons, as illustrated in the main text and this figure. (A) In *VAMP1*, the start codon is split across the boundaries of exons 1 and 2, such that exon 1 contains only two coding nucleotides (AT). TOGA1 fails to annotate this exon, resulting in an incomplete transcript, whereas TOGA2 correctly captures it. (B) *MGAT4D* contains a 12 bp terminal coding exon. TOGA1 misidentifies this exon approximately 2.6 kb upstream of its true location, whereas TOGA2 precisely annotates it despite substantial sequence divergence (human protein DVY, mouse protein DNYY).

**Supplementary Figure 3:**
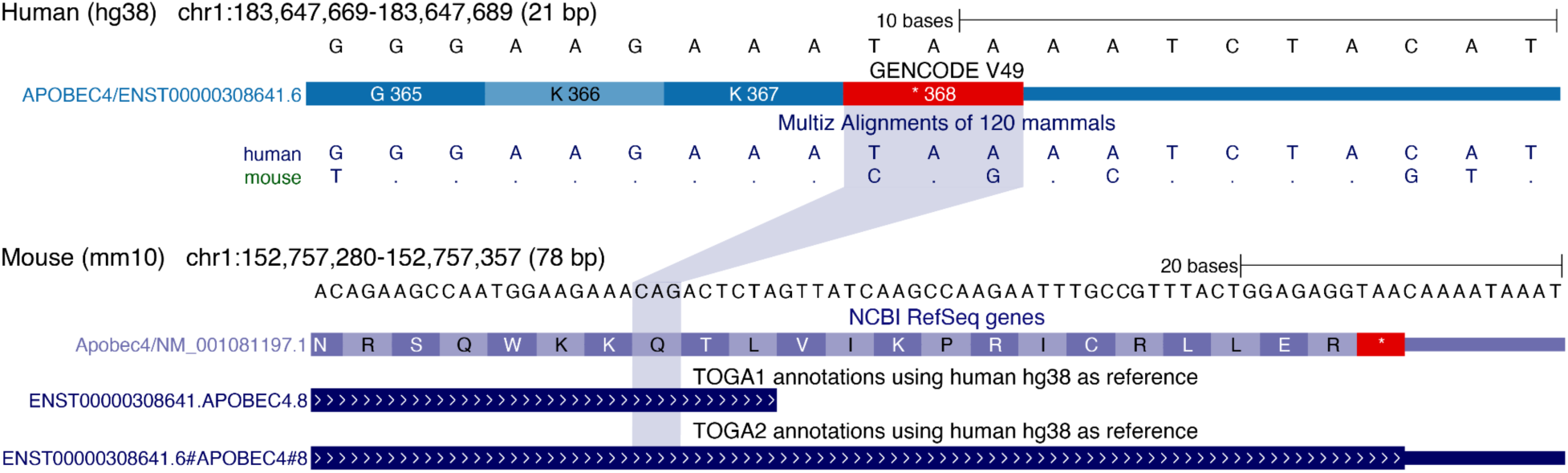
Capturing mutated stop codons improves annotation accuracy of terminal exons. The *APOBEC4* mouse ortholog has mutations in the TAA stop codon, resulting in a CAG sense codon and a 36 bp C-terminal protein extension. Whereas TOGA1 fails to account for this change, TOGA2 detects the mutated stop codon and searches for a downstream in-frame stop codon, which corrects the annotation of this terminal exon.

**Supplementary Figure 4:**
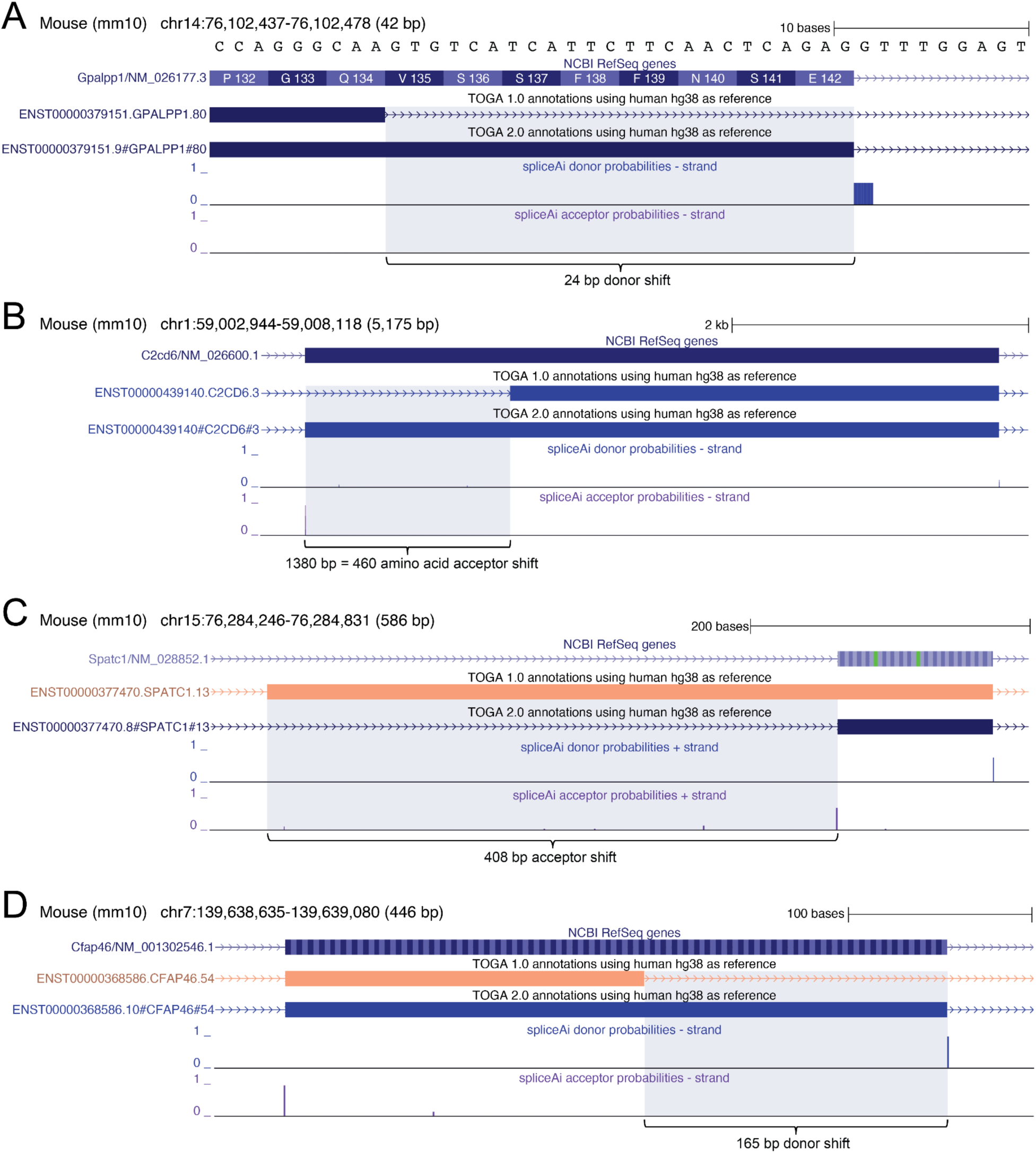
TOGA2 uses deep learning–based splice site predictions to improve exon annotation accuracy. (A) Coding exon 4 of *GPALPP1* exhibits a 24 bp donor splice site shift. While the donor predicted by TOGA1 matches the GTG consensus, SpliceAI correctly predicts the donor 24 bp downstream. TOGA2 uses this information to annotate the exon correctly. (B) Large acceptor shift in the penultimate exon of *C2CD6*. The acceptor has shifted by 1380 bp, resulting in a mouse exon that is longer by 460 amino acids. Such large splice site shifts cannot be predicted by CESAR2 and consequently TOGA1 mis-annotates this exon. In contrast, TOGA2 additionally uses SpliceAI predictions to correctly annotate the exon. (C–D) Examples illustrating that correct splice site identification not only improves exon annotation but also transcript classification. (C) Exon 3 of *SPATC1* exhibits a 408 bp acceptor shift. TOGA1 predicts a CA acceptor site, which is recognized as a splice site mutation, resulting in the classification of this transcript as uncertain loss. TOGA2 captures the splice site shift, resulting in a canonical CAG acceptor and an intact transcript classification. (D) Exon 30 of *CFAP46* exhibits a 165 bp donor shift. TOGA1 predicts a GA donor, resulting in an uncertain loss classification. Using SpliceAI predictions, TOGA2 identifies the correct GTG donor, yielding an intact transcript classification.

**Supplementary Figure 5:**
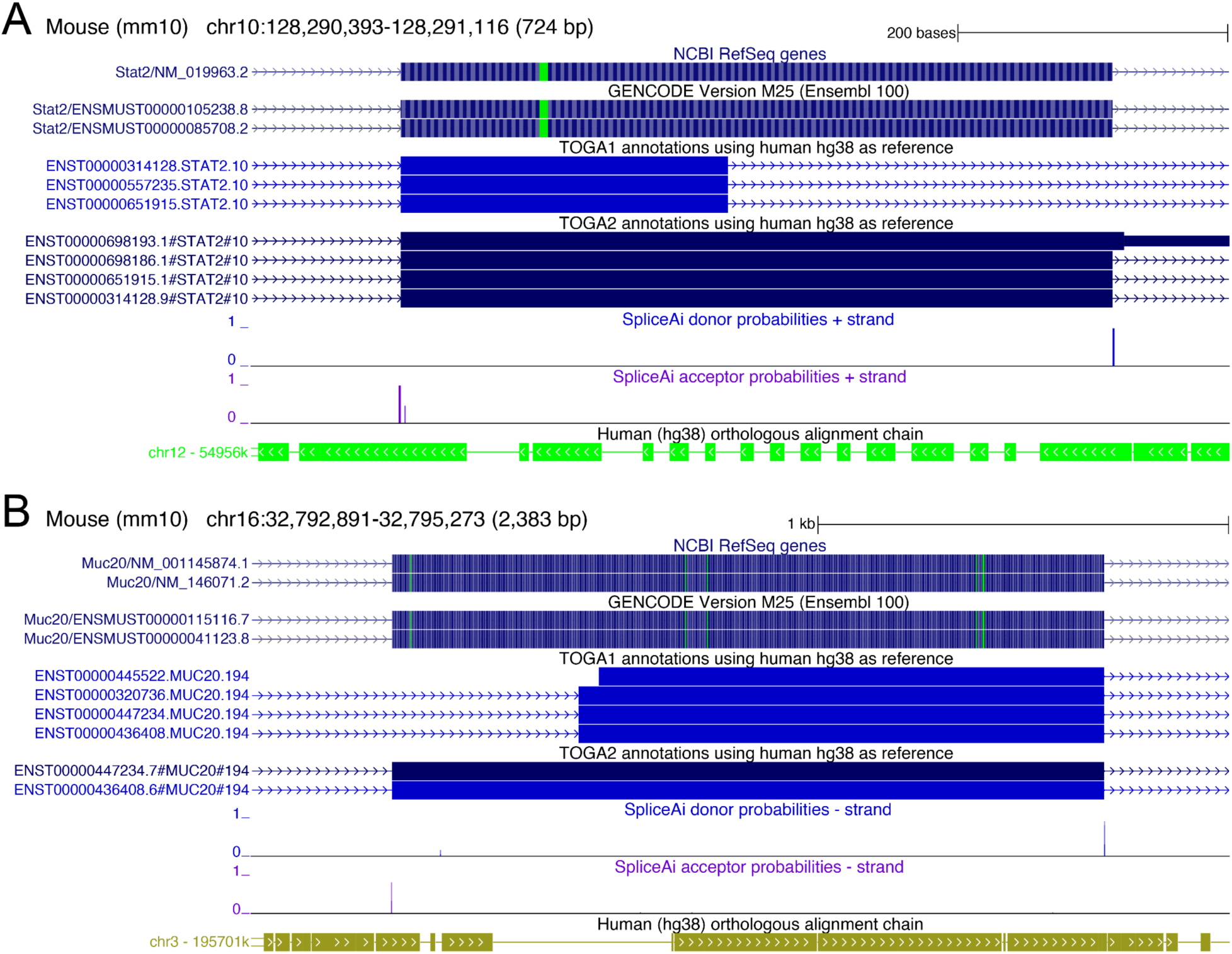
Deep learning-based splice site predictions enable TOGA2 to correctly annotate exons with large length variation. (A) *STAT2* coding exon 22 exhibits numerous insertions relative to human, as shown by the orthologous alignment chain, resulting in a substantially longer exon in mouse (527 bp vs. 311 bp in human). CESAR2’s HMM fails to capture all insertions and instead predicts an upstream donor site, causing TOGA1 to annotate a markedly shorter exon. In contrast, TOGA2 leverages SpliceAI probabilities and shifts the donor site by 285 bp, yielding the correct exon annotation. (B) *MUC20* coding exon 2 contains both large insertions and deletions, resulting in substantial exon length variation (mouse 1734 bp vs. human 1893 bp). CESAR2’s HMM does not fully capture these indels and predicts a downstream acceptor site, leading TOGA1 to annotate a substantially shorter exon. TOGA2 shifts the acceptor 454 bp upstream to the splice site best supported by SpliceAI, resulting in the correct exon annotation.

**Supplementary Figure 6:**
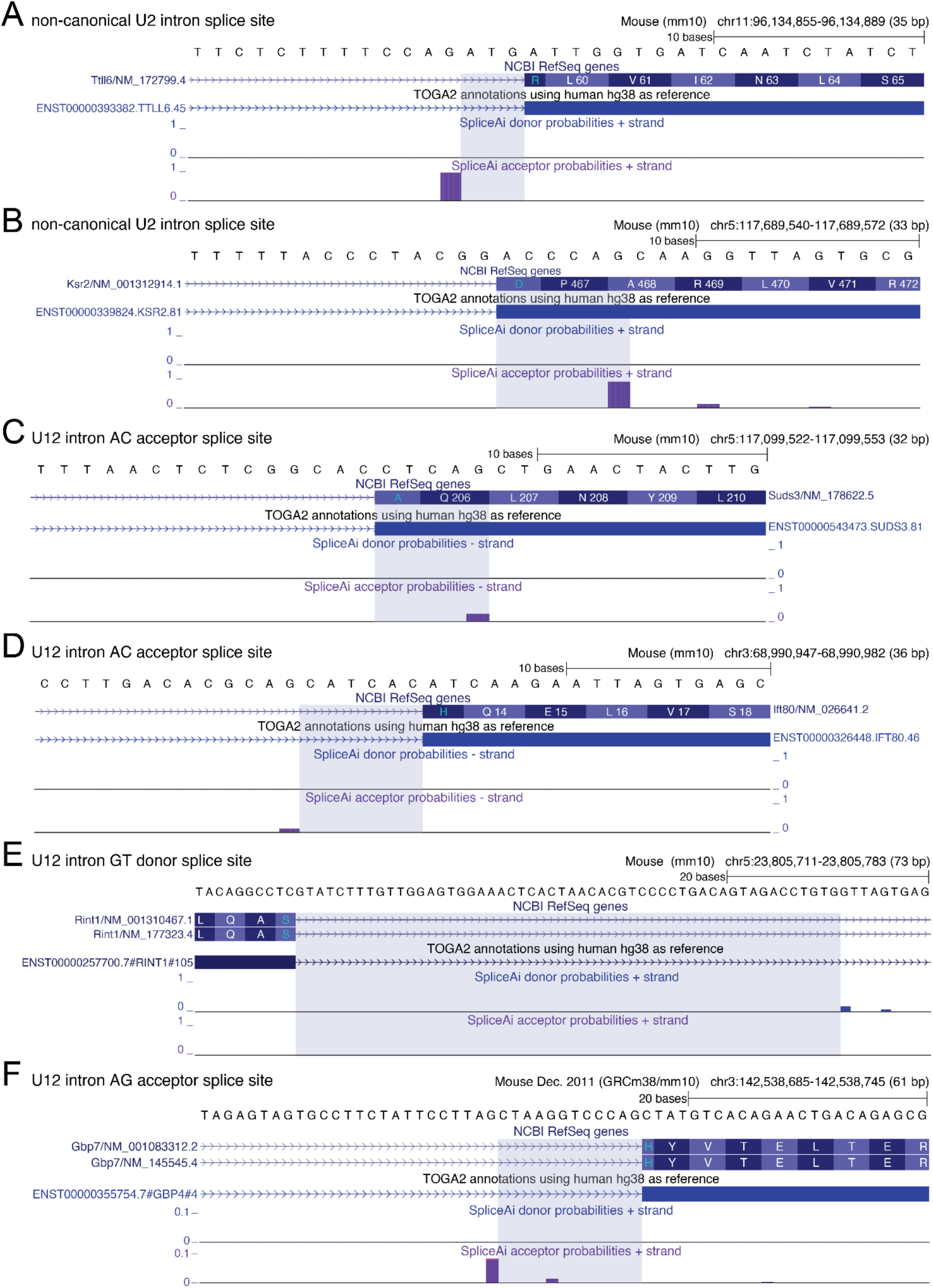
SpliceAI cannot accurately predict non-canonical U2 or U12 splice sites. U12 introns and particularly U2 introns with non-canonical donor (non-GT/GC) or acceptor (non-AG) sites are very rare. The examples presented in this figure show that SpliceAI fails to correctly predict such splice sites. While SpliceAI often predicts nearby but incorrect splice sites, suggesting that it recognizes the presence of an exon, it fails to pinpoint the correct splice site. (A, B) Two examples of non-canonical U2 splice sites with a TG (A) and a GG acceptor (B). (C, D) Two examples of U12 introns with AC acceptor sites. (E, F) More than half of vertebrate U12 introns have GT–AG dinucleotides ^90^. These examples show that SpliceAI fails to predict U12 GT donor sites (E) and U12 AG acceptor sites (F).

**Supplementary Figure 7:**
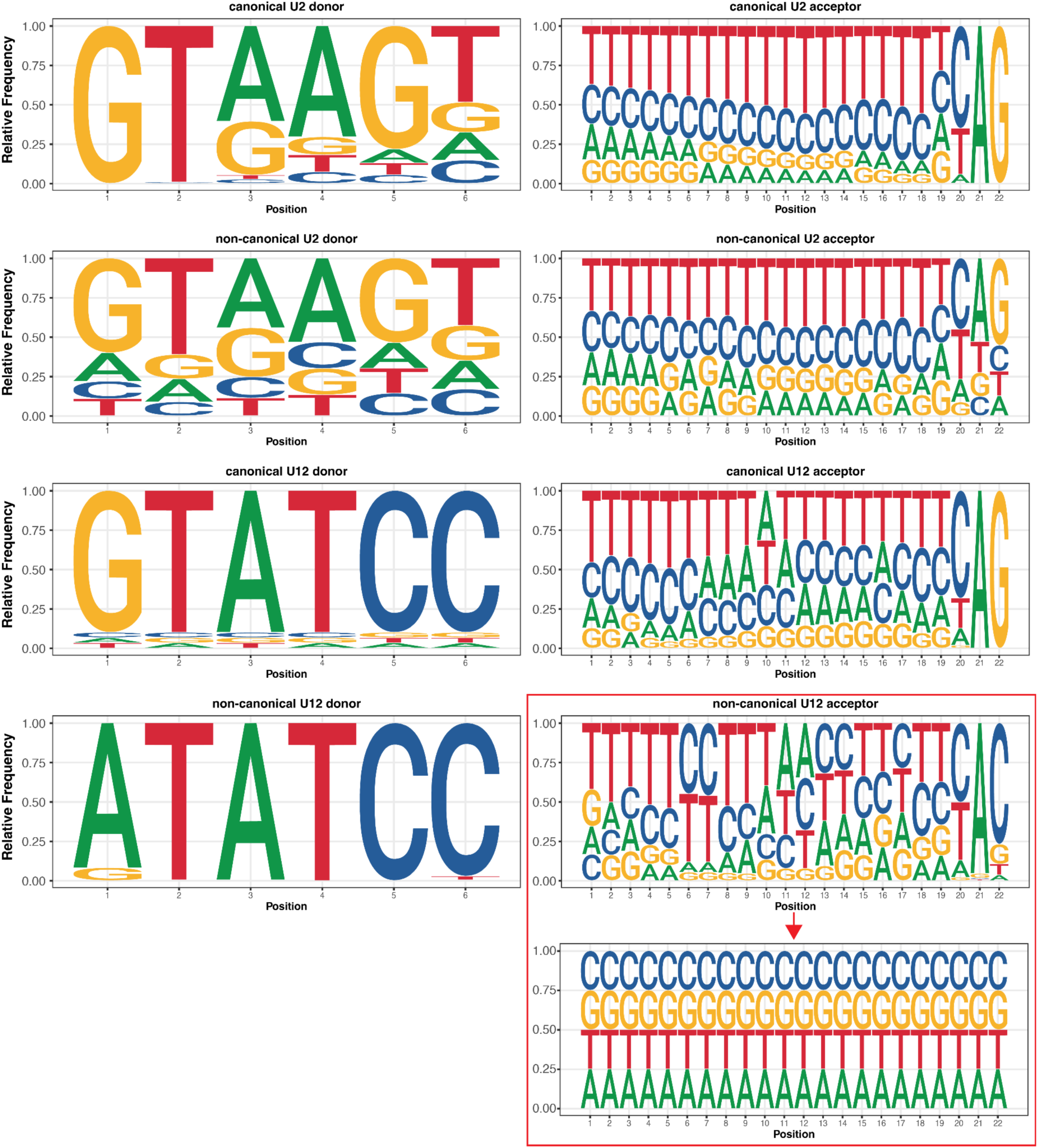
Splice site profiles trained for human introns. Sequence logos show the relative frequency at each position for the four donor and four acceptor types. The previously used profile for non-canonical U12 acceptors (top logo in the red box) contains very low information content but can still lead to incorrect splice site shifts in some cases. Tests using human as reference and mouse (mm10) as query showed that the accuracy of non-canonical U12 acceptor annotation is improved when sequence constraints are relaxed to a uniform frequency distribution (bottom logo in the red box).

**Supplementary Figure 8:**
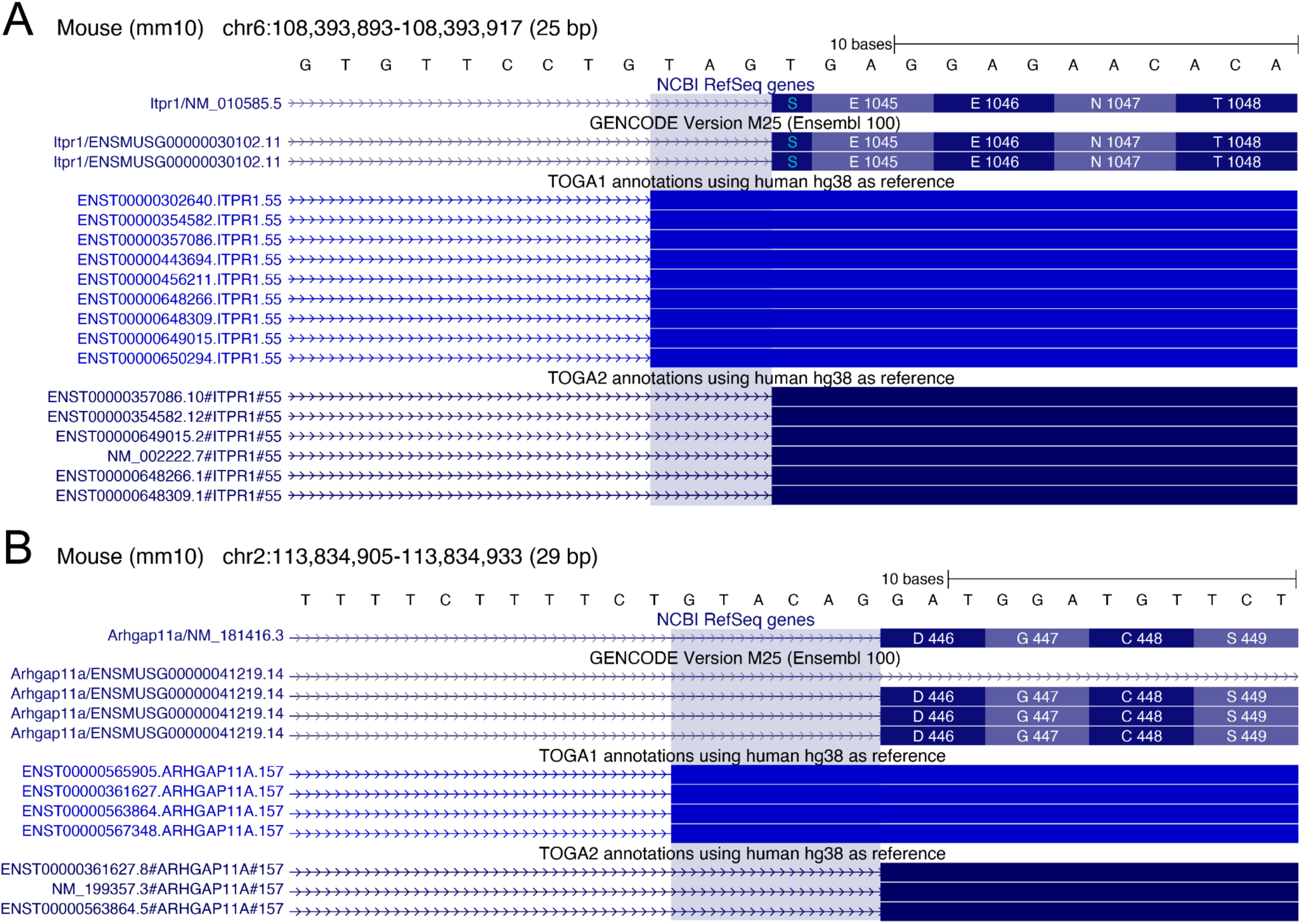
Misclassified U12 introns led to incorrect splice site annotations in TOGA1. (A) Intron 24 of *ITPR1* is a non-canonical U2 intron with GA–AG splice sites. Due to the non-canonical GA donor, this U2 intron was previously misclassified as U12 in U12DB ^111^, causing TOGA1 to apply U12 splice site profiles, which led to the misprediction of the TAG acceptor. TOGA2 instead relies on intron classifications generated by intronIC ^90^, which correctly identifies this intron as a non-canonical U2 intron. TOGA2 applies the canonical U2 acceptor profile to the canonical AG acceptor site, enabling the correct annotation of this exon. By applying the non-canonical U2 donor splice site profile, TOGA2 also correctly annotates the non-canonical GA donor upstream (not shown). (B) Similar to (A), intron 10 of *ARHGAP11A* is misclassified as U12 in U12DB despite having canonical GT-AG splice sites. As a result, TOGA1 applies U12 splice site profiles and mispredicts the TAG acceptor. IntronIC correctly classifies this intron as U2, allowing TOGA2 to accurately annotate both splice sites.

**Supplementary Figure 9:**
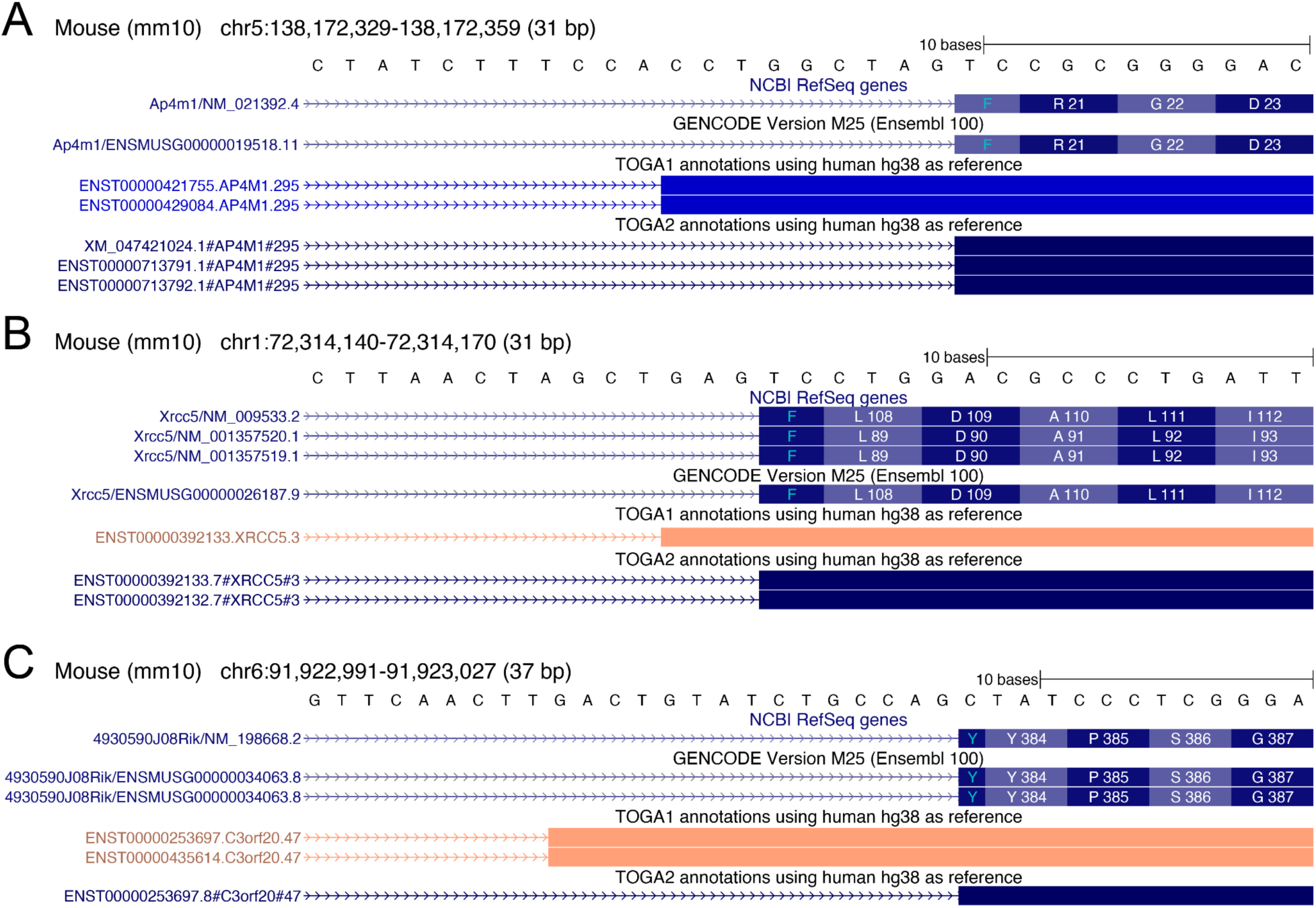
TOGA2 enhances U12 splice site annotation accuracy. UCSC Genome Browser screenshots comparing RefSeq, GENCODE, TOGA1, and TOGA2 annotations, illustrating the impact of updated U12 splice site profiles and the distinction between GT-AG U12 introns and U12 introns with different dinucleotides. (A) *AP4M1* intron 1 is a GT-AG U12 intron. TOGA1 mispredicts the acceptor site 9 bp upstream, whereas TOGA2 correctly annotates the TAG acceptor. (B) *XRCC5* intron 3 is a GT-AG U12 intron. TOGA1 mispredicts the acceptor 3 bp upstream, annotating the neighboring exons as T|GT…GCT|GA. This introduces a TGA stop codon across the exon junction and results in an incorrect uncertain loss classification. In contrast, TOGA2 correctly identifies the acceptor site, annotating the exons as T|GT…GAG|TC, which forms a TTC (Phe) sense codon and results in a fully intact transcript. (C) *C3orf20* (4930590J08Rik) intron 5 is an AT-AG U12 intron. TOGA2 correctly annotates the splice sites as TA|AT…CAG|C, forming a TAC (Tyr) sense codon across the exon junction. In contrast, TOGA1 annotates TA|AT … CTT|G, creating a TAG stop codon and leading to misclassification of the transcript as uncertain loss.

**Supplementary Figure 10:**
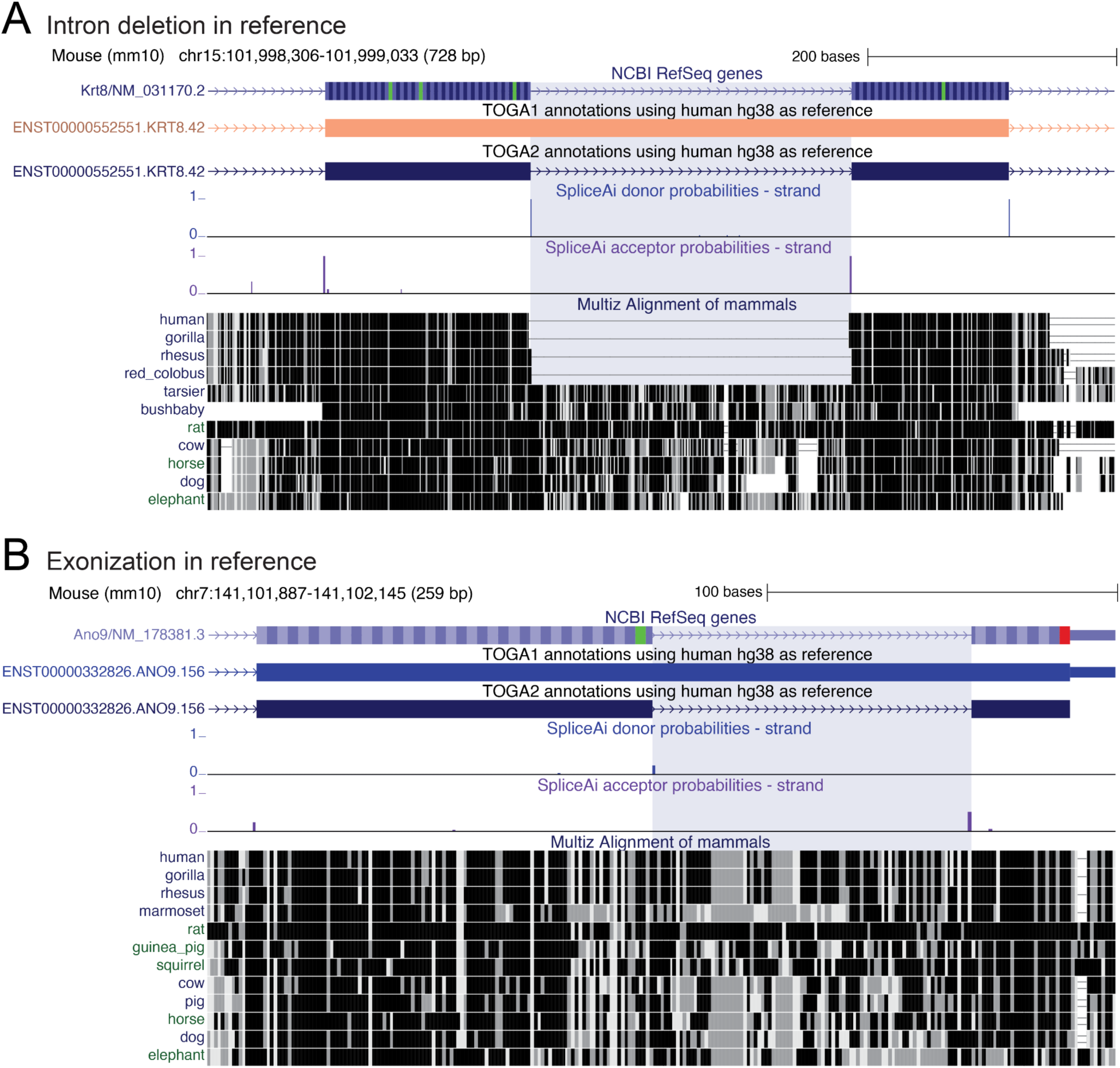
TOGA2 uses deep learning–based splice site predictions to capture gene structure changes. (A) Precise deletion of intron 5 of *KRT8* in the simian ancestor. TOGA1 with human as the reference is unaware of the existence of this intron in mouse (blue highlight) and introduces a large frameshifting insertion, leading to an uncertain loss classification. Using SpliceAI predictions, TOGA2 correctly annotates the intron, resulting in an intact transcript. (B) Exonization of the penultimate intron in *ANO9*. The intronic sequence in mouse (blue highlight) is translatable and annotated as part of an exon in several primates (orangutan, rhesus macaque, marmoset, mouse lemur), indicating an exonization event in the primate lineage. Using human as the reference, TOGA1 is unaware of this intron and treats the query sequence as coding. The transcript is then classified as an intact (and not fully intact), because the intron contains a 2 bp frameshifting deletion. Supported by SpliceAI predictions, TOGA2 correctly annotates the intron, resulting in a fully intact transcript.

**Supplementary Figure 11:**
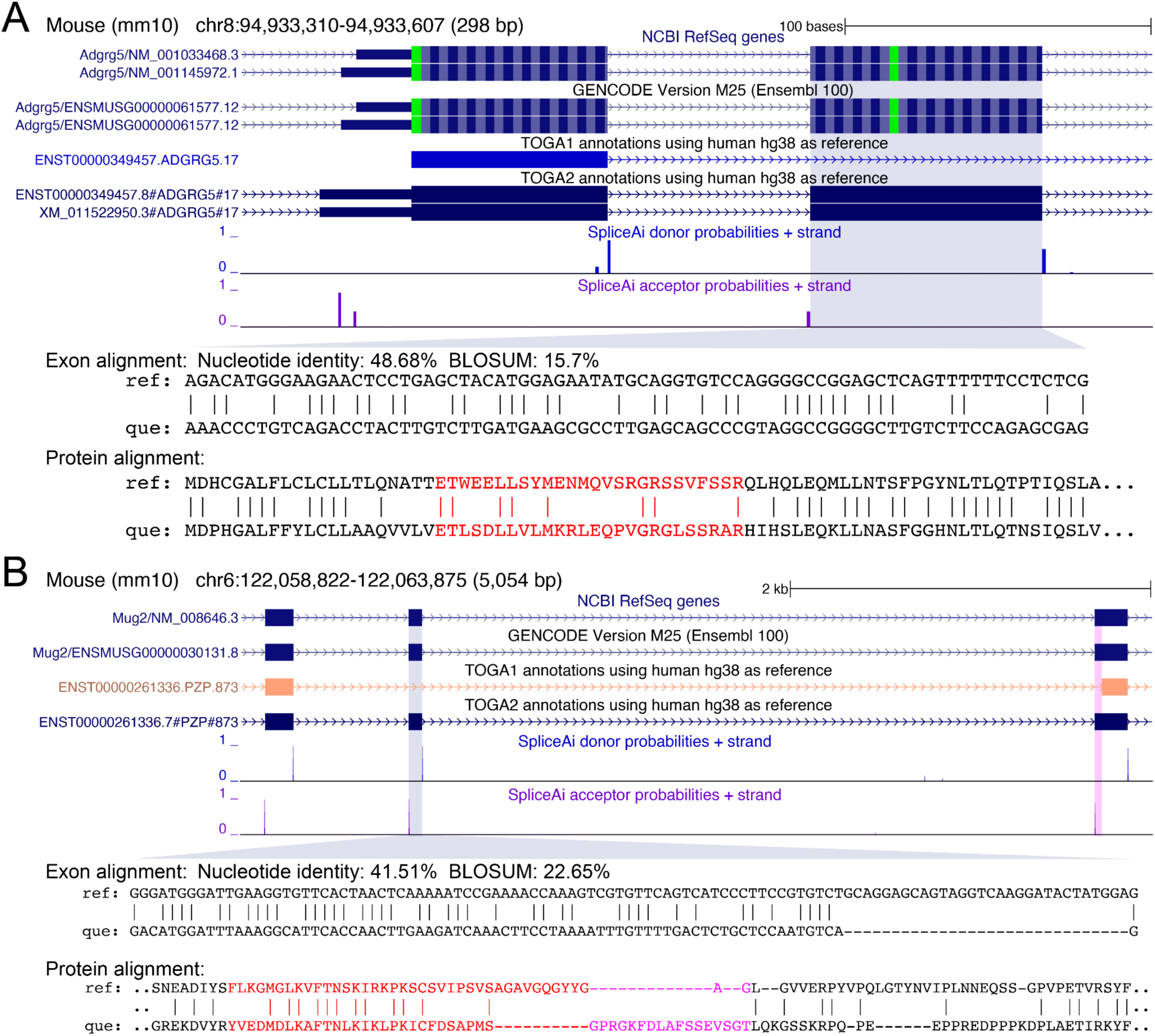
TOGA2 leverages splice site information to annotate highly diverged exons. The Viterbi algorithm used in CESAR2’s HMM may produce alignments between non-homologous sequences, for example when a query exon is deleted, overlaps an assembly gap, or is highly diverged. Therefore, both TOGA1 and TOGA2 apply two alignment metrics – nucleotide alignment identity and protein similarity (BLOSUM score) – to discard alignments for which one or both metrics are below defined thresholds (nucleotide identity ≥45%, BLOSUM ≥20%). These thresholds separate true from randomized exon alignments with a sensitivity of 0.98 and a precision of 0.99 ^38^. While TOGA2 uses the same alignment thresholds as TOGA1, it additionally leverages SpliceAI probabilities to annotate genuine, highly diverged exons when they are flanked by high-confidence splice sites, as illustrated here. (A) Coding exon 2 of *ADGRG5* exhibits numerous base substitutions and encodes a highly diverged protein sequence with minimal similarity (red font in the protein alignment refers to exon 2). TOGA1 fails to annotate this exon because its protein similarity is below the 20% threshold. In contrast, TOGA2 leverages the presence of high-confidence splice sites (SpliceAI probabilities) to correctly annotate this exon despite its low alignment identity. (B) Coding exon 17 of *PZP* (mouse ortholog is called *Mug2*) contains a large deletion (red in the protein alignment), lowering the nucleotide identity below threshold and preventing its annotation by TOGA1. Furthermore, the downstream exon 18 exhibits a splice site shift that elongates the exon in mouse (pink font in the protein alignment), which TOGA1 fails to capture, leading to classification of the transcript as uncertain loss. By leveraging SpliceAI information, TOGA2 correctly annotates exon 17 and captures the exon 18 splice site shift, resulting in a fully intact transcript.

**Supplementary Figure 12:**
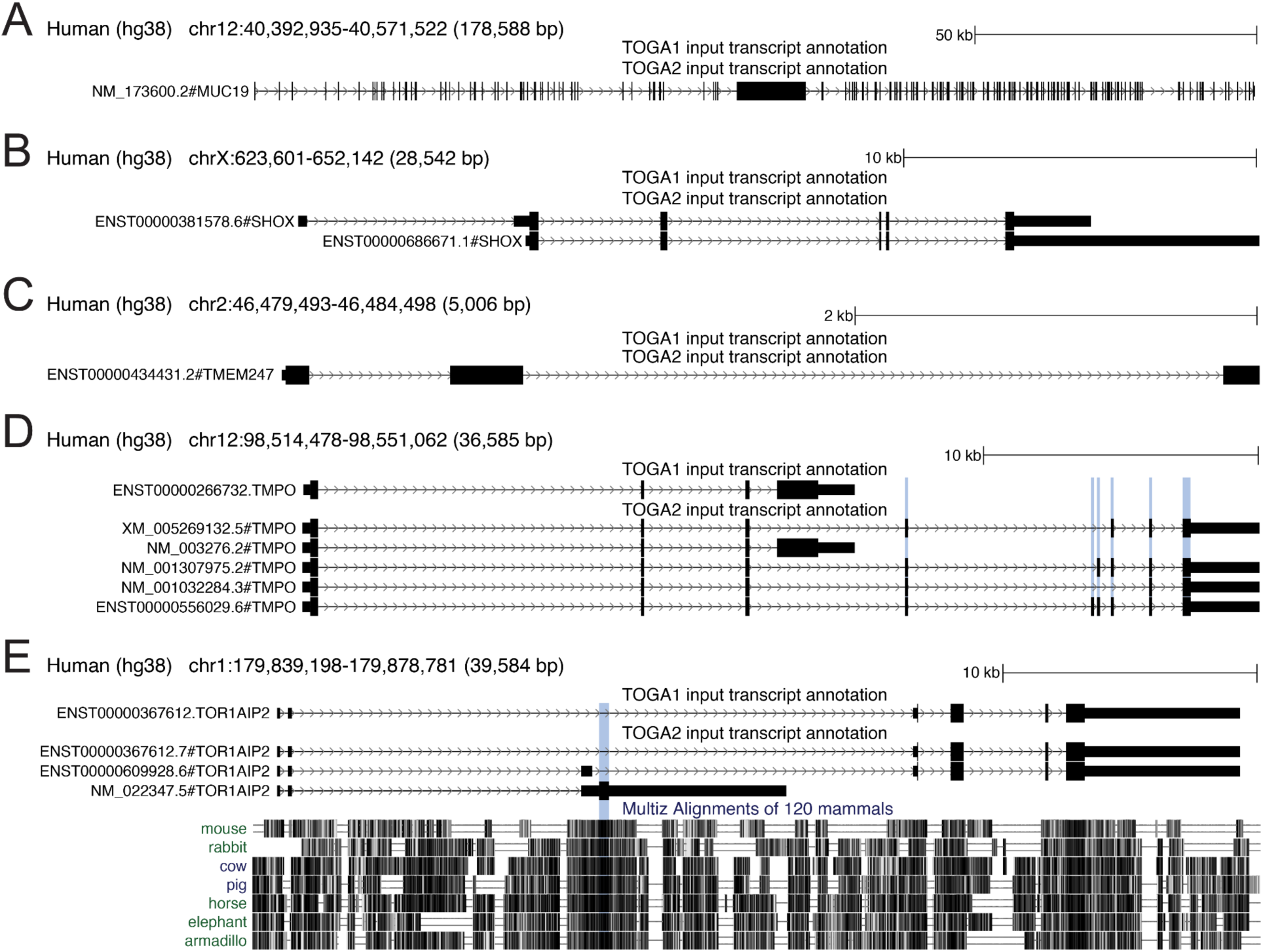
Novel genes and exons included in the TOGA2 input annotations. This figure shows examples of genes and exons missing from the TOGA1 human input annotation but included in the TOGA2 input. (A–C) Entire genes, including *MUC19* (A), *SHOX* (B), and *TMEM247* (C), were missing from the TOGA1 input, but are present in TOGA2. (D) For *TMPO*, TOGA1 includes only a transcript with four coding exons, whereas TOGA2 includes new transcripts that cover up to six additional conserved coding exons (blue highlight). (E) TOGA2 includes a new transcript of *TOR1AIP2* containing an additional coding region (blue highlight) that is highly conserved across placental mammals, as illustrated by representative species from a multiple genome alignment ^102^.

**Supplementary Figure 13:**
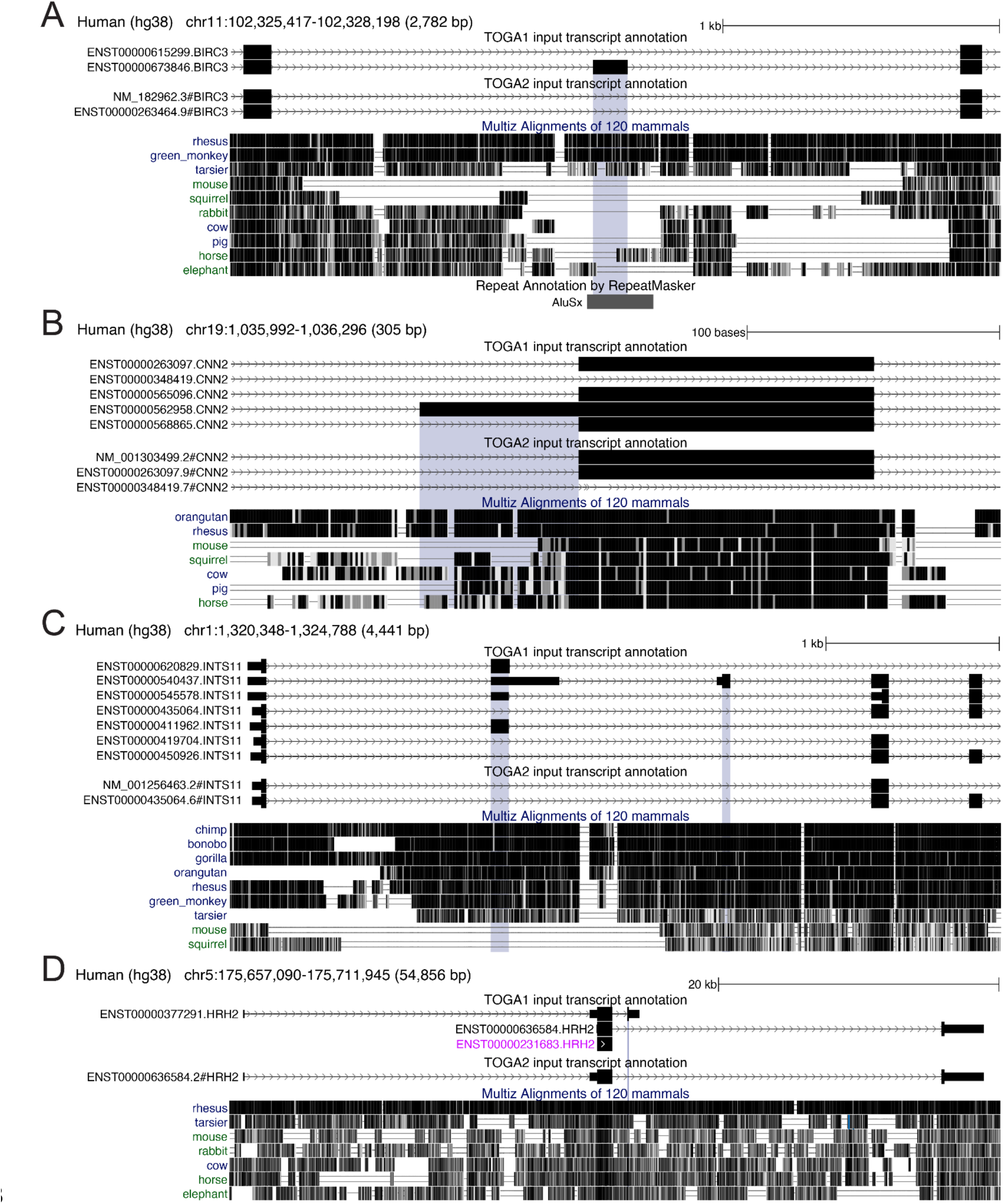
Removal of non-ancestral exons from TOGA2 input annotations. Examples of non-ancestral exons (highlight) that are specific to the reference lineage and are included in the TOGA1 but not the TOGA2 human input annotation. Lack of conservation across placental mammals is illustrated by representative species in a multiple genome alignment ^102^. Notably, lineage-specific exons increase the transcript length and are therefore preferentially kept by approaches that select the longest transcript among all available transcripts. (A) TOGA1 includes a *BIRC3* transcript containing an alternative exon derived from an Alu short interspersed nuclear element that inserted in the primate lineage. Because this exon exhibits a - 1 bp deletion in all non-human primates that have the Alu element, it is likely human-specific. TOGA2 removes this transcript. (B) TOGA1 includes a *CNN2* transcript with an alternative acceptor site that is conserved in chimpanzee, bonobo, and gorilla, but not in orangutan or many other primates. TOGA2 excludes this lineage-specific splice variant. (C) Several *INTS11* transcripts with lineage-specific exons are present in the TOGA1 input. These include an alternative second coding exon (left) that appears to be human-specific, as other primates show frameshifts or lack splice sites, and an alternative first coding exon (right) that is only partially conserved among simian primates, as it lacks a start codon in several species (e.g. gibbon, Angolan colobus, and tarsier). TOGA2 removes these transcripts. (D) TOGA1 includes an *HRH2* transcript with an alternative terminal coding exon that is intact only in human, chimpanzee, and gorilla. TOGA2 removes this transcript, along with a truncated single-exon transcript (purple font).

**Supplementary Figure 14:**
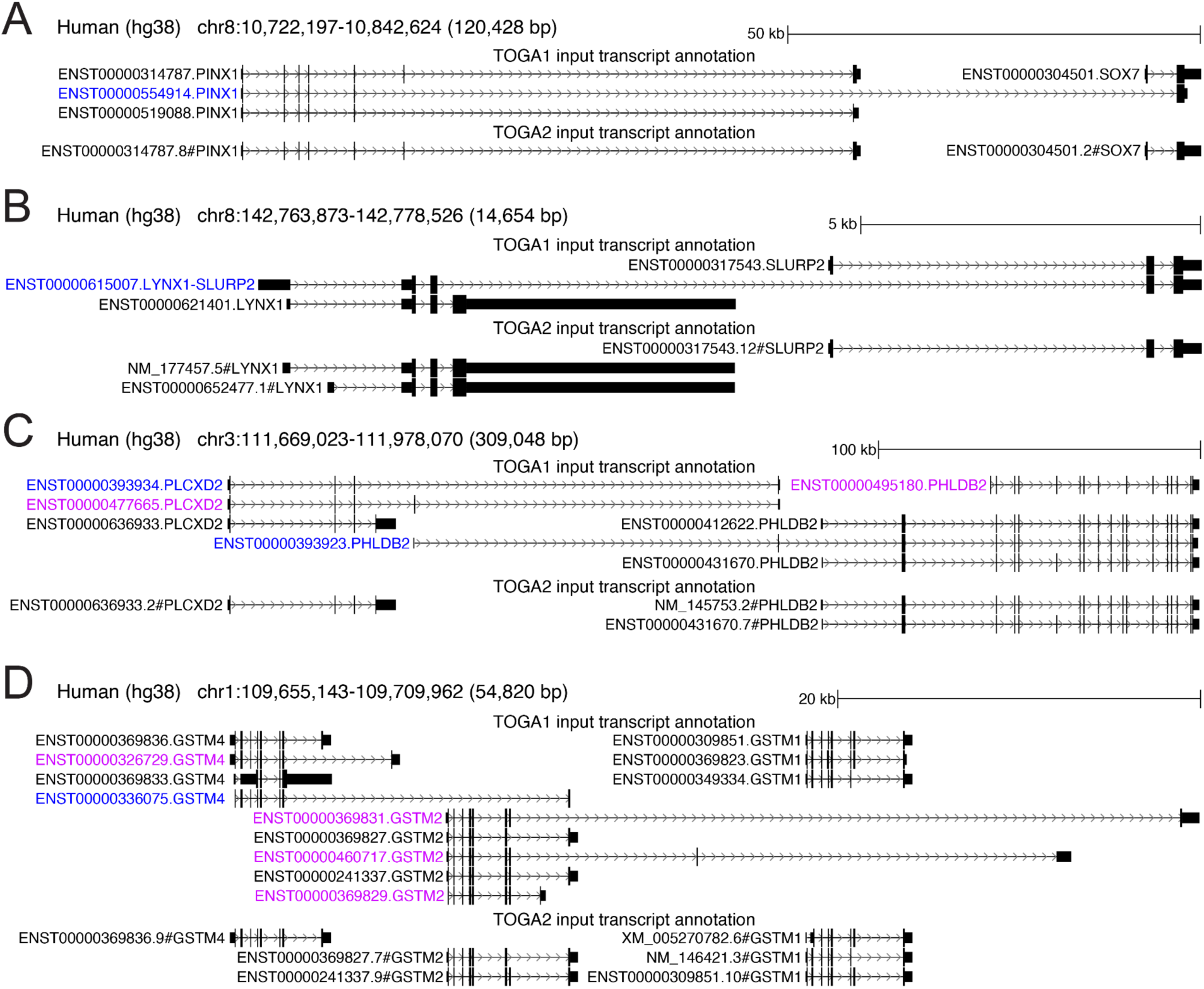
Removal of gene fusion transcripts in TOGA2 input annotations. Fusion transcripts in the reference species that span the CDS of two distinct neighboring genes lead to incorrect orthology assignments in TOGA2. We therefore rigorously removed these transcripts from the input annotations. Fusion transcripts highlighted in blue font in this figure. (A) A fusion transcript connecting *PINX1* and *SOX7* is present in the TOGA1 human input annotation, resulting in a many-to-one orthology classification for both genes. The TOGA2 input annotation removes this transcript, restoring the correct 1:1 orthology classification. (B) A fusion transcript spanning *LYNX1* and *SLURP2* in the TOGA1 input leads to a many-to-one classification. Removing this transcript in TOGA2 results in the correct 1:1 orthology. (C) Several fusion transcripts connecting *PLCXD2* and *PHLDB2* in the TOGA1 input result in incorrect many-to-one classifications. TOGA2 removes these transcripts, restoring one-to-one orthology. Other transcripts containing non-conserved coding exons (purple font) are also removed. (D) A fusion transcript spanning *GSTM4* and *GSTM2* is removed in the TOGA2 input annotations, along with additional transcripts containing non-conserved exons (purple font).

**Supplementary Figure 15:**
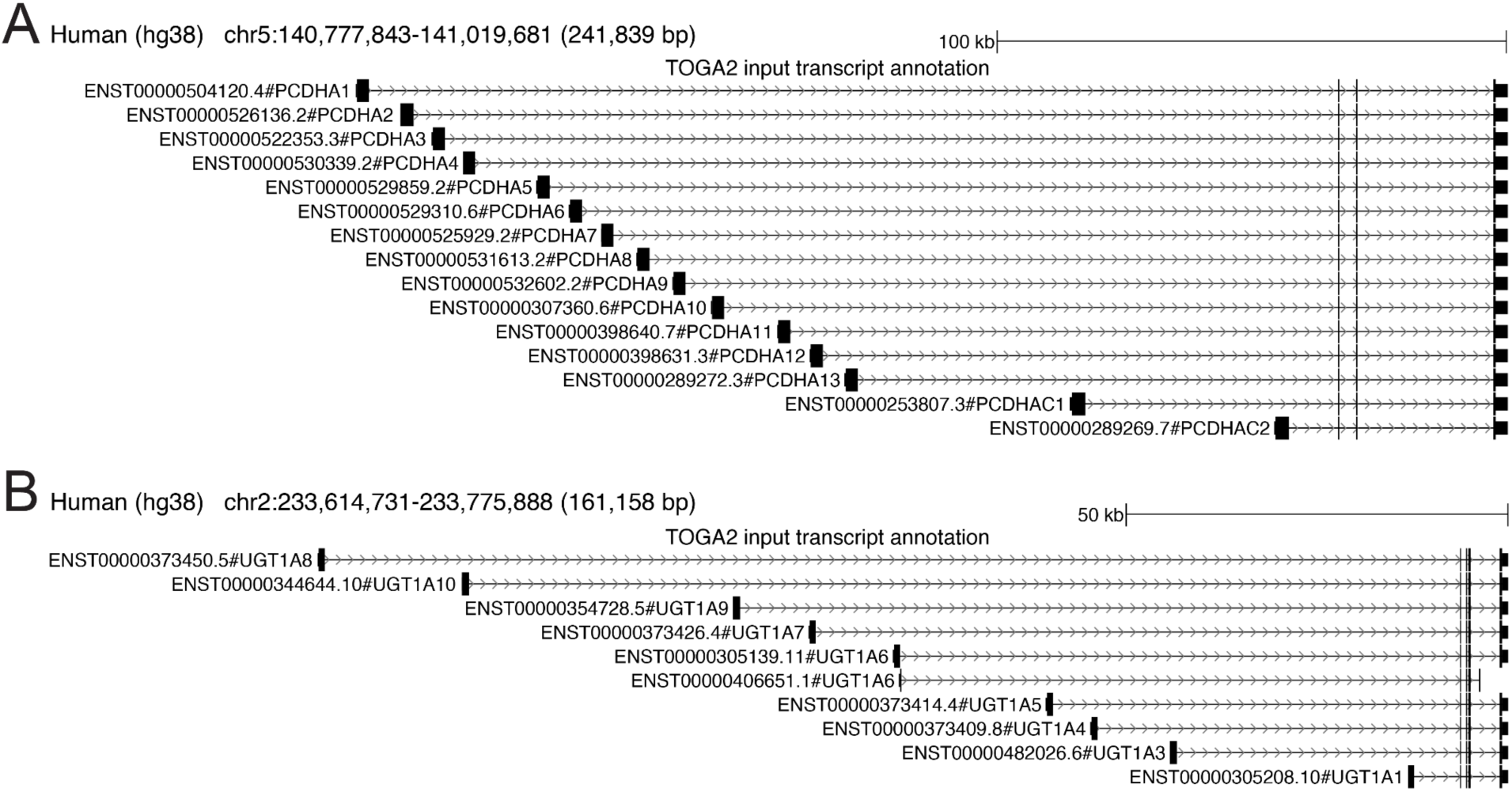
TOGA2 preserves transcripts that resemble fusion events but originate from a single gene. To remove candidate fusion transcripts from the input annotations, TOGA2 identifies transcripts with same-strand coding exon overlap that are assigned to different genes. However, this approach also captures cases where genes with distinct names actually represent alternative transcripts from a single gene. Through manual curation, we removed bona fide fusion transcripts, while retaining such fusion-mimicking transcripts, as illustrated here. (A) The *PCDHA* (protocadherin alpha) gene cluster comprises multiple transcripts that share downstream exons and only differ in their first coding exon. (B) *UGT1A* (UDP-glucuronosyltransferase) genes similarly differ only in their first coding exon while sharing downstream exons. Unlike TOGA1, TOGA2 incorporates prior knowledge of such overlapping reference genes into the gene inference procedure and infers separate query genes. As a result, TOGA2 correctly assigns 1:1 or, in the presence of inactivating mutations, 1:0 orthology relationships to the individual *PCDHA* and *UGT1A* genes, whereas TOGA1 incorrectly assigns many:1 or many:many orthology relationships.

**Supplementary Figure 16:**
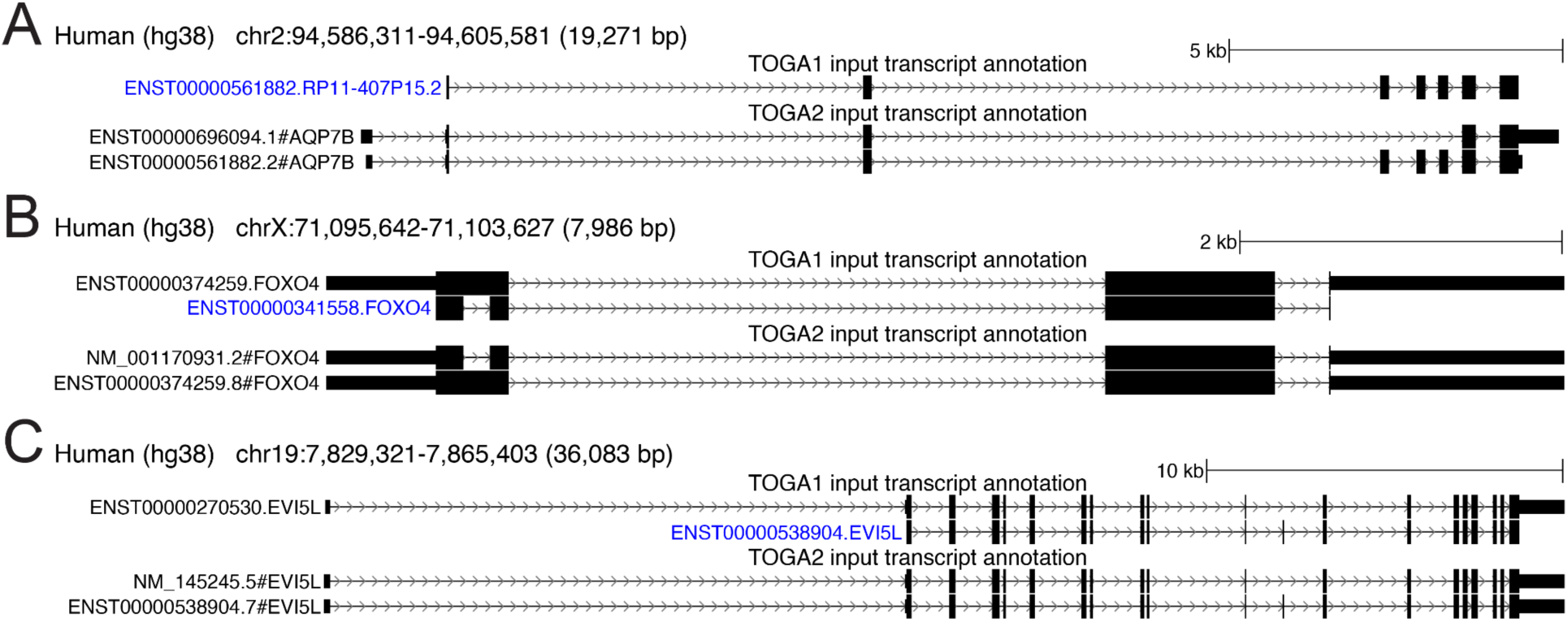
Inclusion of UTR-containing transcripts in TOGA2 input annotations. Reference transcripts lacking annotated UTRs can interfere with TOGA2’s orthology inference, as alignments overlapping UTR regions may be misinterpreted as flanking intergenic sequence, which affects the alignment-based features used for orthology classification. To address this, TOGA2 input annotations preferentially include transcripts with annotated UTRs. Furthermore, TOGA2 can annotate UTRs in the query species. The figure shows three genes for which the human TOGA1 input annotation contains transcripts lacking UTRs (blue font), whereas the TOGA2 input annotation includes transcripts with annotated UTRs. (A) *AQP7B*. This example also illustrates that the gene symbol (previously RP11-407P15.2) is updated. (B) *FOXO4*. (C) *EVI5L*.

**Supplementary Figure 17:**
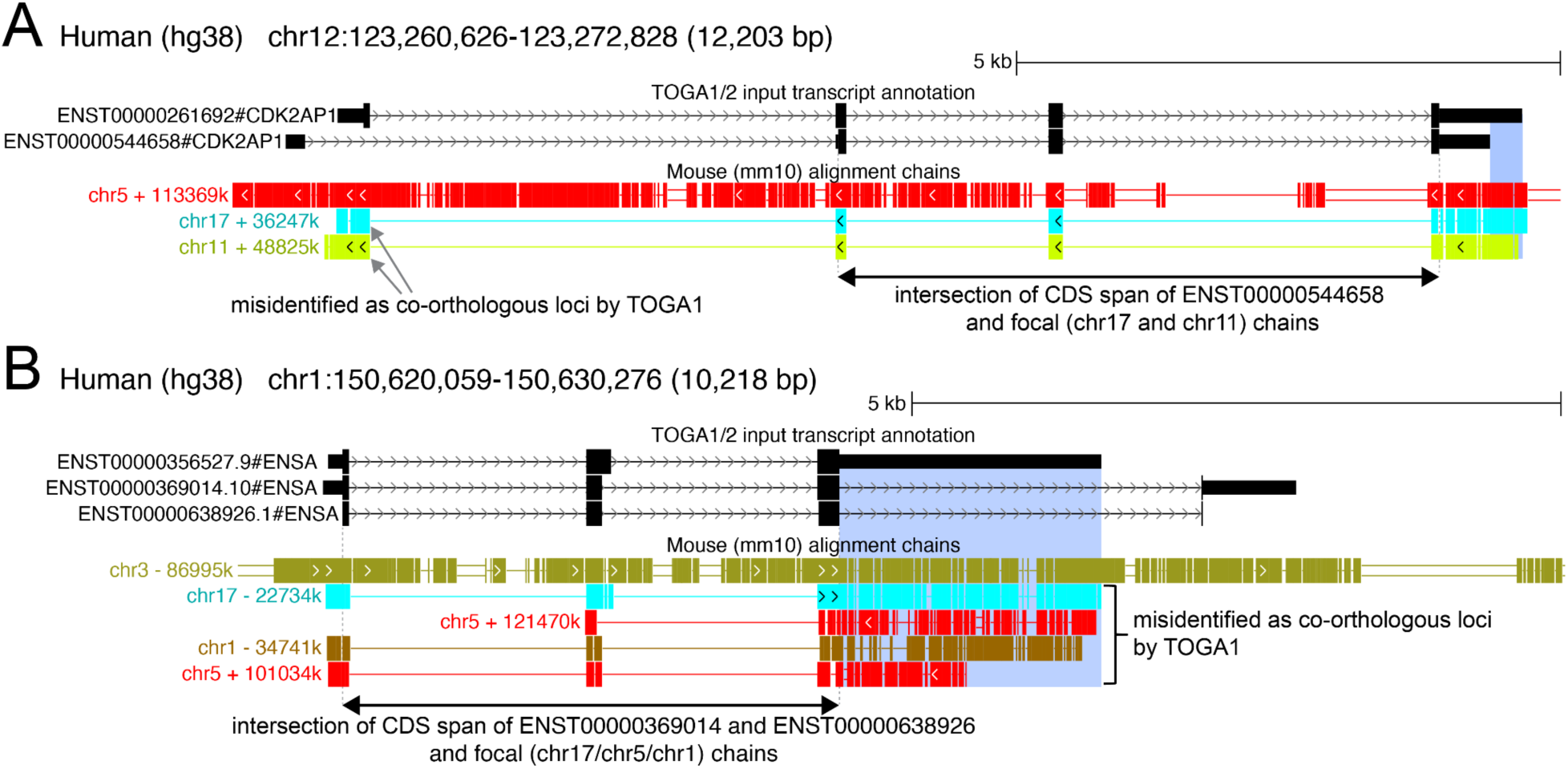
Improved processed pseudogene identification in TOGA2 prevents false orthology assignments. (A) The top chr5 chain represents the only orthologous *CDK2AP1* locus in mouse and is correctly identified as such by both TOGA1 and TOGA2. TOGA1 additionally identifies the two other chains as orthologous mouse loci with orthology probabilities of 0.72 and 0.8, respectively. These chains represent processed pseudogene copies derived from the transcript with the longer 3′ UTR (blue highlight). When the transcript with the shorter 3′ UTR (ENST00000544658) is considered, these aligned regions are incorrectly interpreted by TOGA1 as flanking intergenic alignments, resulting in the incorrect identification of two additional orthologs and a false 1:many orthology relationship. In contrast, TOGA2 uses clipped intron coverage as a new feature, which considers only CDS exons of the focal transcript and the introns between them (indicated by the double-headed arrow). This enables TOGA2 to correctly classify these chains as processed pseudogene copies. Consequently, TOGA2 classifies *CDK2AP1* as a 1:1 ortholog in mouse, annotating the chr11 locus as a retrogene candidate and the chr17 locus as a processed pseudogene. (B) The top chr3 chain represents the only orthologous *ENSA* mouse locus, correctly identified by both TOGA1 and TOGA2. TOGA1 additionally identifies the four other chains as orthologous *ENSA* loci, assigning them high orthology probabilities of 0.93-0.95. However, these chains represent processed pseudogene copies derived from the three-exon *ENSA* transcript containing a long 3′ UTR (blue highlight). When the four-exon transcripts (ENST00000369014 and ENST00000638926) are considered, this aligned region is incorrectly interpreted as an intronic alignment, explaining the high orthology probabilities. While the chr5 and chr1 loci contain processed pseudogene copies lacking an intact reading frame, the gene annotated by TOGA1 in the chr17 locus retains an intact reading frame and is therefore incorrectly classified as a second ortholog, resulting in a false 1:many orthology relationship. In contrast, TOGA2 correctly classifies the four other chains as processed pseudogene copies, resulting in classifying *ENSA* as a 1:1 ortholog in mouse, and annotating a retrogene candidate in the chr17 locus and processed pseudogenes in the remaining three loci.

**Supplementary Figure 18:**
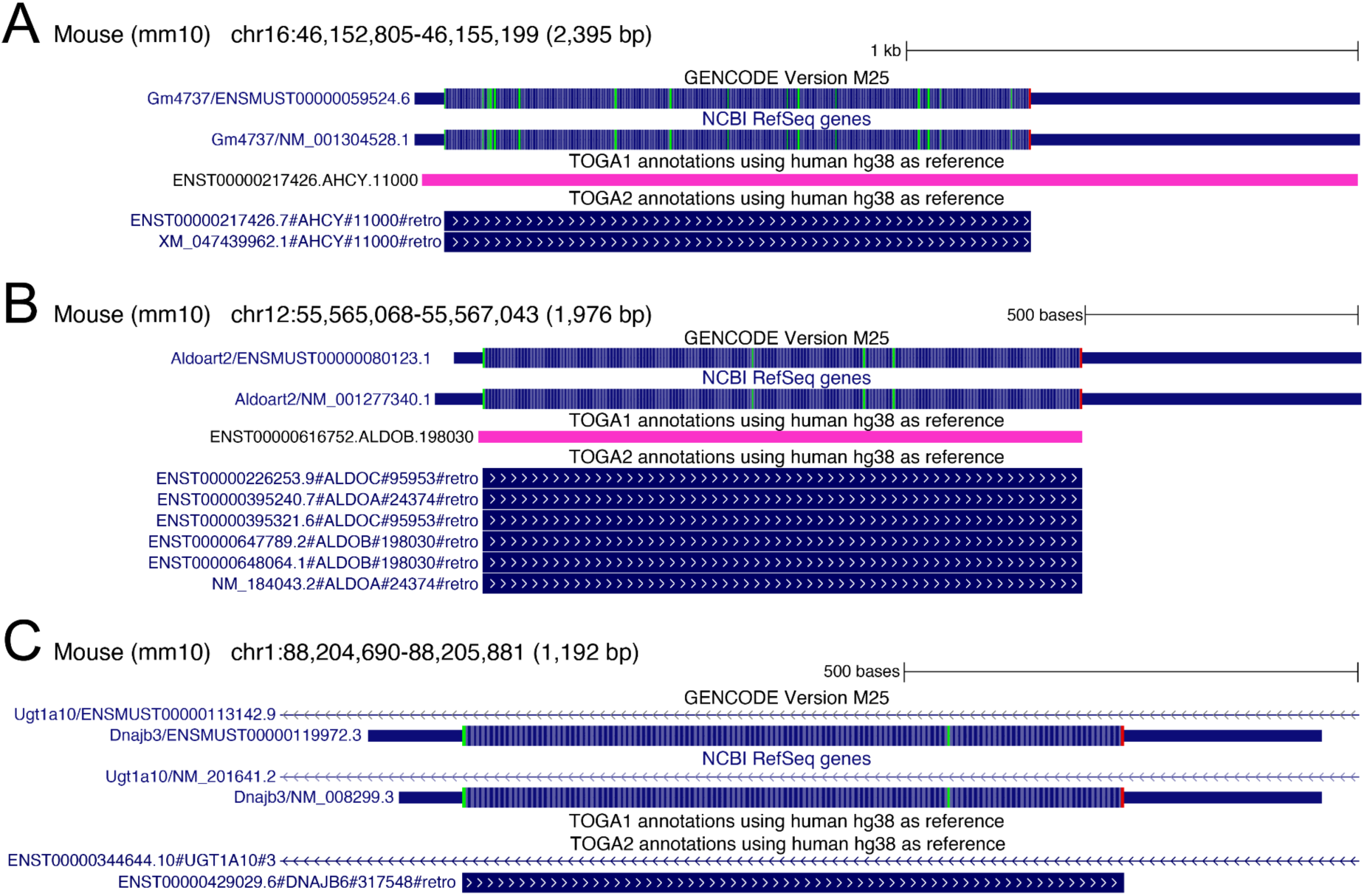
TOGA2 improves processed pseudogene identification and annotates retrogene candidates. (A-B) In both examples, TOGA1 and TOGA2 identify a processed pseudogene in the respective mouse locus. Whereas TOGA1 does not distinguish processed pseudogene copies with intact reading frames (retrogene candidates), TOGA2 recognizes these loci as retrogene candidates and annotates the putative coding sequences. The resulting TOGA2 annotations precisely match genes annotated by GENCODE and RefSeq. Notably, in panel A, GENCODE and RefSeq annotate the gene as *Gm4737* (an automatically derived gene identifier), whereas TOGA2 identifies it as a retrocopy of *AHCY*. (C) Only TOGA2, but not TOGA1, identifies a processed pseudogene in this locus, illustrating the improved processed pseudogene detection in TOGA2. The annotated retrogene candidate precisely matches a gene annotated by GENCODE and RefSeq.

**Supplementary Figure 19:**
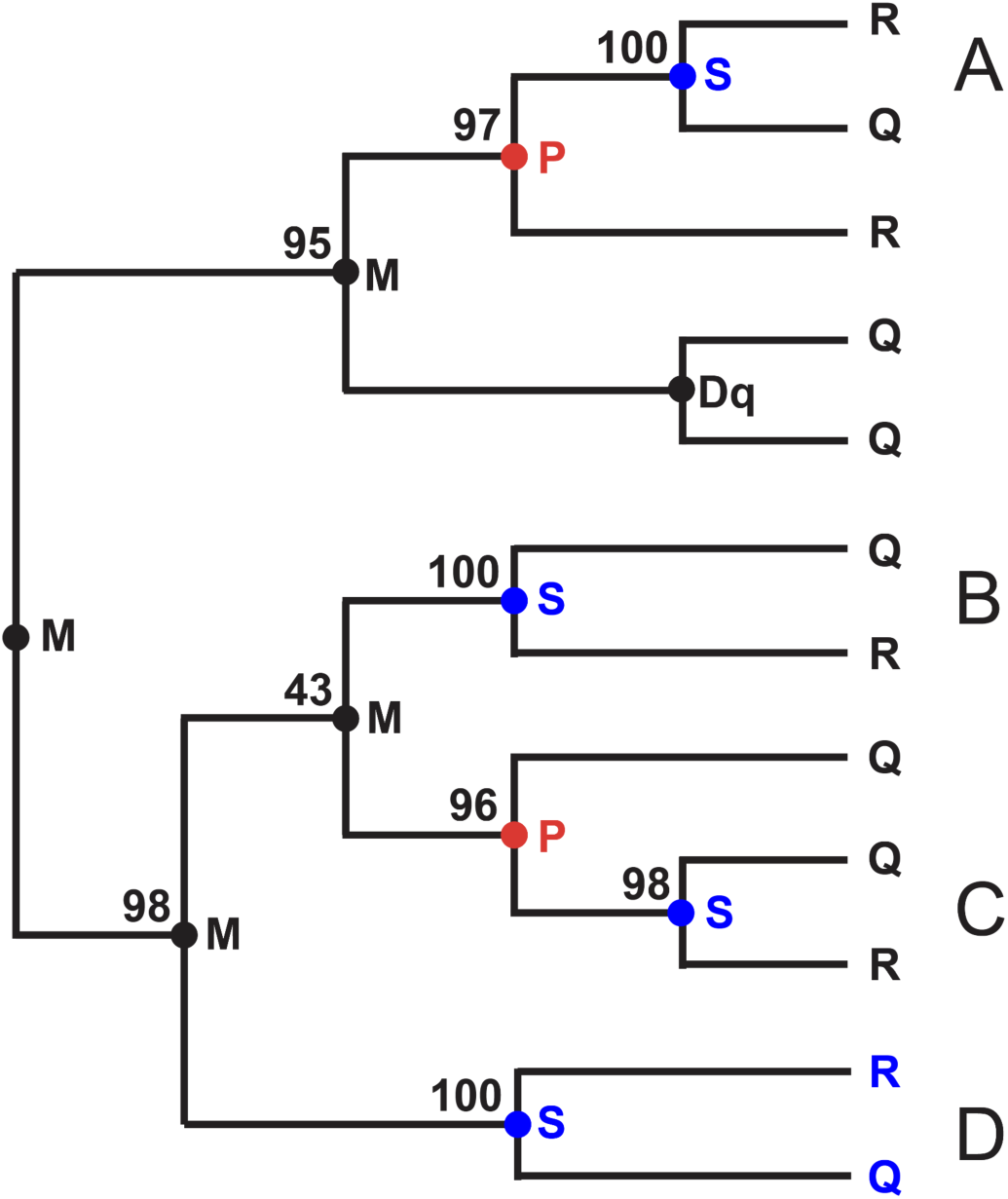
Illustration of the gene tree node-labeling algorithm used to identify well-supported 1:1 orthologs. The figure shows a gene tree containing reference (R) and query (Q) genes together with internal node labels and bootstrap support values. The tree contains four candidate 1:1 ortholog pairs, labeled A-D. (A) TOGA2 does not resolve this pair as a 1:1 ortholog, because a problematic node is present on the path to the root. Resolving this pair would result in a sister subtree that contains only a reference gene without any remaining orthology relationship. (B) TOGA2 does not resolve this pair because the path to the root contains a node with low bootstrap support (43), indicating insufficient confidence in the orthology relationship. (C) TOGA2 does not resolve this pair because the path to the root contains both a problematic node and a node with low bootstrap support (43). (D) This pair is identified as an additional 1:1 ortholog (blue font), because the path to the root contains no problematic nodes, all visited nodes exhibit high bootstrap support, and orthology relationships for the remaining genes in the tree are preserved.

**Supplementary Figure 20:**
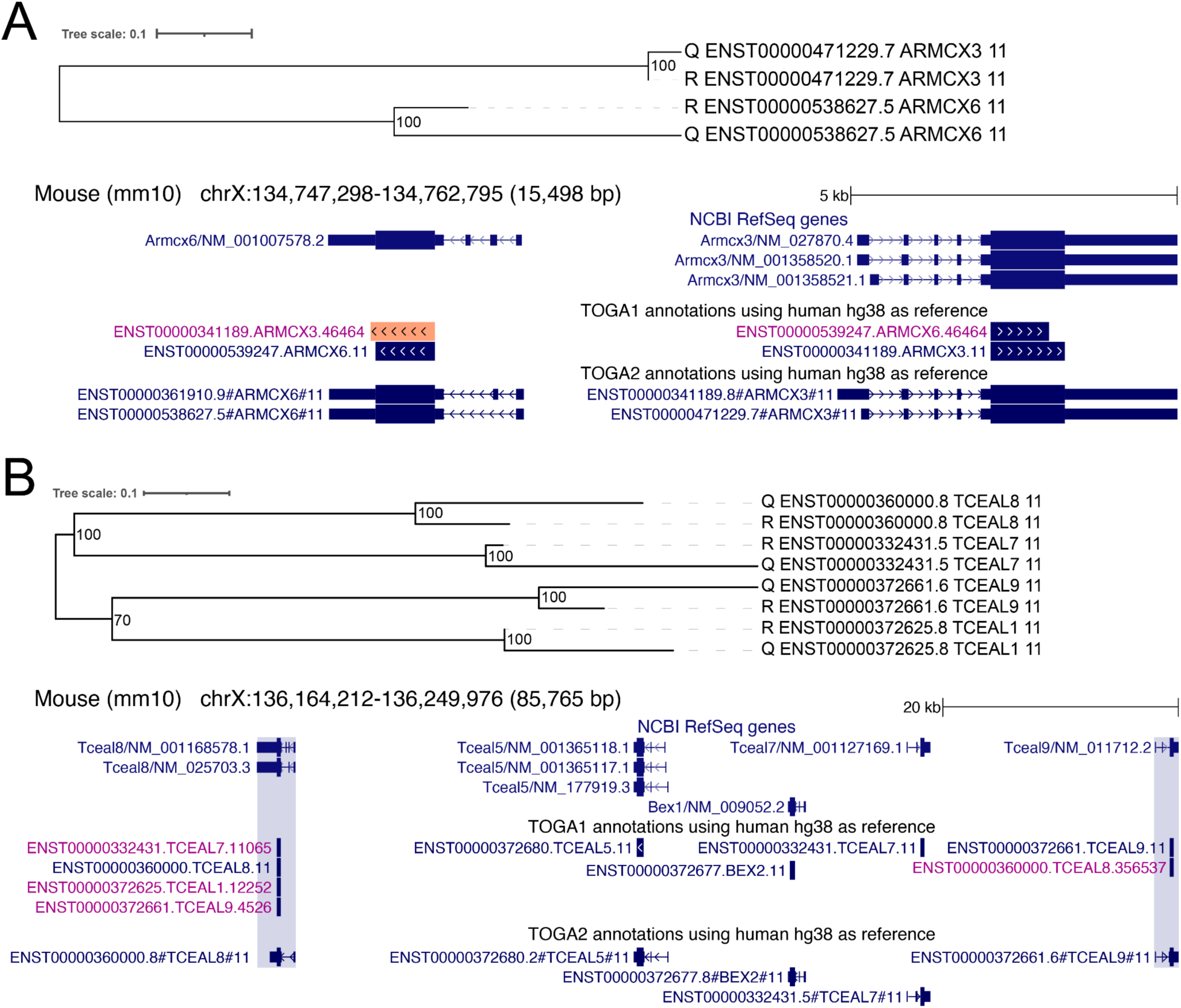
Examples of newly identified 1:1 orthologs using gene tree reconciliation. (A) Top: *ARMCX3* and *ARMCX6* initially form a many:many (2:2) orthology group in mouse. However, the gene tree supports both genes as reciprocal 1:1 orthologs with 100% bootstrap support, enabling TOGA2 to separate them into two 1:1 orthology relationships. Bottom: Genome browser screenshot showing the mouse locus containing *ARMCX3* and *ARMCX6*. Because TOGA1 does not perform gene tree reconciliation, it incorrectly annotates *ARMCX6* in the *ARMCX3* locus and vice versa (gene labels shown in red). In contrast, TOGA2 assigns the correct orthology relationships, matching the NCBI RefSeq annotation. (B) Top: *TCEAL1*, *TCEAL7*, *TCEAL8*, and *TCEAL9* initially form a many:many orthology group. Gene tree reconciliation identifies *TCEAL7* and *TCEAL8* as 1:1 orthologs with strong bootstrap support. In contrast, the split of *TCEAL1* and *TCEAL9* is supported by only 70% bootstrap support, which does not satisfy the 90% support threshold required by TOGA2. After identifying the new 1:1 orthology relationships, TOGA2 removes conflicting orthology relationships involving *TCEAL7* and *TCEAL8* (*TCEAL1-TCEAL8*, *TCEAL8-TCEAL7*, and *TCEAL8-TCEAL9*). As a result, *TCEAL1* and *TCEAL9* are no longer linked to other orthologs and consequently also become 1:1 orthologs. Bottom: Genome browser screenshot showing that TOGA1 assigns incorrect orthology relationships (gene labels shown in purple), whereas TOGA2 resolves them correctly. *TCEAL1* is located further away from this locus and is omitted for visual clarity.

**Supplementary Figure 21:**
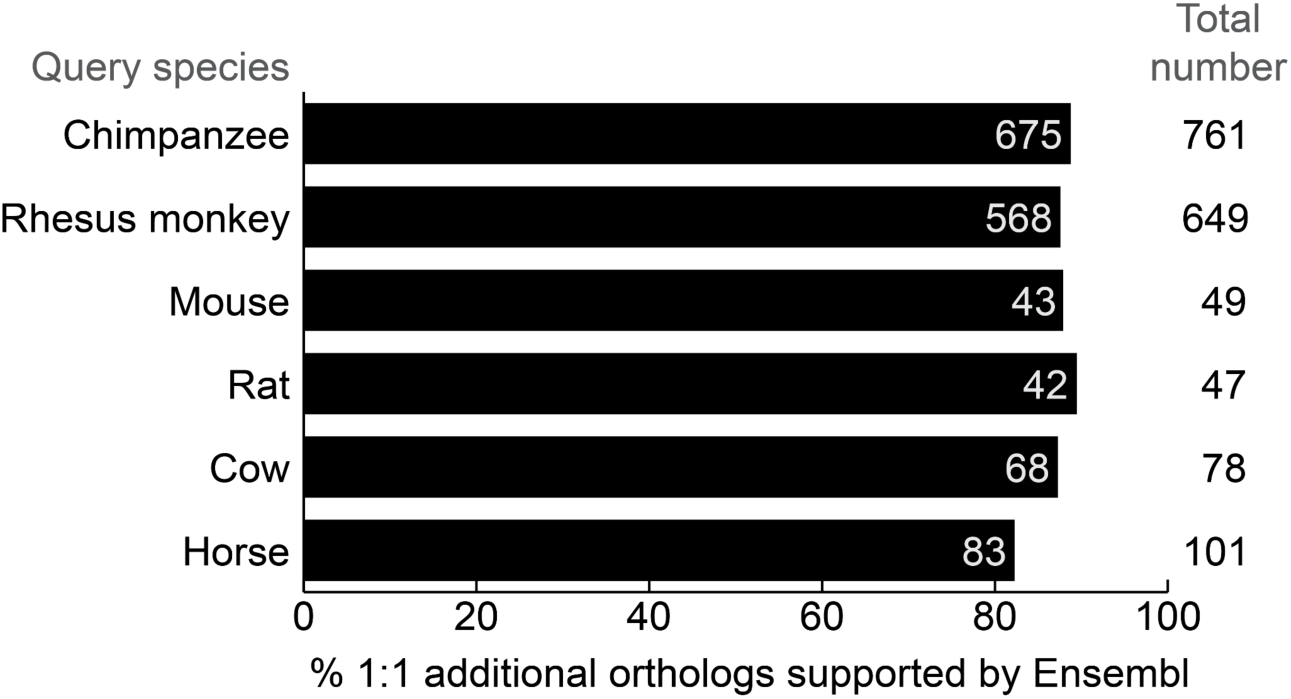
TOGA2 improves the resolution of orthology relationships. Using the new gene tree reconciliation approach, TOGA2 identifies additional 1:1 orthologs compared to TOGA1. With human as reference, the bar charts show that a consistently high percentage of these newly identified 1:1 orthologs are also classified as 1:1 orthologs by Ensembl Genes 115 (Dyer et al., 2025), indicating a high accuracy. The absolute numbers of newly identified and Ensembl-supported 1:1 orthologs are also shown.

**Supplementary Figure 22:**
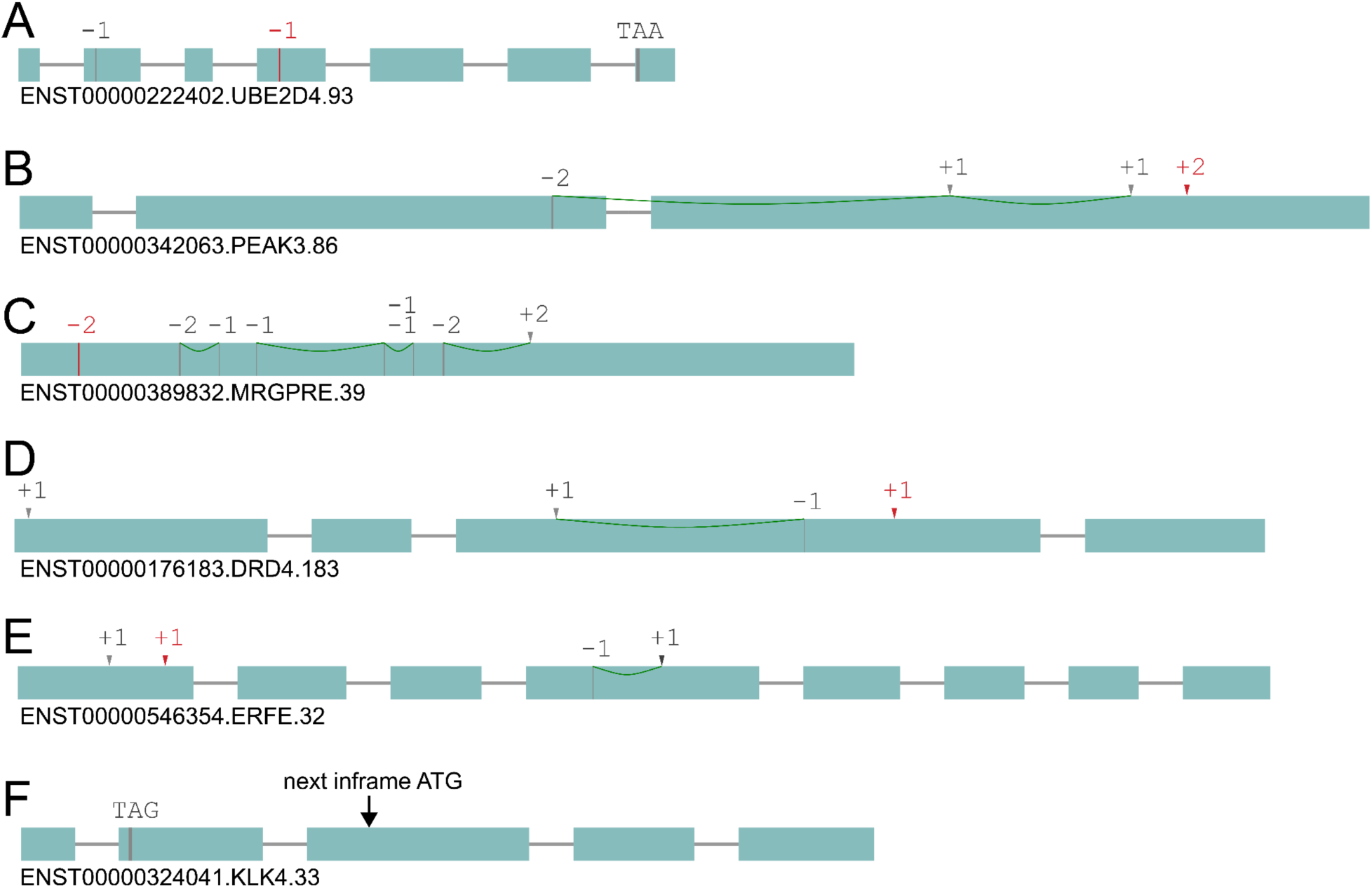
TOGA2 improves identification of lost genes. Exon-intron structure visualizations showing inactivating mutations. Premature stop codons are shown as black vertical lines, frameshifting deletions as red vertical lines, and frameshifting insertions as red arrowheads. Compensating frameshifts are connected by green lines. Mutations shown in grey are masked by TOGA1 because they occur within the first or last 10% of the reading frame or because they are compensating frameshifts. The examples shown in panels A-E were classified as uncertain loss by TOGA1. By applying the improved loss identification rules described in the Methods, TOGA2 classifies these transcripts as lost. Panel F illustrates the improved classification of N-terminal mutations, resulting in a change from intact for TOGA1 to uncertain loss for TOGA2. (A) TOGA2 unmasks the N-terminal frameshift and the C-terminal stop codon because the transcript contains an unmasked inactivating mutation (a frameshift in exon 4). (B–D) Large portions of the coding sequence are affected by compensating frameshifts. In contrast to TOGA1, TOGA2 treats the region between compensating frameshifts as deleted. (E–F) Examples containing both compensating frameshifts and terminal mutations that become unmasked because additional inactivating mutations are present in the transcript. (F) The stop-codon mutation is located within the first 10% of the reading frame; however, the next in-frame ATG where translation could potentially reinitiate occurs at 38% of the reading frame (marked by an arrow), which would truncate approximately one third of the protein. Whereas TOGA1 classified this transcript as intact, TOGA2 unmasks this mutation, resulting in an uncertain loss classification.

**Supplementary Figure 23:**
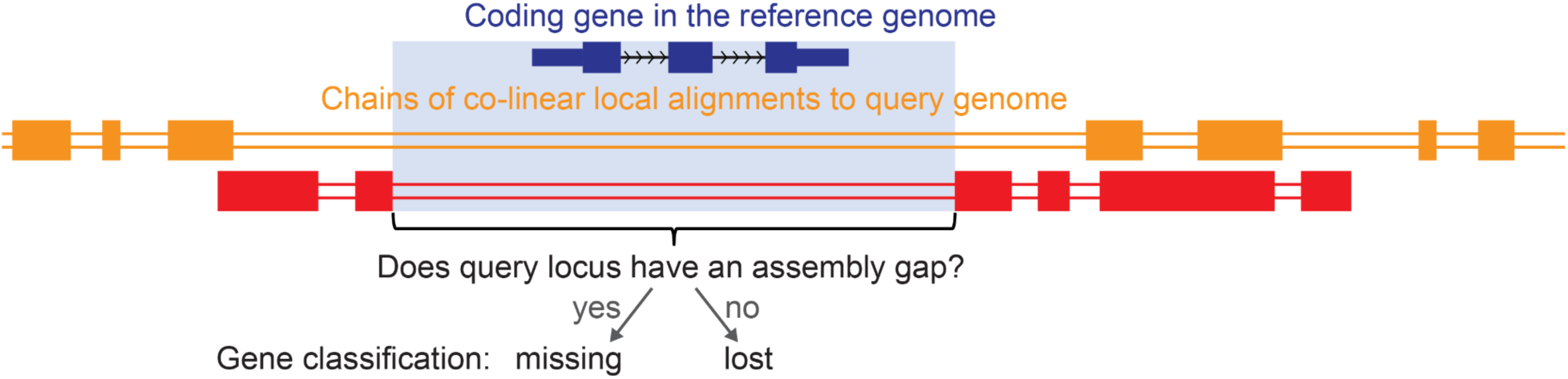
Illustration of the “most nested spanning chain” to distinguish missing from lost genes. For the focal gene, two alignment chains span the gene but contain no aligning blocks overlapping its coding exons. TOGA2 identifies the most likely orthologous chain as the most nested spanning chain by determining, for each chain, the reference interval between the closest upstream and downstream aligning blocks and selecting the chain with the smallest such interval. The query locus corresponding to this chain is then examined for assembly gaps. If the locus contains an assembly gap, the gene is classified as missing, indicating assembly incompleteness at the likely orthologous locus. Otherwise, the gene is classified as lost.

**Supplementary Figure 24:**
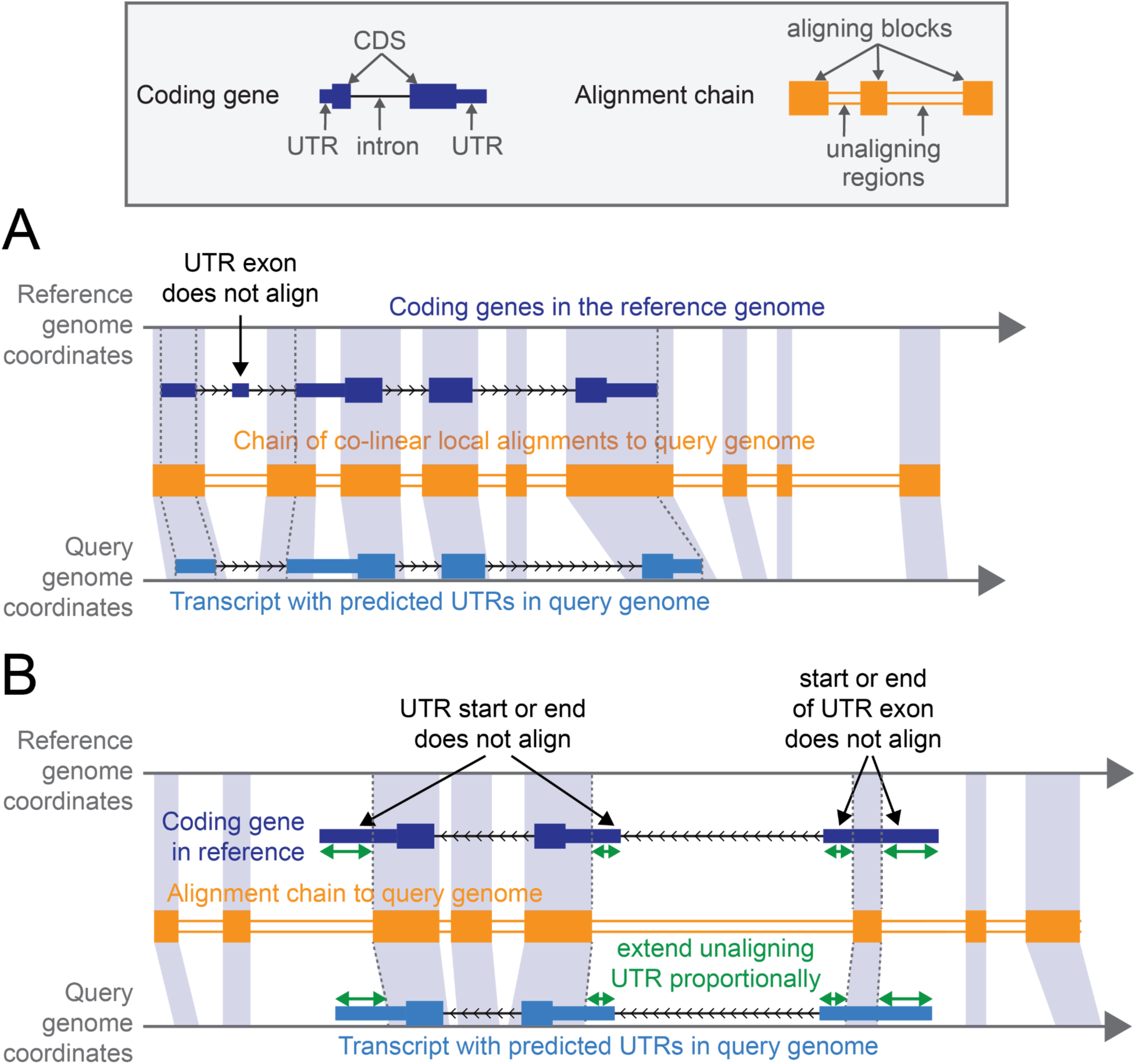
Illustration of TOGA2’s procedure to annotate UTRs. (A) Illustration of a gene for which the start and end coordinates of UTR exons and UTR regions align between reference and query. The corresponding query coordinates, inferred from aligning chain blocks, are used to add UTR information to the exon-intron structure of the query transcript. Note that the non-aligning UTR exon is ignored, because its position cannot be inferred. (B) Illustration of a transcript for which the start or end coordinates of UTR regions do not align to the query genome. In this case, TOGA2 first uses the coordinates of the first and last aligning bases within the UTR regions to infer the corresponding UTR positions in the query. Next, as indicated by the green arrows, the predicted UTRs are proportionally extended according to the length of the unaligning UTR regions in the reference, yielding the final UTR prediction in the query genome.

**Supplementary Figure 25:**
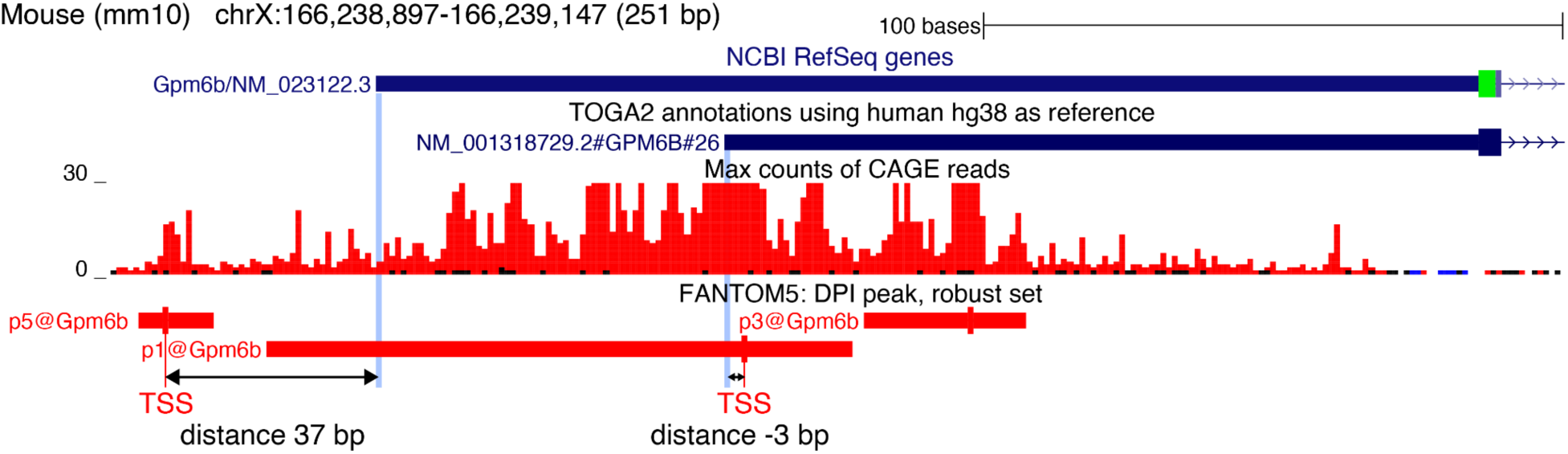
Overlap with CAGE peaks and distance to the closest CAGE-defined TSS. illustration of RefSeq- and TOGA2-annotated TSSs overlapping a CAGE peak. The distance between the RefSeq TSS and the closest CAGE-defined TSS is 37 bp, whereas the distance between the TOGA2 TSS and the closest CAGE-defined TSS is -3 bp. Note that for RefSeq the closest CAGE-defined TSS is located in a different CAGE peak than the peak overlapped by the RefSeq TSS.

**Supplementary Figure 26:**
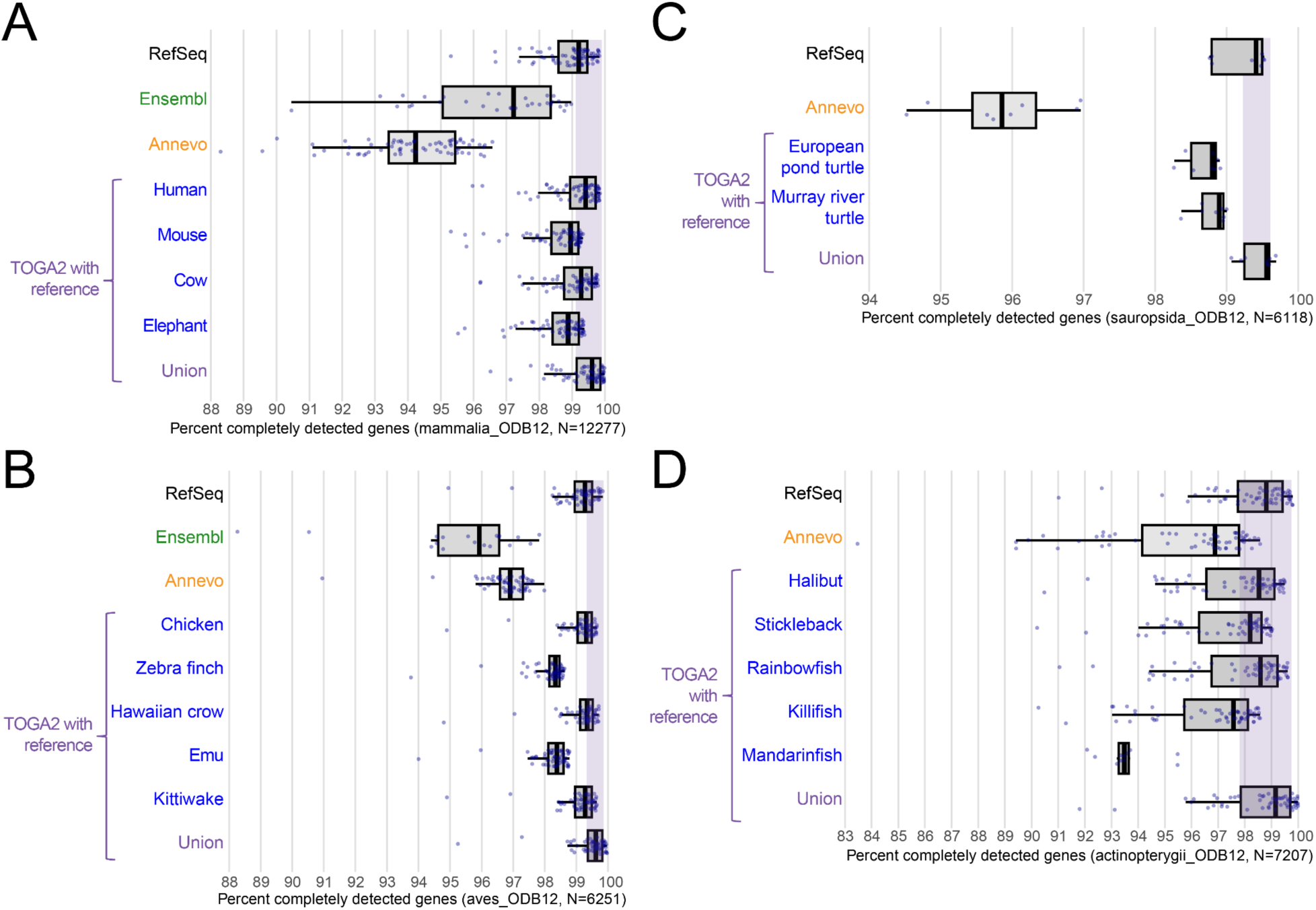
TOGA2 generally achieves higher annotation completeness. Box plots show the percentage of completely detected, near-universally conserved genes from clade-specific BUSCO ODB12 gene sets in 70 placental mammals (A), 56 birds (B), eight turtles (C), and 58 percomorph fish (D). Annotations generated by TOGA2 using single reference species and the union of all reference species are compared with NCBI RefSeq, Ensembl, and Annevo. Ensembl annotations are included only when available for the same genome assembly. Each dot represents one assembly, corresponding to those analyzed in the main text figure. Boxes show the first and third quartiles, with the median indicated by the horizontal line. Whiskers extend to 1.5 times the interquartile range.

**Supplementary Figure 27:**
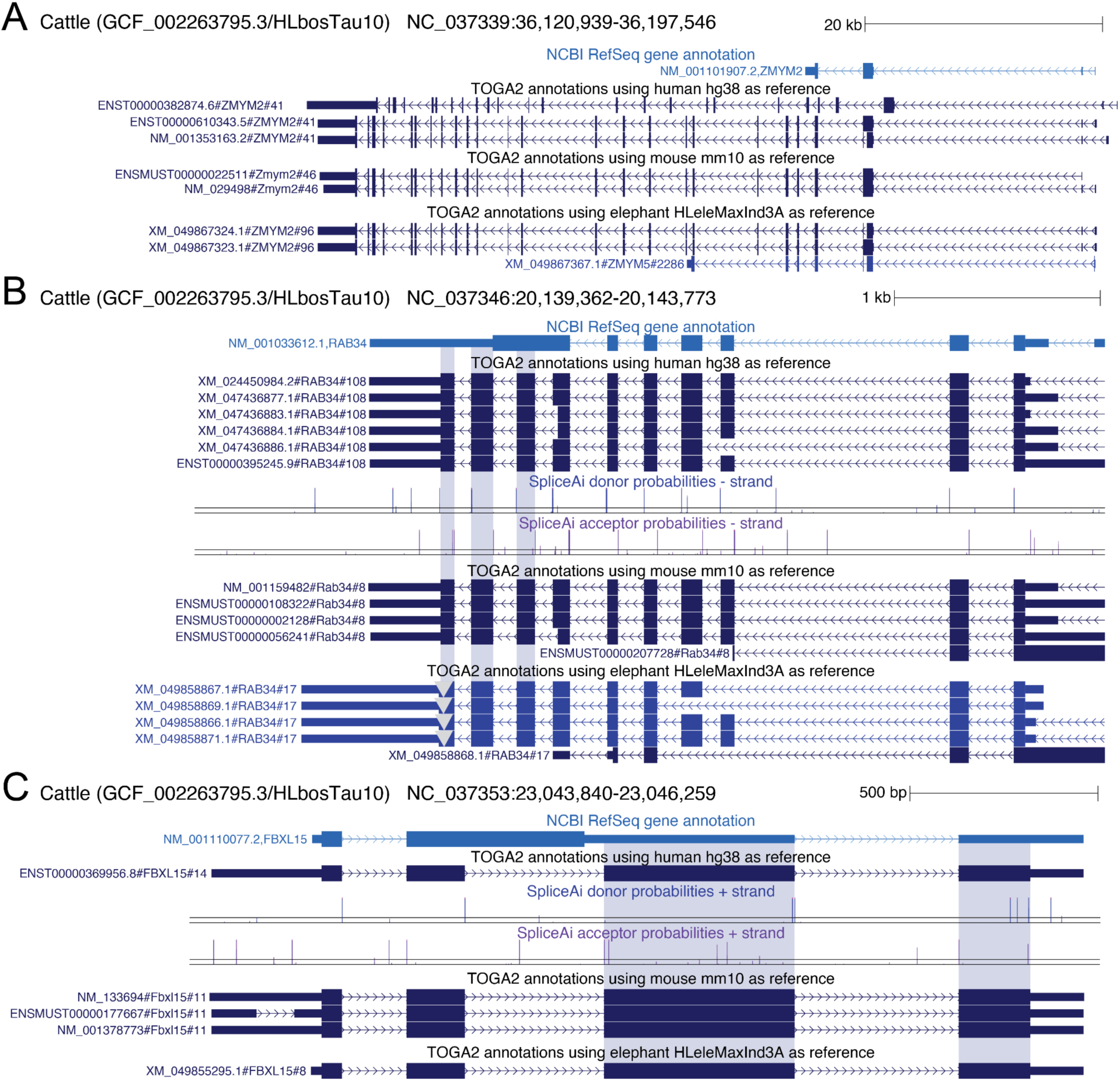
TOGA2 increases annotation completeness for cow. Examples of near-universally conserved genes present in the mammalia_odb12 gene set that are completely detected in TOGA2 annotations of the cow GCF_002263795.3 assembly, but are fragmented or missing in the RefSeq annotation. (A) For *ZMYM2*, RefSeq identifies only the first two coding exons, resulting in a highly incomplete gene annotation. In contrast, TOGA2 using human, mouse, and elephant as references consistently identifies all downstream conserved coding exons. (B) For *RAB34*, RefSeq fuses the last four coding exons with their intervening introns, presumably due to the use of an incompletely spliced transcript, resulting in a truncated protein sequence. TOGA2 using all three references correctly identifies the individual exons, all having SpliceAI support. (C) Similarly, for *FBXL15*, RefSeq annotates a transcript that retains the second intron, resulting in a truncated protein, whereas TOGA2 using all three references correctly identifies the three coding exons.

**Supplementary Figure 28:**
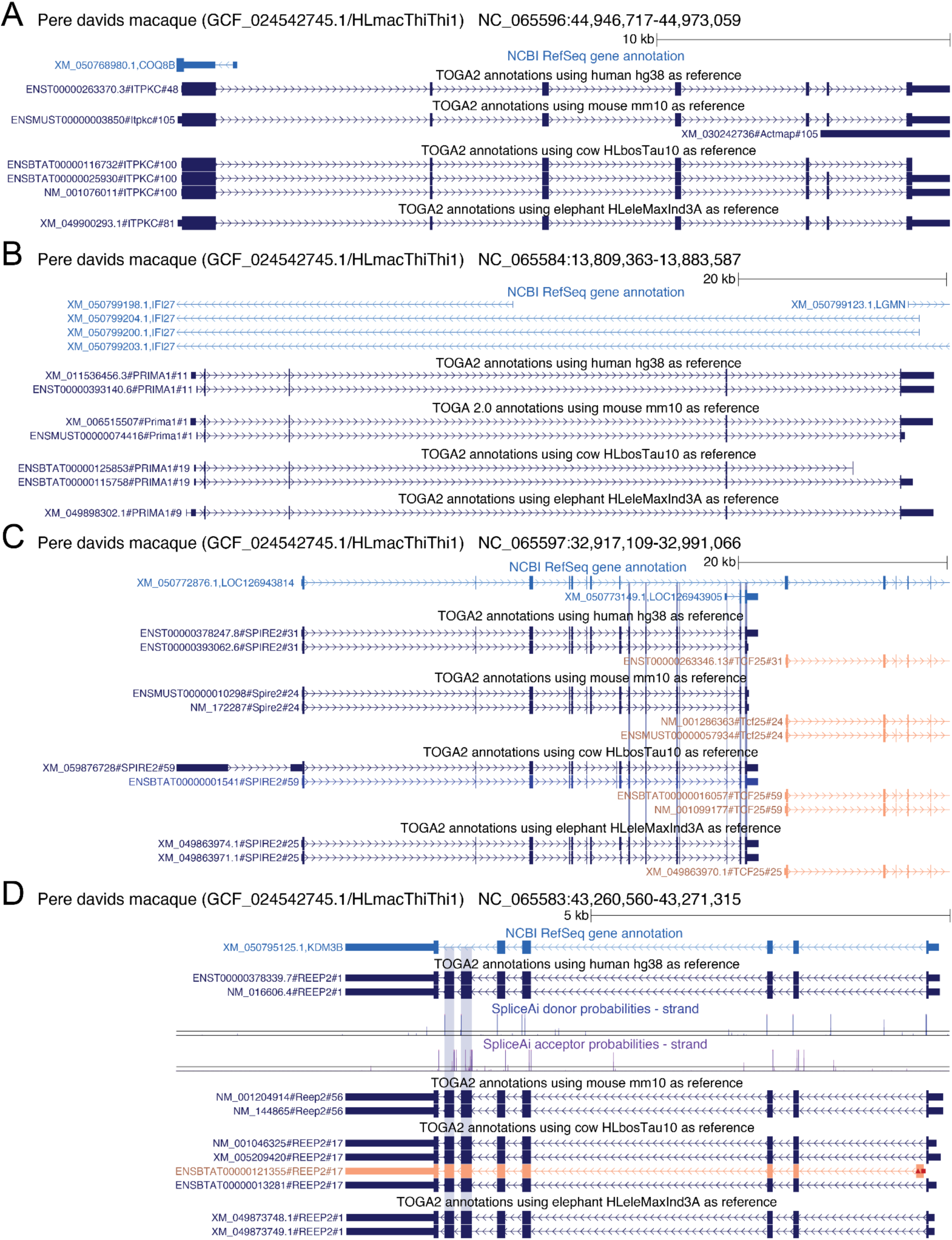
TOGA2 increases annotation completeness for Père David’s macaque. Examples of near-universally conserved genes present in the mammalia_odb12 gene set that are completely detected in TOGA2 annotations of the Père David’s macaque (also called Tibetan macaque) GCF_024542745.1 assembly, but are fragmented or missing in the RefSeq annotation. (A,B) *ITPKC* (A) and *PRIMA1* (B) are completely missing in the RefSeq annotation, but are correctly annotated by TOGA2 using human, mouse, cow, and elephant as references. (C) *SPIRE2* is misannotated by RefSeq, which fails to identify several conserved coding exons, annotates the last two coding exons as part of a separate gene (LOC126943905), and incorrectly fuses *SPIRE2* with exons from the neighboring *TCF25* gene. In contrast, TOGA2 using all four references correctly resolves both genes. *TCF25* is classified by TOGA2 as an uncertain loss, because the downstream exon 9 (not shown) contains a -1 frameshifting deletion, which may be an assembly base error. Instead of representing this frameshift, RefSeq creates a 2 bp intron to maintain the reading frame. (D) For *REEP2*, RefSeq fails to identify the sixth and seventh coding exons, whereas TOGA2 using all references correctly identifies both exons, which are supported by SpliceAI.

**Supplementary Figure 29:**
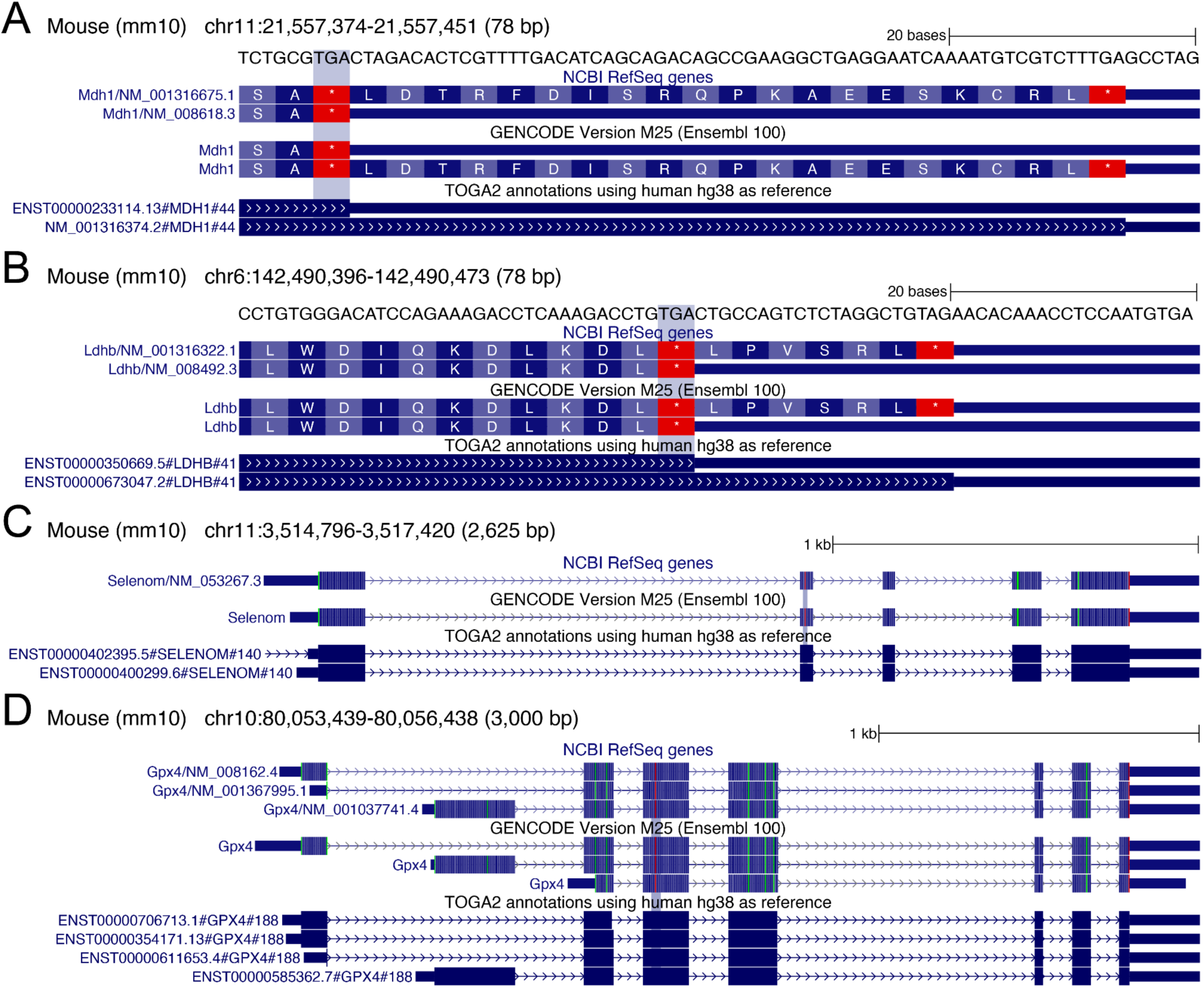
TOGA2 correctly annotates stop codon readthrough and selenocysteine-containing genes. UCSC Genome Browser screenshots of the mouse (mm10) genome showing RefSeq, GENCODE, and TOGA2 annotations for genes exhibiting stop codon readthrough (A, B) or containing selenocysteine-encoding TGA codons (C, D). TOGA2 correctly annotates the full coding sequence of these genes despite the presence of internal stop codons, because such codons are present in the input annotation and are masked during the CESAR2 alignment step. In contrast, other annotation methods typically terminate the coding sequence at such internal stop codons, resulting in truncated selenoprotein annotations or failure to capture readthrough-dependent coding transcript variation.

**Supplementary Figure 30:**
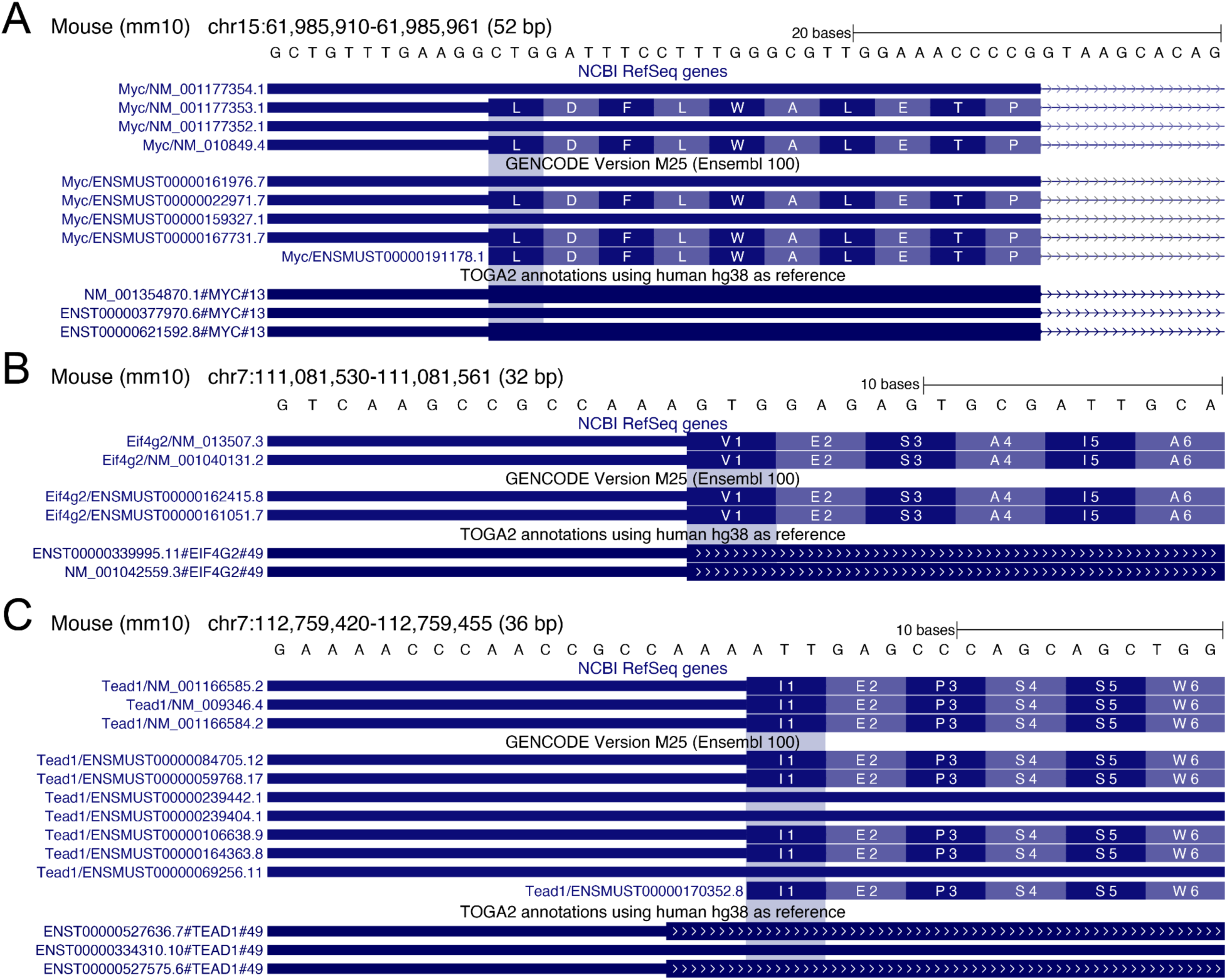
TOGA2 can annotate non-ATG start codons. While most gene annotation methods require a canonical ATG start codon, TOGA2 can annotate non-ATG start codons if they are present in the input annotation. (A) *MYC* contains a CTG start codon that is correctly annotated by TOGA2. (B) *EIF4G2* contains a GTG start codon that is correctly annotated by TOGA2. (C) *TEAD1* contains an ATT start codon. In this case, TOGA2 mispredicts the start site 3 bp upstream, illustrating that not all non-ATG start codons are captured correctly.

**Supplementary Figure 31:**
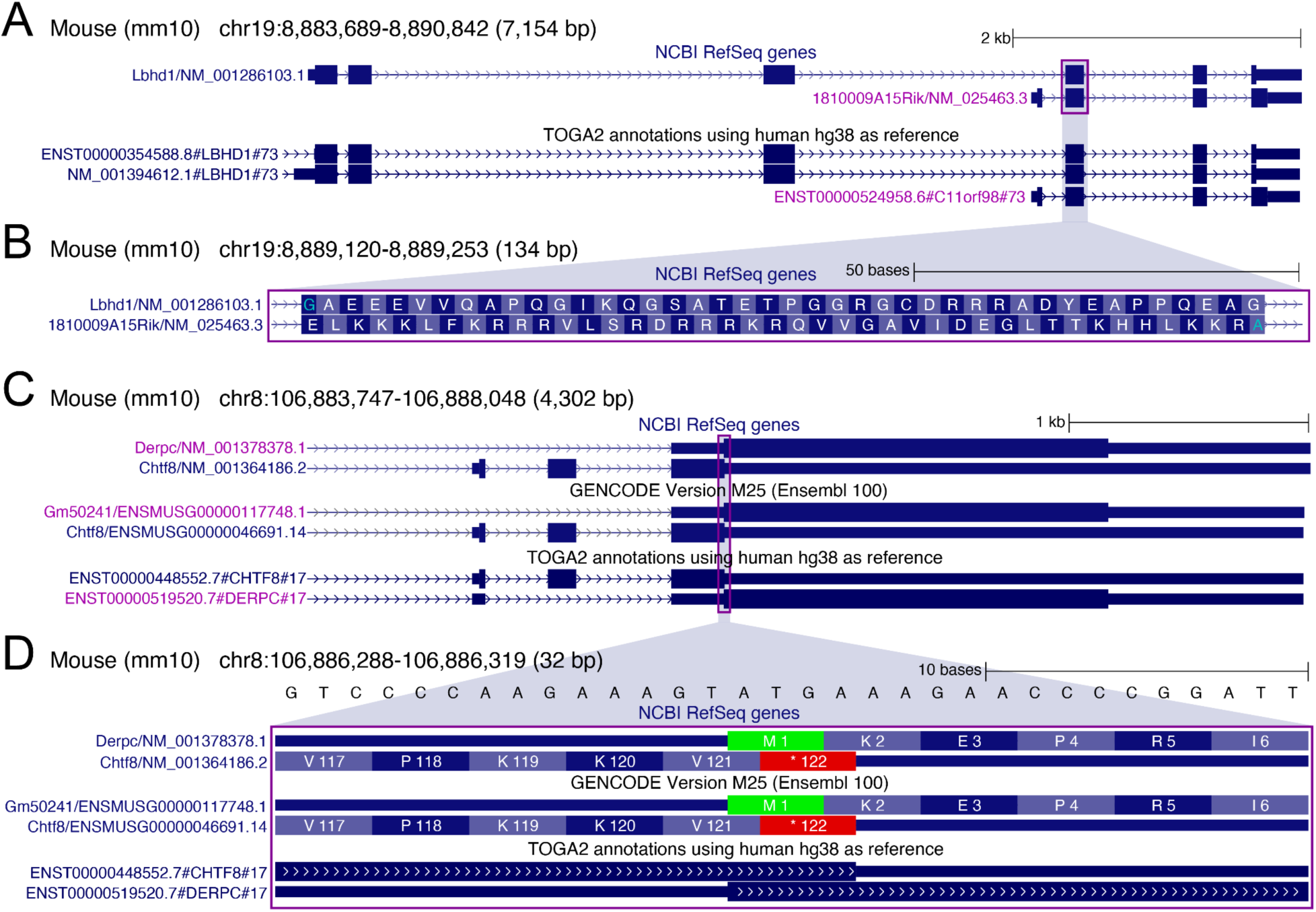
TOGA2 correctly annotates distinct genes with overlapping ORFs and infers 1:1 orthology. (A) *LBHD1* (blue font) and *C11orf98* (purple font; annotated as 1810009A15Rik in mouse) are annotated in human and mouse as two genes with distinct ORFs, although their transcripts share the last three exons. TOGA2 correctly annotates both genes. In contrast to TOGA1, which inferred many:one orthology, TOGA2 identifies both genes as 1:1 orthologs. Such cases are inherently challenging because TOGA infers genes in the query genome based on “same-strand, coding exon overlap”. When two distinct genes overlap in the query, as in this case, this strategy infers a single query gene. TOGA2 incorporates information from the reference genome, where the overlap between the two genes is already present, thereby enabling correct 1:1 orthology assignment. (B) Zoom-in of the third last exon (purple box in A), showing that this exon encodes two ORFs in different reading frames. (C) *CHTF8* (blue font) and *DERPC* (purple font) have distinct ORFs, but share the same last exon. TOGA2 correctly annotates both genes and, by incorporating information that the genes overlap in the reference genome, identifies them as 1:1 orthologs. (D) Zoom-in showing that the end of the *CHTF8* ORF overlaps the beginning of the *DERPC* ORF.

**Supplementary Figure 32:**
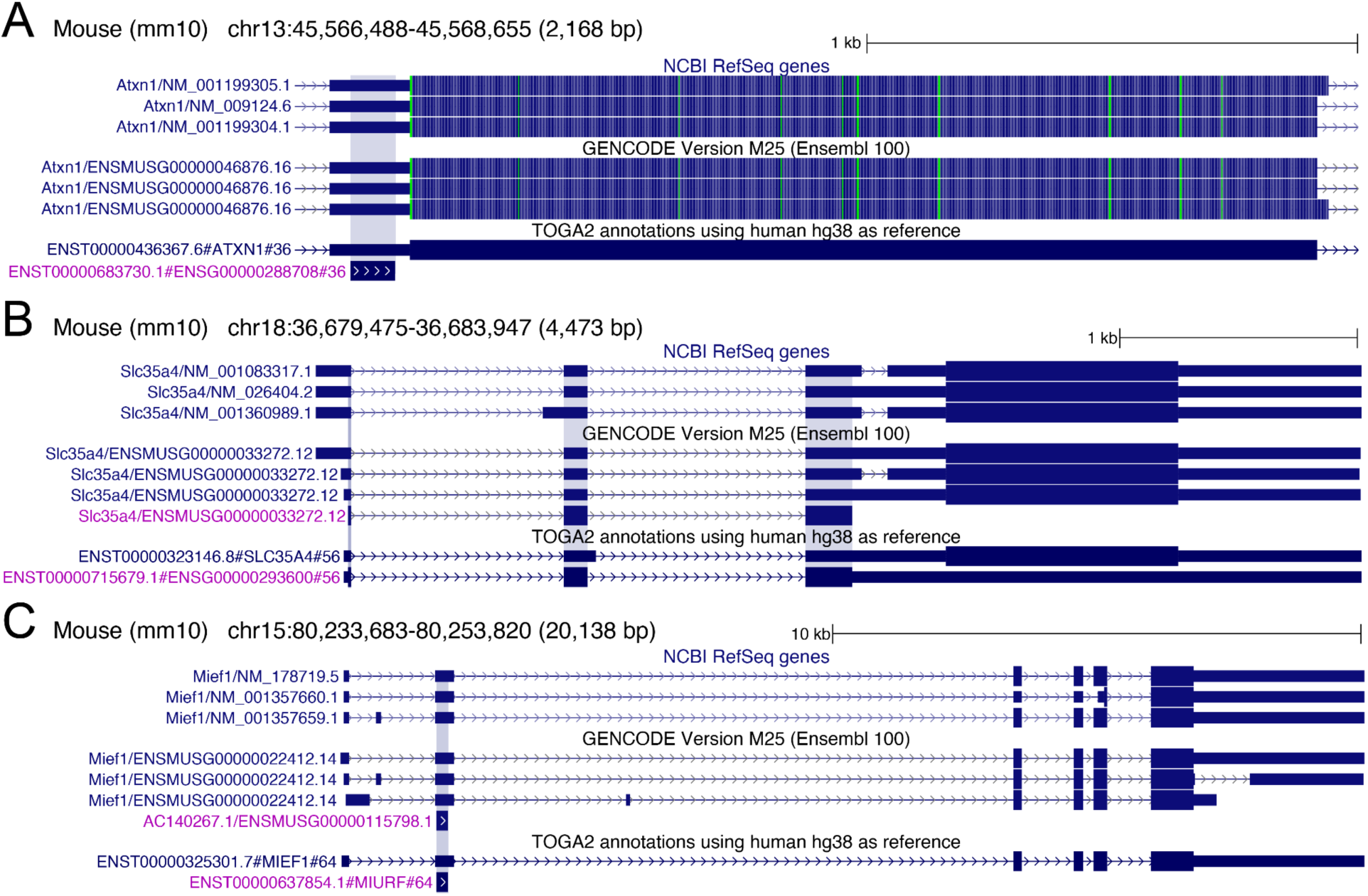
TOGA2 correctly annotates genes with dual ORFs. (A) *ATXN1* encodes a conserved upstream 29 amino acid alternative ORF (Alt-ATXN1) that is annotated only by TOGA2. Alt-ATXN1 is a nuclear RNA-binding protein that directly interacts with ATXN1 ^112^. (B) *SLC35A4* encodes an upstream 103 amino acid alternative ORF (AltSLC35A4) annotated by GENCODE and TOGA2, but not by RefSeq. AltSLC35A4 is an inner mitochondrial membrane protein that contributes to protection against oxidative stress ^113^. (C) *MIEF1* encodes an upstream 70 amino acid alternative ORF (MIURF) annotated by GENCODE and TOGA2, but not by RefSeq. MIURF is a mitochondrial translation regulator that binds to the mitoribosome ^114^. The transcript encoding the alternative ORF is shown in purple font, and its coding region is highlighted with a blue background.

**Supplementary Figure 33:**
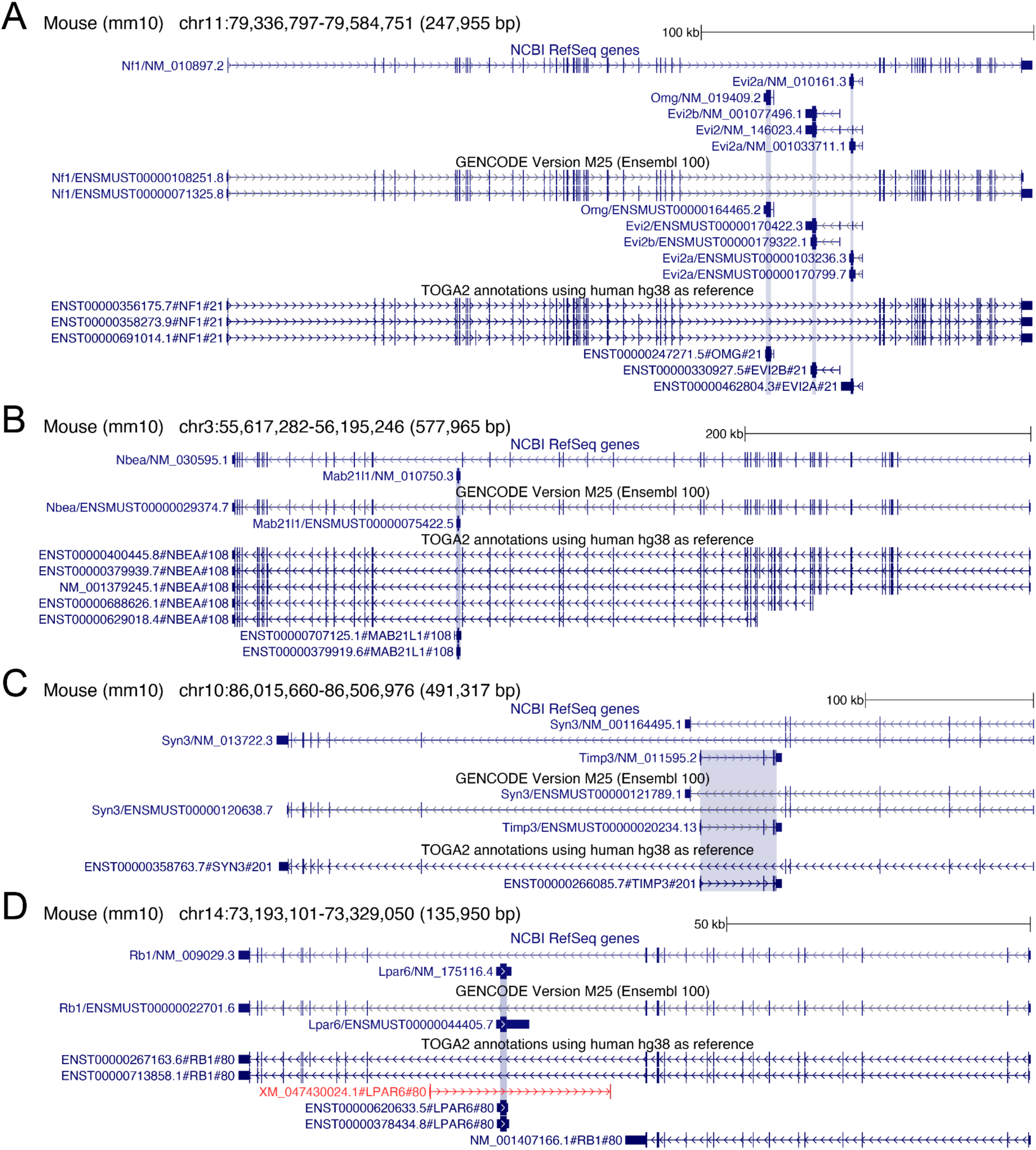
TOGA2 accurately annotates nested genes. Several examples illustrating TOGA2’s ability to annotate genes located within introns of other host genes. (A) *OMG*, *EVI2B*, and *EVI2A* are nested within a large intron of the host gene *NF1*. (B) *MAB21L1* is nested within the host gene *NBEA*. (C) *TIMP3* is nested within *SYN3*. (D) *LPAR6* is nested within *RB1*.

**Supplementary Figure 34:**
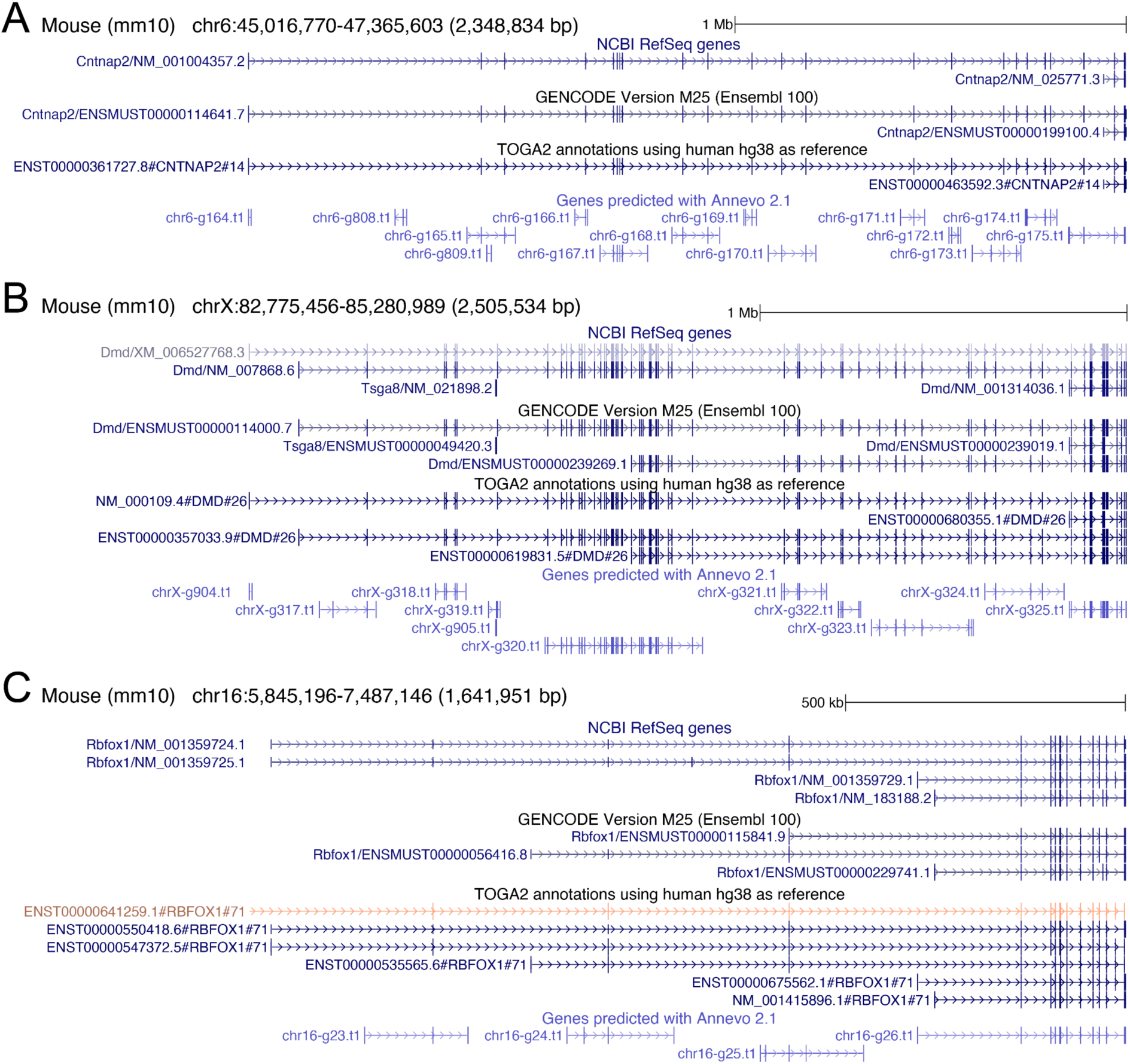
TOGA2 accurately annotates genes with long introns. While the median intron length in human is ∼1.5 kb, some genes contain exceptionally large introns and span large genomic regions, which poses challenges for gene prediction. Despite having long introns, TOGA2 correctly annotates such genes. In contrast, the *ab initio* gene predictor Annevo ^28^ frequently predicts these genes as multiple fragments. We note, however, that *ab initio* gene prediction is substantially more challenging than homology-based gene annotation. (A) *RBFOX1* contains several introns exceeding 300 kb in length, and the gene locus spans more than 1.5 Mb. (B) *DMD* contains multiple introns exceeding 200 kb, and the entire gene locus spans > 2.3 Mb. (C) The first intron of *CNTNAP2* is 593,933 bp long, and the gene locus spans > 2.2 Mb.

**Supplementary Figure 35:**
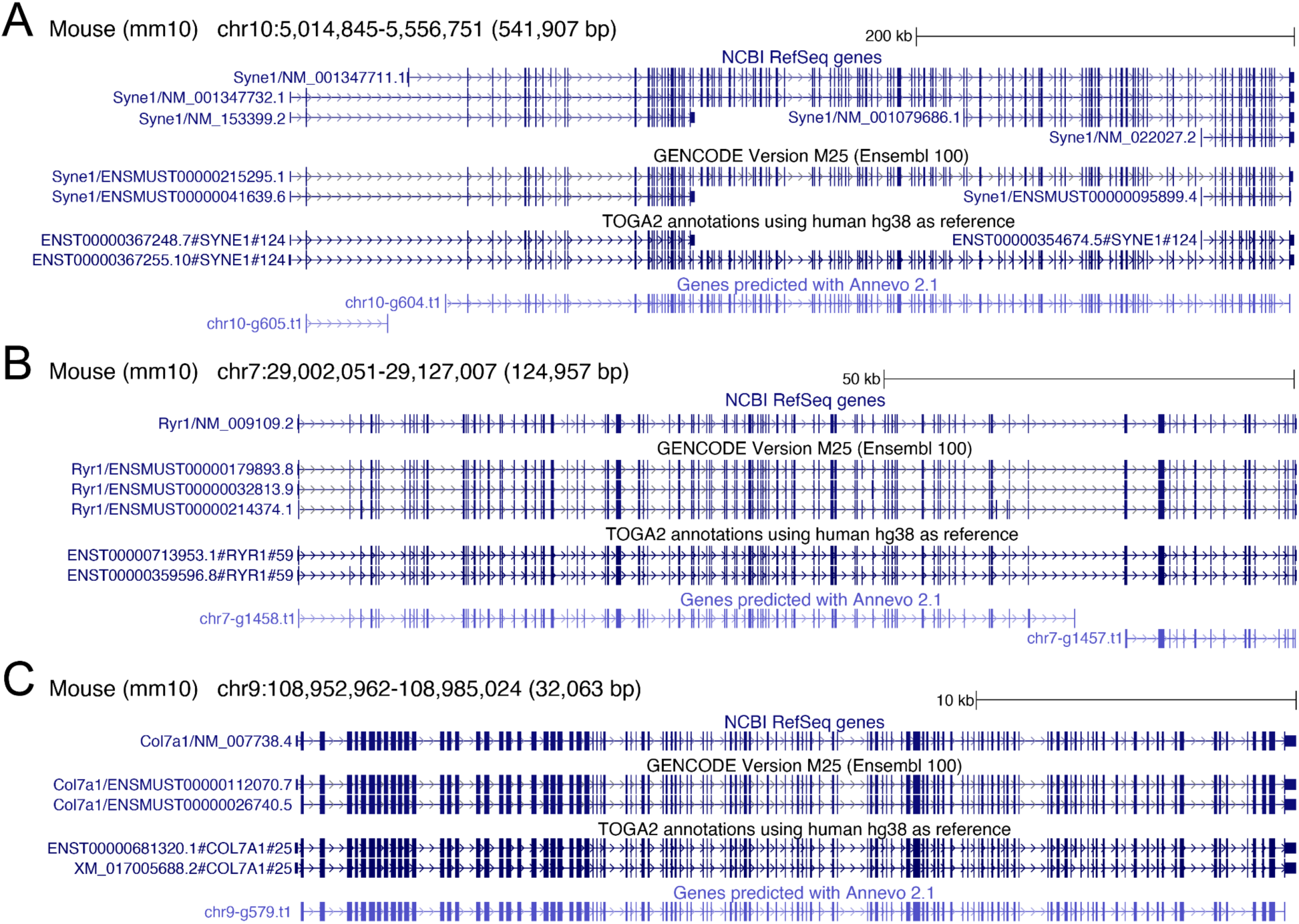
TOGA2 accurately annotates genes with many exons. While the median number of exons per gene in human is 8, some genes contain exceptionally high exon counts. Such gene structures can pose challenges for *ab initio* gene predictors in some cases; however, TOGA2 correctly annotates such genes. As noted above, we emphasize that *ab initio* gene prediction is substantially more challenging than homology-based annotation. (A) *SYNE1* has isoforms comprising up to 146 exons. (B) *RYR1* has isoforms encoding up to 106 exons. (C) *COL7A1* has isoforms encoding 119 exons.

**Supplementary Figure 36:**
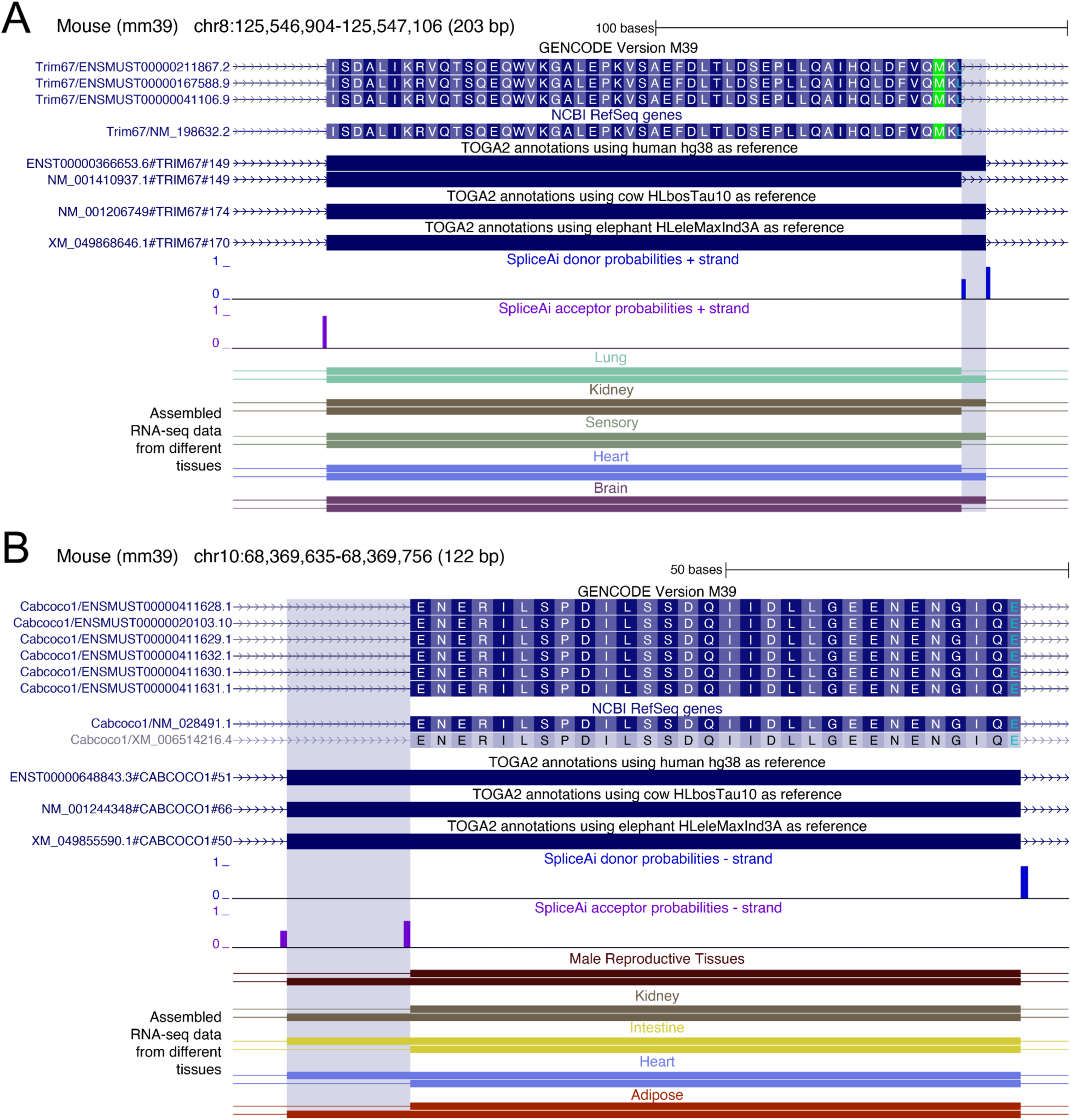
TOGA2 identifies alternative splice sites not represented in the mouse GENCODE VM39 annotation. (A) Using all three mammalian reference species, TOGA2 predicts an alternative donor splice site in the *TRIM67* gene that extends the exon by 6 bp (blue highlight). (B) TOGA2 predicts an alternative acceptor splice site in *CABCOCO1* that extends the exon by 18 bp (blue highlight). This longer exon variant is supported by all three reference species. Both alternative splice sites are not represented in the GENCODE or RefSeq annotations, but RNA-seq transcripts from multiple tissues support their usage.

**Supplementary Figure 37:**
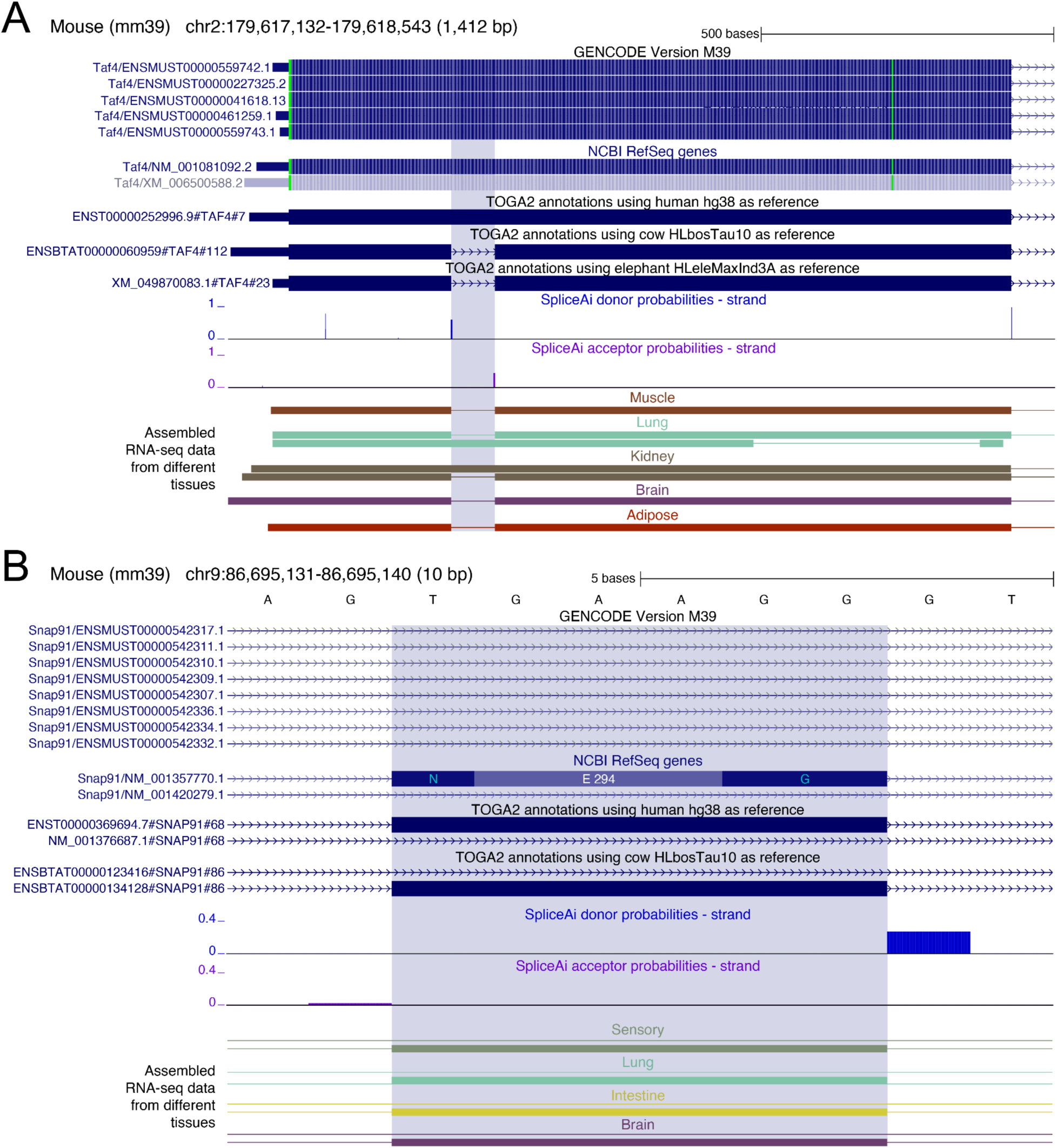
TOGA2 identifies novel alternative introns and exons not represented in the mouse GENCODE VM39 annotation. (A) Using cow and elephant as reference species, TOGA2 predicts a 75 bp intron within the first exon of *TAF4*. This intron is translatable, meaning its inclusion preserves the reading frame. This intron is not represented in the GENCODE or RefSeq annotations; however, RNA-seq transcripts from multiple tissues indicate that most transcripts remove this intron. (B) Using human and cow as reference species, TOGA2 predicts a 6 bp alternative exon in *SNAP91*. This exon is supported by RefSeq and RNA-seq data from multiple tissues but is not represented in the GENCODE annotation. For visual clarity, only a subset of the transcripts annotated by GENCODE, RefSeq, and TOGA2 is shown.

**Supplementary Figure 38:**
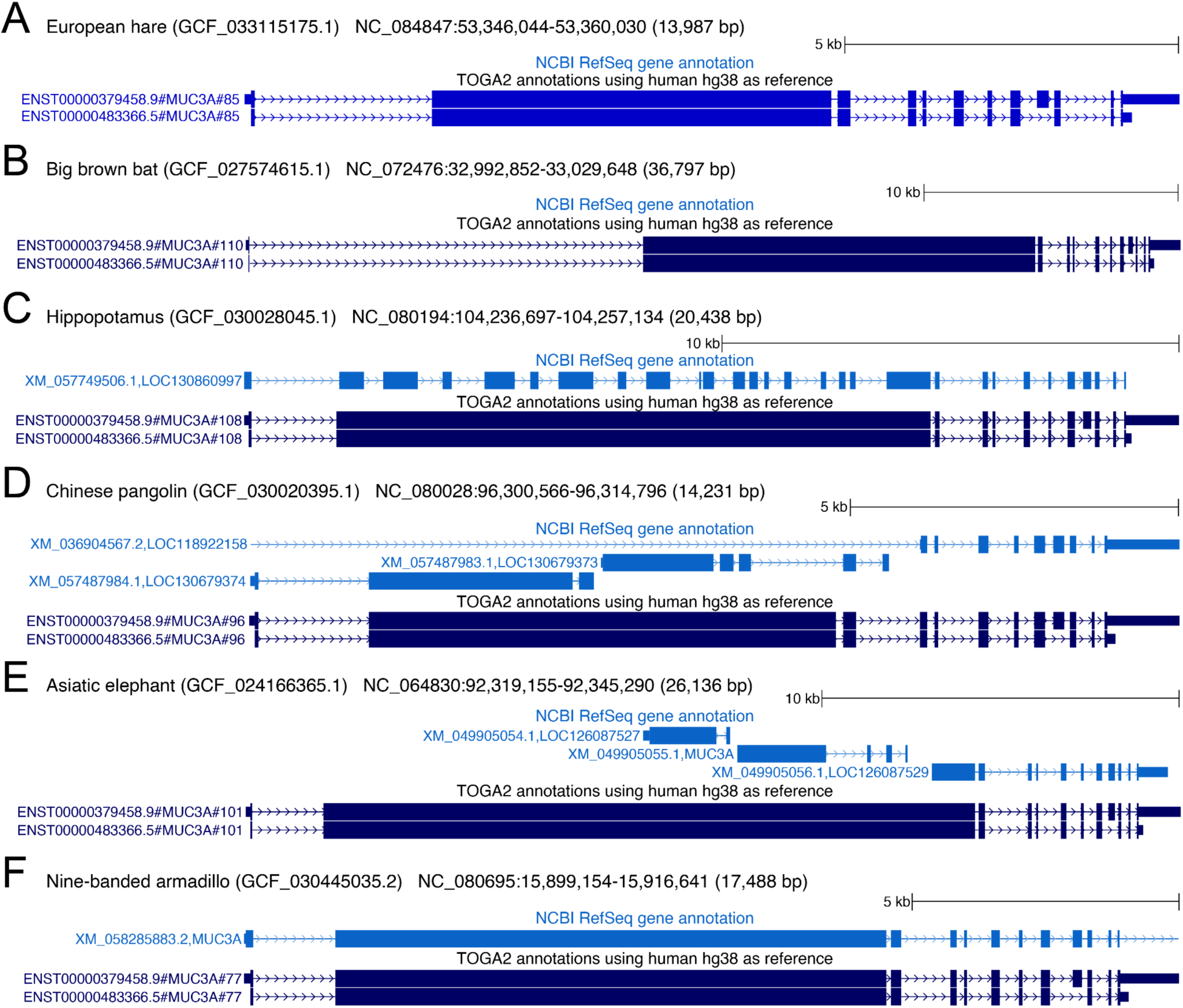
TOGA2 annotates the long-exon-containing *MUC3A* transcript across taxonomically diverse placental mammals. (A–F) TOGA2 identifies *MUC3A* transcripts containing the large second coding exon in species representing diverse placental mammal orders, including hare (A), bat (B), hippopotamus (C), pangolin (D), elephant (E), and armadillo (F). Across these species, NCBI RefSeq either does not annotate *MUC3A* (A,B) or annotates different, often fragmented exon-intron structures (C–E). In armadillo, RefSeq incorrectly fuses *MUC3A* with the downstream *MUC12* gene (not shown in panel F). In hippopotamus and armadillo, RefSeq additionally annotates translation start codons that are located 130 bp (hippopotamus) and 75 bp (armadillo) upstream of the conserved ATG start codon identified by TOGA2.

**Supplementary Figure 39:**
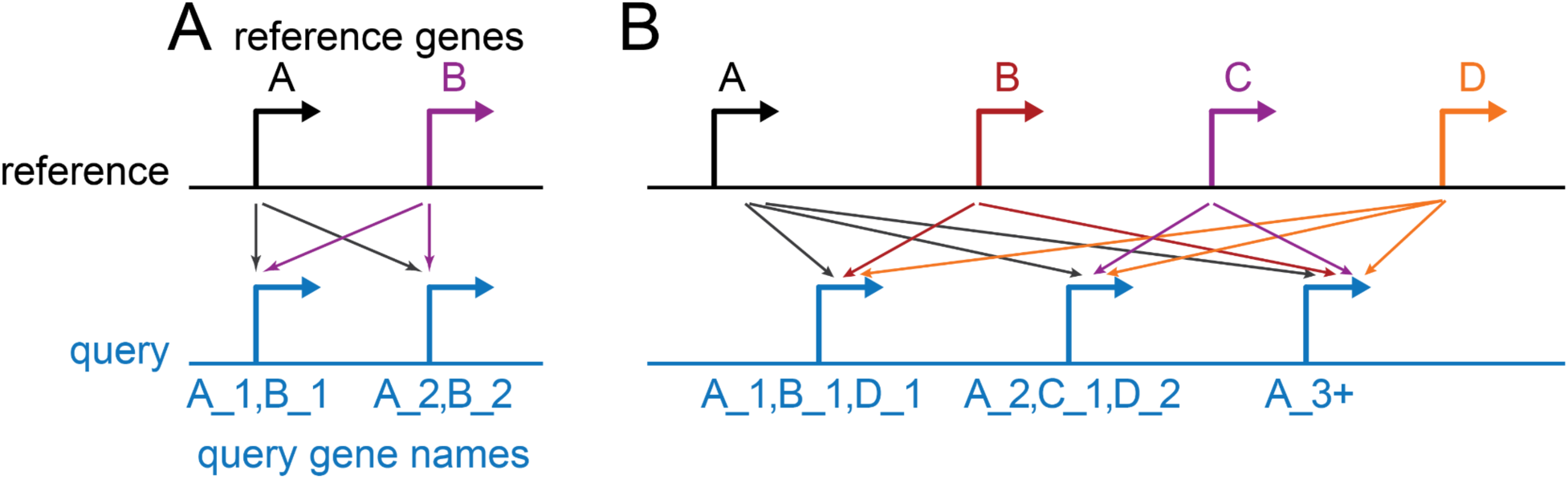
Illustration of gene names assigned to many:many orthologs. (A) Both query genes are orthologous to reference genes *A* and *B*. To avoid assigning identical names, TOGA2 distinguishes them by adding the suffixes _1 and _2. (B) The three query genes, distinguished by _{num} suffixes, have orthology relationships to three or all four reference genes. Their names reflect the corresponding comma-separated lists of orthologous reference genes, except for the third query gene, where TOGA2 selects a representative reference ortholog and appends a “+” symbol.

**Supplementary Figure 40:**
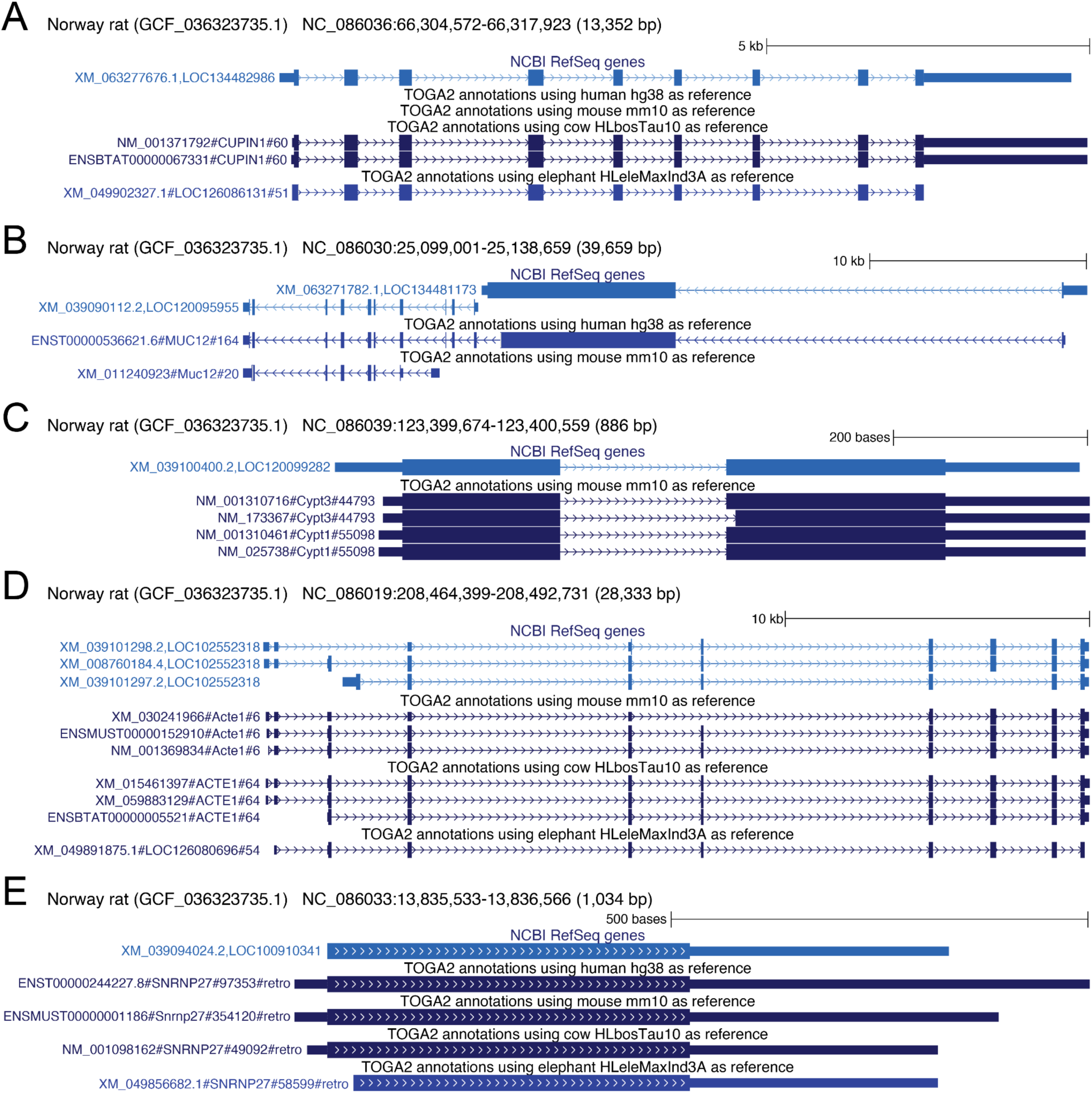
TOGA2 assigns informative gene symbols to many RefSeq LOC genes. We selected the model organism Norwegian rat to illustrate representative examples of RefSeq locus (LOC) gene identifiers that match TOGA2-annotated orthologs with informative gene symbols. We note that many other analyzed species contain higher numbers of RefSeq LOC genes. (A) *LOC134482986* matches the cow ortholog of *CUPIN1*. While *CUPIN1* is lost in both human and mouse and is therefore absent from the input annotation for these references, the elephant ortholog is also annotated with a LOC gene symbol. (B) *LOC120095955* and *LOC134481173* likely represent fragmented annotations that both match the human *MUC12* ortholog. *LOC120095955* also corresponds to a likely fragmented mouse *Muc12* ortholog. (C) Transcripts of *LOC120099282* match mouse orthologs of *Cypt1* and *Cypt3*. No other TOGA2 reference annotates an orthologous gene at this locus in rat. (D) Transcripts of *LOC102552318* match TOGA2-annotated orthologs of *ACTE1*. In elephant, the *ACTE1* ortholog is also annotated with a LOC gene symbol. In human, *ACTE1* is lost and therefore absent from the TOGA2 input annotation. (E) *LOC100910341* represents a retrogene of *SNRNP27*.

**Supplementary Figure 41:**
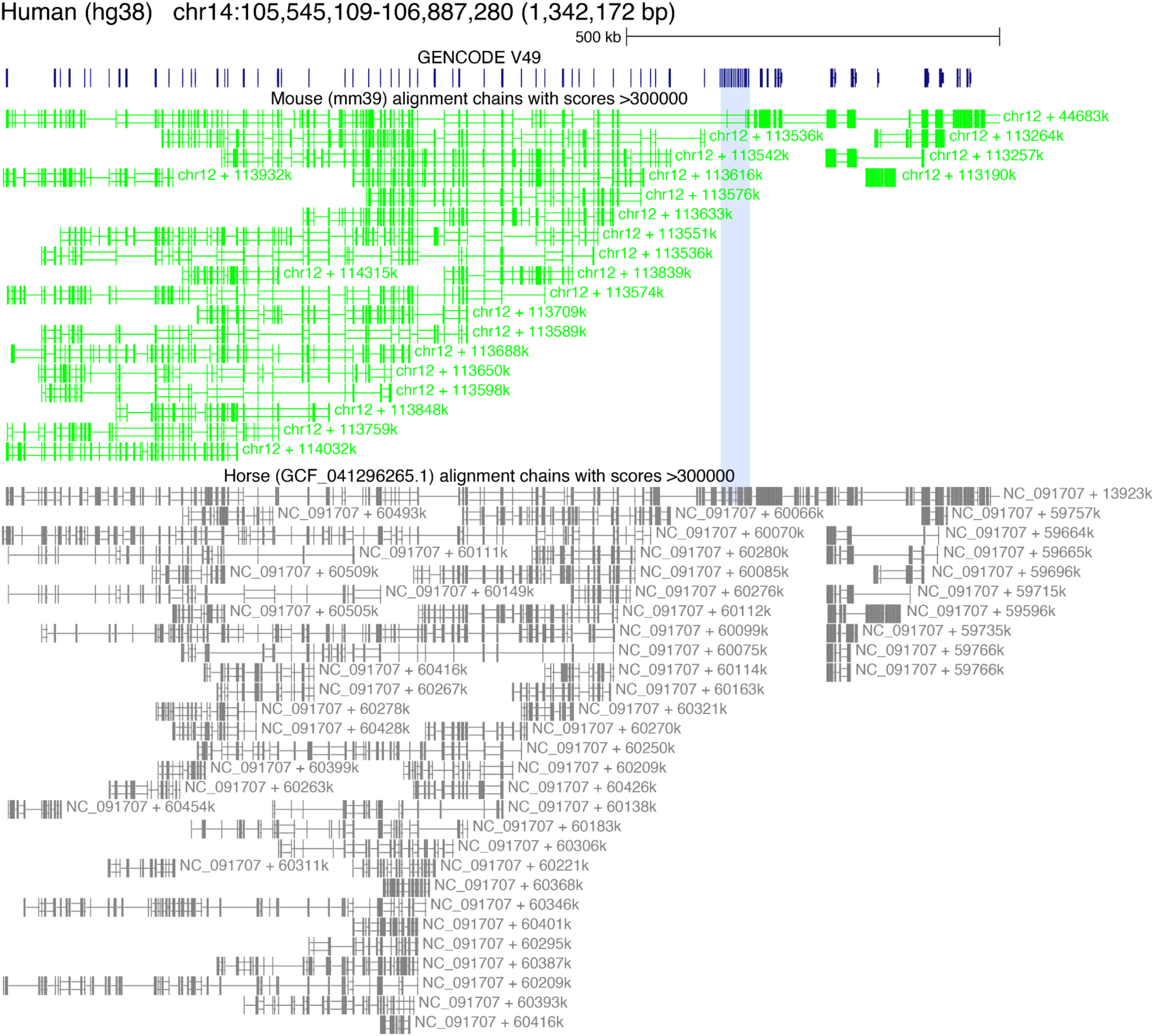
V(D)J segments are captured by sensitive whole-genome alignment chains across placental mammals. Human genome browser screenshot showing the IGH locus and alignment chains to mouse and horse. Many V(D)J segments align across these placental mammals, especially in horse, which exhibits lower sequence divergence from human (substitutions per site) than mouse. The D-segment locus (blue highlight) contains substantially fewer alignments. Because V(D)J segments are highly similar to one another, many overlapping alignment chains exist. For visual clarity, only chains with scores greater than 300,000 are shown.

**Supplementary Figure 42:**
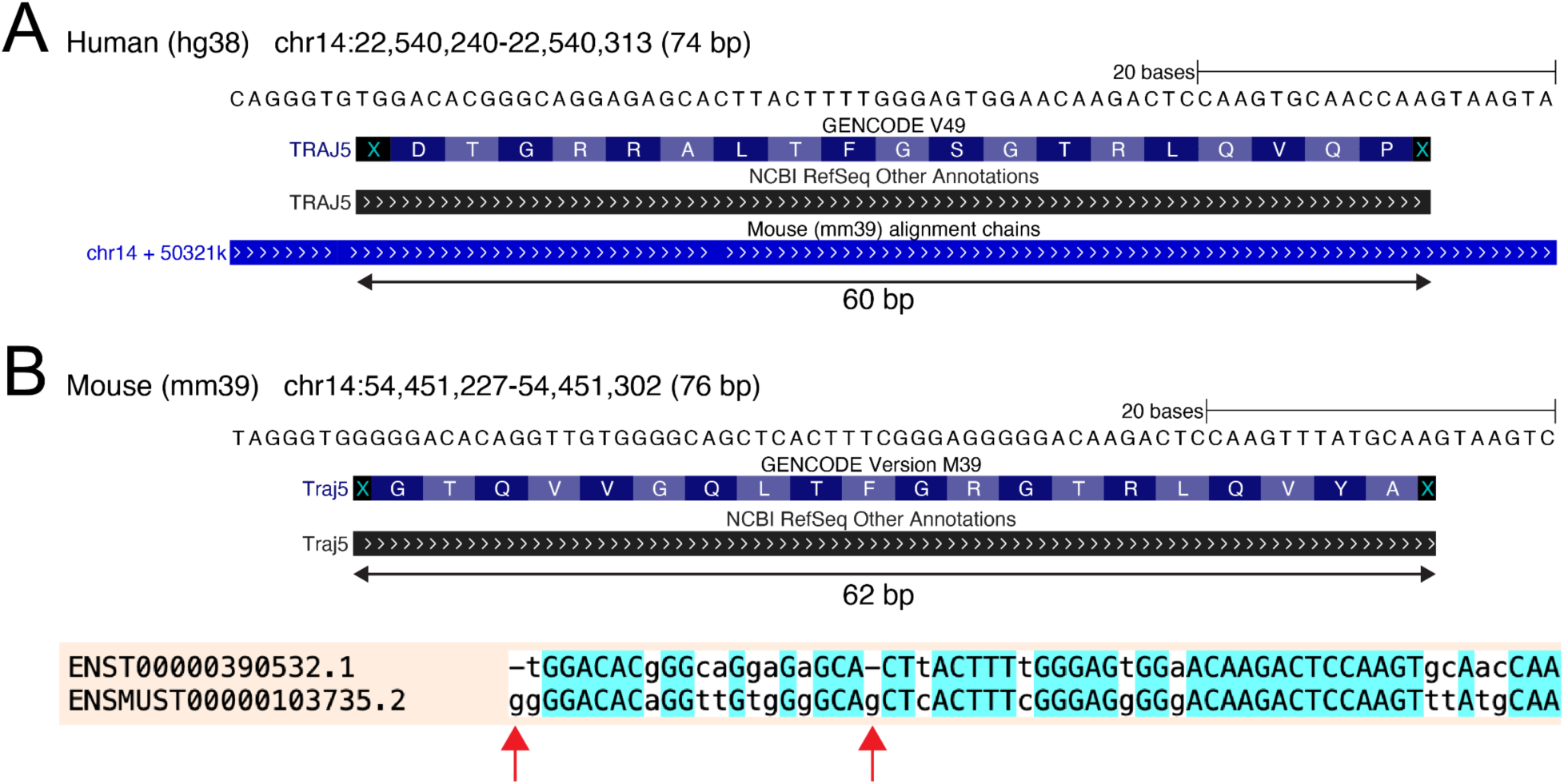
Immunoglobulin and T-cell receptor segments can contain frameshifting mutations. (A) Genome browser screenshot showing the human *TRAJ5* locus (T-cell receptor alpha locus, J segment 5), which aligns to a single orthologous locus in mouse. (B) The orthologous *TRAJ5* locus in mouse mm39. Whereas the human *TRAJ5* segment is 60 bp long, the mouse segment is 62 bp long due to two frameshifting mutations (red arrows). Because non-templated nucleotides are added during V(D)J recombination, such frameshifts can still give rise to functional T-cell receptors. For this reason, TOGA2 considers only internal stop codons when assessing reading-frame integrity.

**Supplementary Figure 43:**
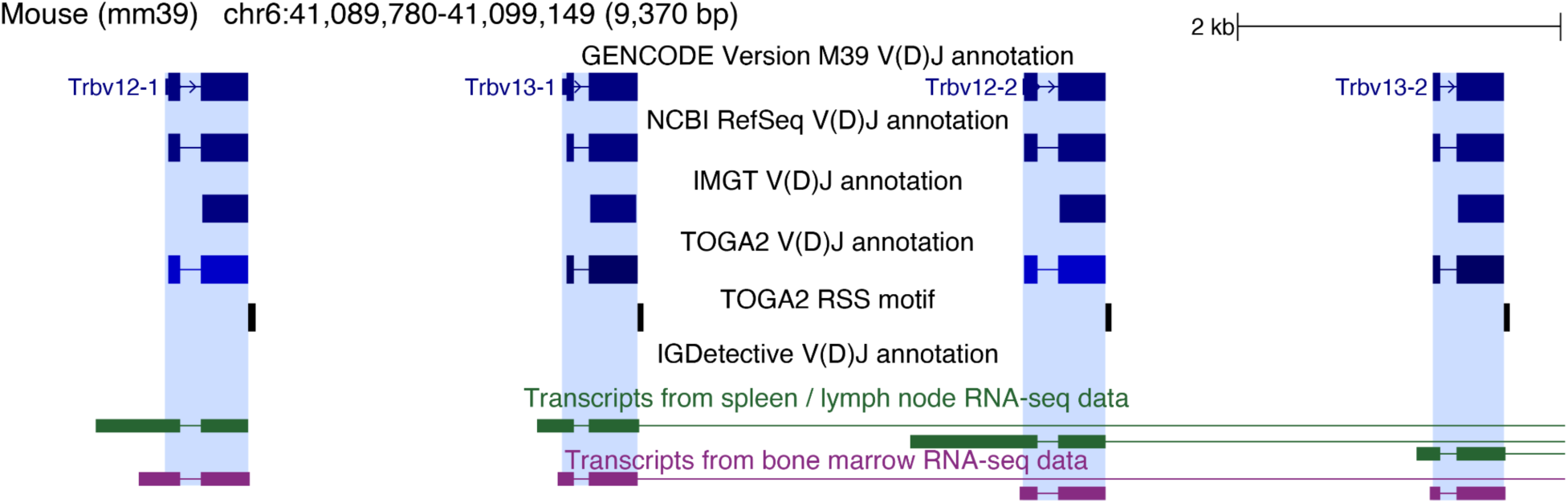
T-cell receptor V segments identified by TOGA2 but not by IGDetective. Genome browser screenshot showing several V segments in the T-cell receptor beta locus that are identified by TOGA2 but are not reported by IGDetective. These segments, including their leader exons, are supported by RNA-seq data.

**Supplementary Figure 44:**
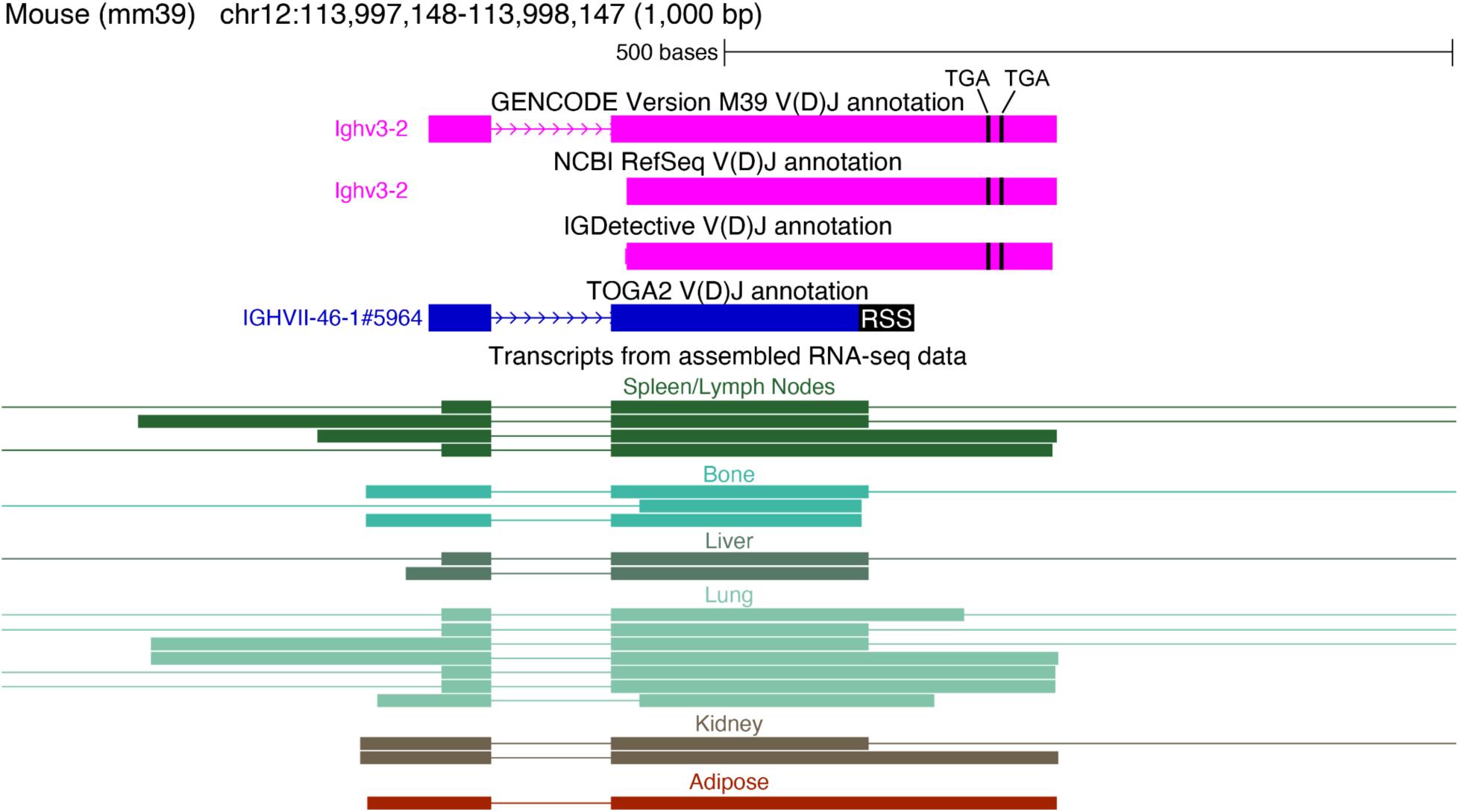
Example of a putative alternative, transcript-supported RSS. GENCODE, RefSeq, and IGDetective classify this *IGH* V segment as non-functional because the open reading frame is interrupted by two TGA stop codons (black lines). In contrast, TOGA2 identifies an RSS further upstream (black), placing the two TGA codons downstream of the predicted V segment. RNA-seq transcripts from multiple tissues support the usage of both the annotated RSS and the upstream RSS identified by TOGA2.

**Supplementary Figure 45:**
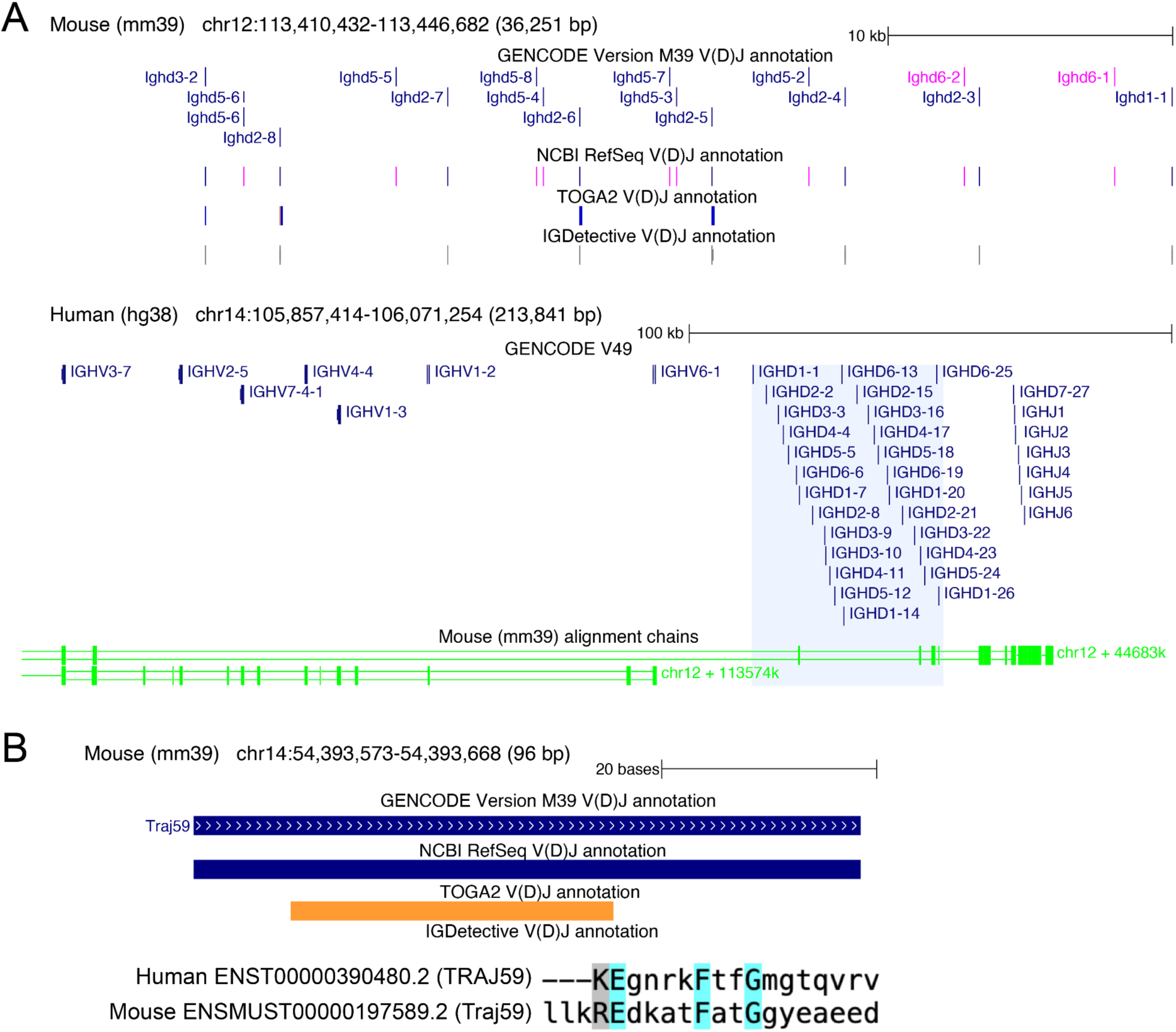
Examples illustrating limitations of TOGA2 in annotating D and J segments. (A) The top panel shows D segments in the mouse *IGH* locus. Compared with IGDetective, TOGA2 fails to correctly annotate several segments classified as functional by GENCODE. The primary reason is that the short length and high sequence divergence of D segments limit whole-genome alignments. Despite using sensitive genome alignment parameters, only three D segments are captured by the orthologous human-mouse alignment chain, as shown in the bottom panel. (B) Example of a J segment that is incorrectly annotated by TOGA2. The orange color indicates that TOGA2 could not identify a recombination signal sequence (RSS). The sequence alignment (bottom) shows that the orthologous human and mouse segments are highly diverged, sharing only three identical amino acids. As a result, TOGA2’s CESAR2-based alignment identifies a segment candidate that is too short, preventing correct identification of the adjacent RSS.

**Supplementary Figure 46:**
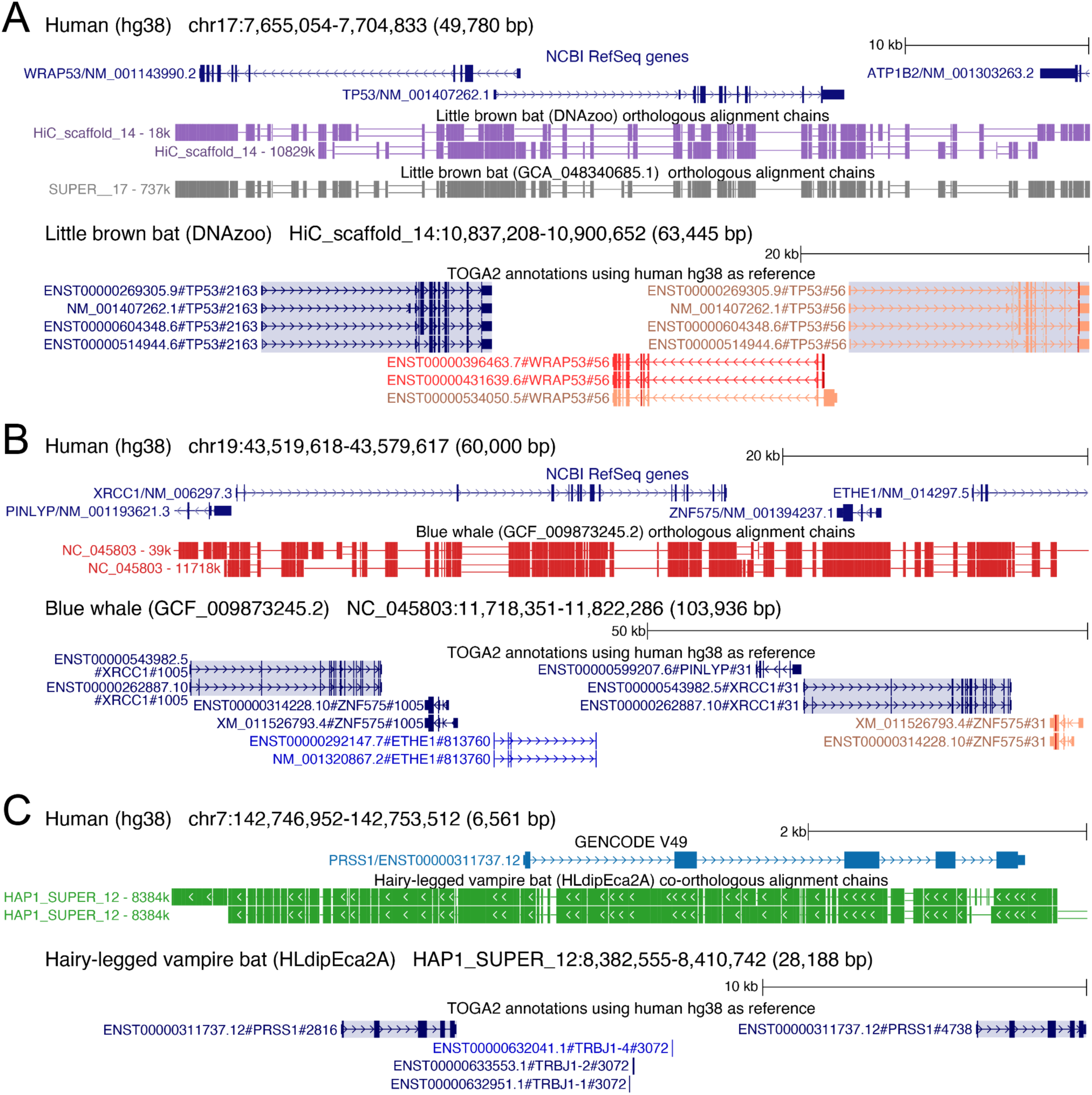
Examples of gene duplications detected by TOGA2. (A) Top panel: TOGA2 detects two orthologous loci corresponding to the *TP53* gene in the DNAZoo assembly of the little brown bat ^115^, which is based on the Zoonomia Illumina-based GCF_000147115.1 assembly ^116^, indicating a *TP53* duplication previously noted in ^117^. Bottom panel: TOGA2 infers a 1:many orthology relationship for *TP53* in the little brown bat and annotates both gene copies (blue highlight). The downstream copy contains a -1 frameshifting deletion, leading TOGA2 to classify it as uncertain loss. However, a high-quality HiFi-based assembly of the little brown bat (GCA_048340685.1) contains only a single orthologous *TP53* locus ^55^, indicating that the putative duplication in the DNAZoo assembly either represents a polymorphic event not fixed in the species or an assembly artifact caused by incorporation of both haplotypes into the assembly. (B) Top panel: TOGA2 detects a previously described duplication of *XRCC1* in the blue whale ^118^, indicated by two co-orthologous alignment chains covering *XRCC1* and the neighboring *ZNF575* locus. Bottom panel: TOGA2 infers 1:many orthology relationships for *XRCC1* and *ZNF575* and annotates both gene copies (*XRCC1* is highlighted). The second *ZNF575* copy contains a -1 frameshifting deletion and is therefore classified as uncertain loss. (C) TOGA2 identified the duplication of the pancreatic trypsin-encoding gene *PRSS1* in the Hairy-legged vampire bat genome ^119^, visible as two co-orthologous alignment chains (top), and annotates two fully intact gene copies in this bat species (bottom).

**Supplementary Figure 47:**
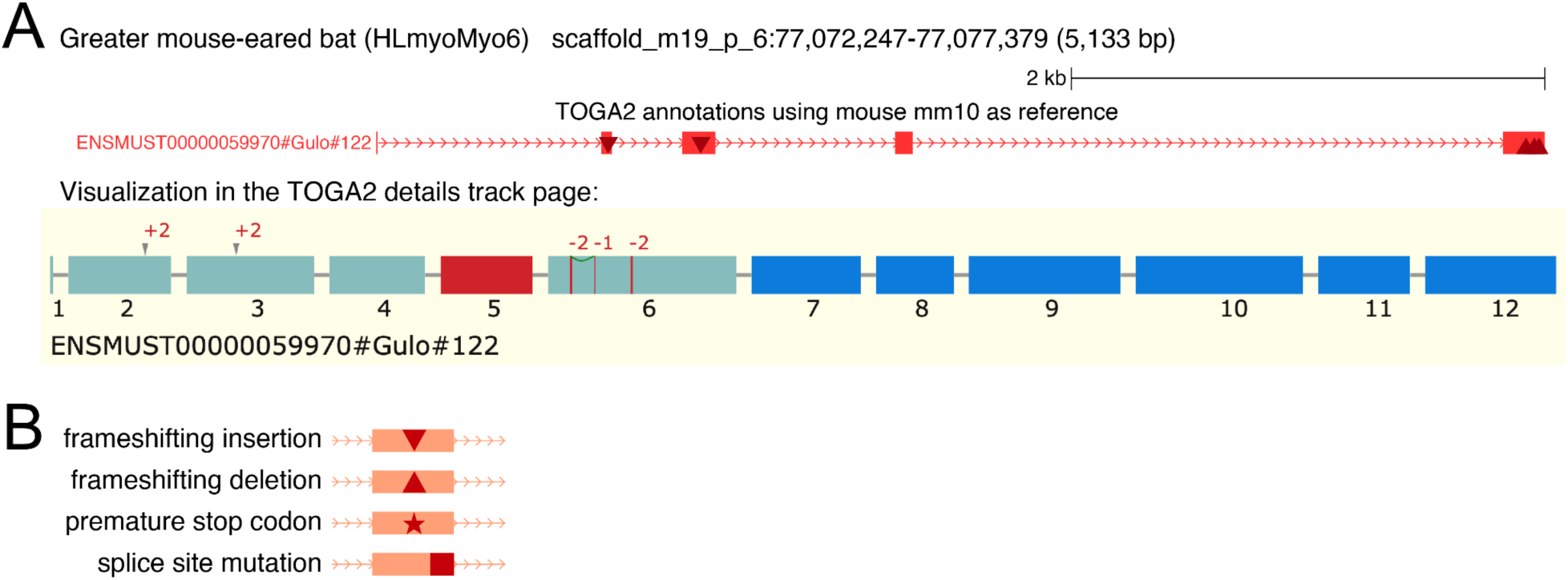
Annotation of lost genes and visualization of inactivating mutations with UCSC decorator tracks. (A) Top: UCSC Genome Browser screenshot showing the *Gulo* gene, which is lost in *Myotis* bats ^120^. Triangles indicate frameshifting insertions and deletions. Bottom: Visualization in the TOGA2 details page. Because exons 5 and 7-12 are deleted in the *Myotis myotis* genome ^44^, the TOGA2 annotation browser track (top) visualizes only the five exons that remain present in the genome. (B) UCSC decorator symbols used to visualize different types of inactivating mutations.

**Supplementary Figure 48:**
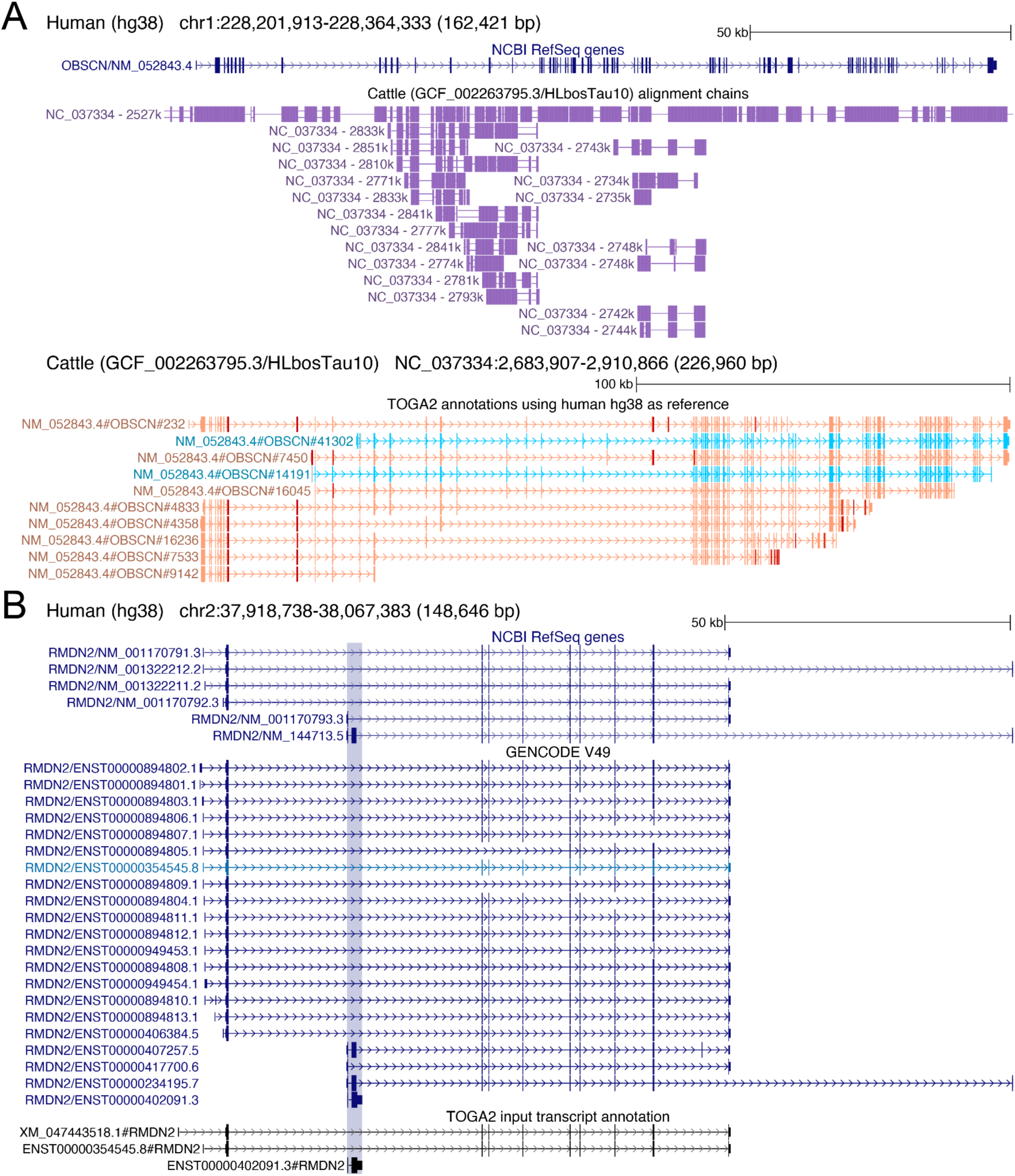
TOGA2 handles rare situations that complicate query gene inference. A) The muscle gene *OBSCN* is highly repetitive (top panel). As illustrated for cow as the query species, this results in many alignment chains mapping different parts of the same gene to the same query locus. For simplicity, only a single *OBSCN* transcript is shown. Because these chains contain intronic alignments, TOGA2 correctly recognizes them as orthologous, despite covering only a portion of the gene. The bottom panel illustrates that search space extension then results in the same transcript being annotated multiple times through different alignment chains in the same cow locus, but exons directly supported by the alignment chains do not necessarily overlap. Because TOGA2 normally assesses same-strand coding exon overlap only for exons supported by the alignment chain, this situation could incorrectly lead to inference of multiple query genes. TOGA2 addresses this by collapsing projections originating from the same transcript within the same query locus into a single inferred query gene. (B) *RMDN2* has multiple isoforms differing by exon skipping events, alternative promoters, and a distal downstream exon. Of these, TOGA2’s evolutionarily informed transcript selection retains three transcripts, including a short two-exon transcript that shares no overlapping coding exons with the other two transcripts. Consequently, projections of the short transcript would not overlap coding exons of the other transcripts in the query genome and would therefore be incorrectly assigned to a separate query gene. TOGA2 addresses this by collapsing projections from the same reference gene into a single inferred query gene if they are located within the same query locus span defined by projections from other transcripts of the same gene.

**Supplementary Figure 49:**
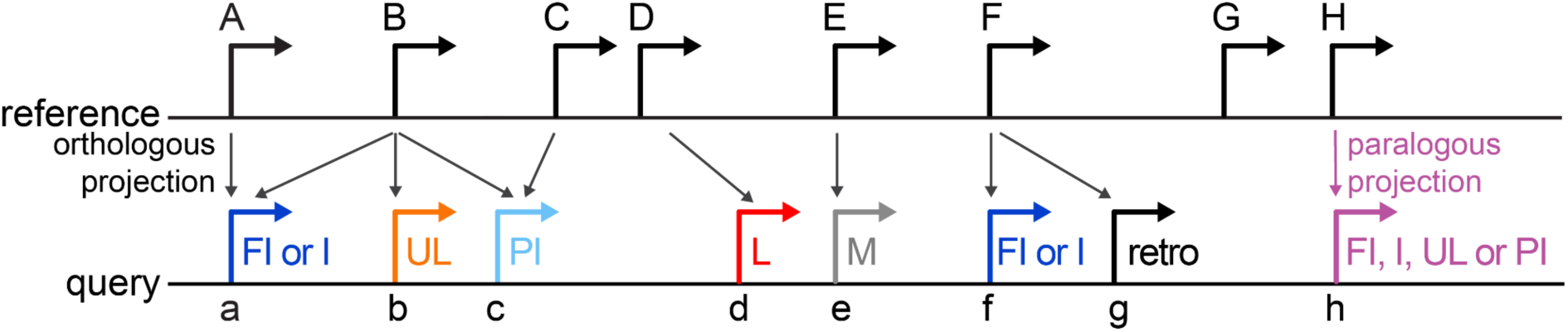
Illustration of TOGA2’s run summary and query gene counting procedure. Query genes inferred from orthologous projections with loss status FI, I, PI, or UL are counted, together with genes inferred solely from paralogous projections with loss status FI, I, PI, or UL. Retrogene candidates are reported separately. Lost and Missing genes are summarized as separate categories.

